# Potential hydrodynamic cytoplasmic transfer between mammalian cells: Cell-projection pumping

**DOI:** 10.1101/531798

**Authors:** Hans Zoellner, Navid Paknejad, James Cornwell, Belal Chami, Yevgeniy Romin, Vitaly Boyko, Sho Fujisawa, Elizabeth Kelly, Garry W. Lynch, Glynn Rogers, Katia Manova, Malcolm A.S. Moore

## Abstract

We earlier reported cytoplasmic fluorescence exchange between cultured human fibroblasts (Fib) and malignant cells (MC). Others report similar transfer via either tunneling nanotubes (TNT) or shed membrane vesicles and this changes the phenotype of recipient cells. Our current time-lapse microscopy showed most exchange was from Fib into MC, with less in the reverse direction. Although TNT were seen, we were surprised transfer was not via TNT, but was instead via fine and often branching cell projections that defied direct visual resolution because of their size and rapid movement. Their structure was revealed nonetheless, by their organellar cargo and the grooves they formed indenting MC, while this was consistent with holotomography. Discrete, rapid and highly localized transfer events, evidenced against a role for shed vesicles. Transfer coincided with rapid retraction of the cell-projections, suggesting a hydrodynamic mechanism. Increased hydrodynamic pressure in retracting cell-projections normally returns cytoplasm to the cell body. We hypothesize ‘cell-projection pumping’ (CPP), where cytoplasm in retracting cell-projections partially equilibrates into adjacent recipient cells via micro-fusions that form temporary inter-cellular cytoplasmic continuities. We tested plausibility for CPP by combined mathematical modelling, comparison of predictions from the model with experimental results, and then computer simulations based on experimental data. The mathematical model predicted preferential CPP into cells with lower cell stiffness, expected from equilibration of pressure towards least resistance. Predictions from the model were satisfied when Fib were co-cultured with MC, and fluorescence exchange related with cell stiffness by atomic force microscopy. When transfer into 5000 simulated recipient MC or Fib was studied in computer simulations, inputting experimental cell stiffness and donor cell fluorescence values generated transfers to simulated recipient cells similar to those seen by experiment. We propose CPP as a potentially novel mechanism in mammalian inter-cellular cytoplasmic transfer and communication.

**SIGNIFICANCE:** Time-lapse observations of co-cultured cells led us to hypothesize what we believe to be a novel hydrodynamic mechanism transferring cytoplasm between cells. Similar transfer by other mechanisms markedly affects cell behavior. Combined mathematical modelling, satisfaction of predictions from the mathematical model in cell culture experiments, and separate computer simulations that generate outcomes similar to experimental observations, support our hypothesized mechanism.

## INTRODUCTION

We earlier described the exchange of membrane and cytoplasmic protein between cultured human fibroblasts (Fib) and malignant cells (MC) (1). Others have made similar observations, and describe this as via either tunneling nanotubes (TNT) or exosomes and other shed membrane vesicles, and this is often associated with changes in cell phenotype (2-17), while we have also seen phenotypic change (1, 18, 19). Mitochondrial exchange has been considered particularly interesting (2, 4, 5, 20). TNT are long straight tube-like connections, typically suspended above the culture substrate, and establish cytoplasmic continuity between individual cells as a form of temporary partial fusion. They seem drawn out from pre-existing inter-cellular micro-fusions when touching cells migrate from one another, but may also form from fused adjacent filopodia (6-8, 12, 13, 21-23).

The current study was initially to examine the possibility that TNT accounted for our earlier observed inter-cellular transfer (1). Time-lapse confocal laser scanning microscopy (CLSM), however, revealed neither TNT or shed vesicles were involved, and led us to hypothesize what we believe to be a novel mechanism we term ‘cell-projection pumping’ (CPP). We now report these data, and describe and test our CPP hypothesis by a combination of: mathematical modelling; comparison of predictions from the model with experimental data; and comparison of computer simulation based on our mathematical model with experimental results.

With regard to CLSM results, it is important to appreciate necessity to use permanent labels, such as the fluorescent lipophilic markers 1,1’-dioctadecyl-3,3,3’,3’-tetramethylindodicarbocyanine perchlorate (DiD) and 3,3’-dioctadecyloxacarbocyanine perchlorate (DiO), to demonstrate total cytoplasmic transfer over time, because such labels accumulate and persist long after degradation of the originally labelled structures. By contrast, cell and organellar turn-over renders highly specific organellar or protein labels unreliable for detecting cumulative cytoplasmic transfer between cells (1). Also important, is that Fib have appreciably greater cell surface stiffness measured by atomic force microscopy (AFM) compared with MC (24). Further, punctate organellar labelling aids recognition of transfer events in time-lapse microscopy.

## MATERIALS AND METHODS

Information below is to aid interpretation of experiments. Detail to reproduce results is online in Supplemental Information.

### Cell culture and fluorescent labelling

Cell culture was as earlier described (1, 18, 19, 24, 25). Culture media for CLSM and AFM experiments were prepared by the Memorial Sloan-Kettering Cancer Centre Culture Media Core Facility (New York, NY). Culture reagents for holotomography were from (Sigma-Aldrich, St. Louis, USA), Bovogen (VIC, Australia), and CSL Biosciences, (VIC, Australia). Cell culture and microscopy reagents and plasticware were from established suppliers. Human dermal Fib (HDF) were from The Coriell Institute (Camden, NJ). SAOS-2 osteosarcoma cells were from the American Type Culture Collection (VA, USA). MM200-B12 melanoma cells were from The Millennium Institute (Westmead, NSW, Australia). DiD, DiO and Bacmam 2.0 Cell lights Nuclear-GFP baculovirus, were from Molecular Probes by Life Technologies (Grand Island, NY). Lipophilic DiD and DiO membrane label were used because: these markers are concentrated in organelles; punctate organellar labelling aids visualization of transfer events in time-lapse recordings; and organelles are suspended in cytoplasm so that organellar movement traces cytoplasmic flow. Labelling solutions of DiD for Fib (1mM) and DiO for MC (2mM) were applied to cells for 30 min in the case of DiD, and 1h for DiO. Monolayers were washed prior to overnight culture and further washing before co-culture. In some experiments, MM200-B12 were transfected with green fluorescent protein (GFP) expressing baculovirus.

### Co-culture conditions

Fib were seeded from 1 to 2 × 10^4^ cells per cm^2^ into either 25cm^2^ AFM culture plates (24), or culture well coverslips for overnight culture before labelling for AFM and CLSM experiments. Holotomography was with cells seeded in 35mm glass bottom dishes. MC were seeded at near confluence in either 25 cm^2^ flasks or 6 well culture plates prior to labeling. MC were then harvested with trypsin-EDTA and seeded over Fib in DMEM-α with BSA (4%) at 4 × 10^4^ cells per cm^2^ for up to 24 h co-culture.

### Time-lapse CLSM

Eight separate visual fields of Fib co-cultured with GFP labelled MM200-B12 were recorded for 25 h at 3 min intervals (1.13 mm^2^ surface area). Nine separate visual fields of DiO pre-labelled MM200-B12 were recorded for 8 h 15 min at 5 min intervals (0.76 mm^2^ surface area). Monolayers were fixed with paraformaldehyde after co-culture. CLSM was by a Zeiss LSM 5Live line-scanning confocal microscope.

### Time-lapse holotomography

Holotomography time-lapse images were collected at 2 min intervals using a Nano Live 3D Cell Explorer (Lausane, Switzerland). Recordings were of culture surface areas measuring 6.4 × 10^−3^ mm^2^ for: 6 h 10 min; 6 h 38 min; 18 h 58 min; and 19 h 24 min.

### Combined atomic force and fluorescence microscopy

An Asylum Research MFP-3D-BIO atomic force microscope coupled with a Zeis Axio Observer A1 fluorescence microscope was used for combined fluorescence-AFM recordings of randomly selected cells in paraformaldehyde fixed monolayers (24). Bright field and fluorescence images (DiD, DiO) were recorded prior to AFM scanning. A 1 μm AFM spherical polystyrene probe was used to record 16 x 16 points of force curves over 50 μm x 50 μm areas. Asylum Research, Software Version IX was used to determine Young’s modulus for each point by the Hertz model (24, 26). Fluorescence and AFM data were correlated for individual cells (24).

ImageJ software (http://imagej.net/Contributors) was used to analyze fluorescence images of individual cells: determining cell surface profile area, and summating red and green fluorescence separately. Fluorescence intensity in both channels was expressed in ‘Fluorescence Units’ (summated fluorescence / surface profile area). Cells were designated as having either ‘high’ or ‘low’ labelling from the opposing cell type. Median AFM stiffness was determined for individual cells, while stiffness fingerprints were also made of cells according to group to address sampling limitations as earlier described (24).

### Computer simulation of CPP between Fib and SAOS-2 populations

Simulations were of cells constructed with fluorescence and stiffness properties determined from experimental observations, and permitted to interact with each other at random as in co-culture. All simulation was in MATLAB (MATLAB by MathWorks Inc; scripts provided in Supplemental information).

Estimated cumulative distribution functions (ECDF) were developed from experimental median cell stiffness and fluorescence data (MATLAB Scripts provided in Supplemental Information). ECDFs were then used to generate simulated populations of cells with distributions for stiffness and fluorescence approximating experimental data. We generated 5100 donor Fib and SAOS-2, and 5000 recipient SAOS-2 and Fib respectively (Supplemental Information Fig. S1).

CPP was simulated by integration of curves for flow rate over time into receptor cells developed according to a mathematical model described in Results and detailed in Supplemental Information. Relevant variables for simulations were loaded into lists that had distributions bounded by target minimum and maximum values established before each simulation. Variables modelled in this way were: the number of Donor Cells A each Receptor Cell B could interact with; the number of transfer events possible between each Receptor Cell B and Donor Cell A; the cell-projection retraction rate *(U)* for each transfer event; cell-projection length at time 0 *(L*_*0*_); the radius *(r)* of each cell-projection; and the viscosity of cytoplasm *(η).*Values for these parameters were inferred from CLSM observations, with exception of *η* taken from the literature (27, 28). Time permitted for each transfer event was an exception, as maximum time = *L*_*0*_*/U*, and random choice was made from a pre-determined proportionate range between 0 and *L*_*0*_*/U.*

Average SAOS-2 and Fib cell height was determined from AFM data (3.89 × 10^−6^m and 2.36 × 10^−6^m respectively), while average SAOS-2 and Fib cell surface area (1.53 × 10^−9^m^2^ and 5.34 ×10^−9^m^2^ respectively) was by image analysis from separate experiments. These were used to calculate fluorescence from volume transfers.

Volume and fluorescence transfers for each simulated cell pairing were determined and summated simulation results compared with experimental results. Maximal pressure generated during individual simulated CPP events was also recorded. Distributions of input variables as well as simulation outcomes were plotted in histograms (Supplemental Information Figs. S2, S3).

## RESULTS AND DISCUSSION

### Transfer of organelles between Fib and MC

Organelles were clearly marked by both DiD and DiO. Organellar fluorescence overwhelmed plasma membrane labelling, so that plasma membranes were poorly defined by these labels. Obvious uptake of DiD labelled Fib organelles was in 11 of 106 DiO pre-labelled MC over 8 h 15 min co-culture, and 7 out of 71 GFP pre-labelled MC over 25 h co-culture (Figs. 1a, 2a,b; Supplemental Information movies S1, S2). Only once were DiO labelled organelles seen transferred from a MC to a Fib via broad cell-projections appearing as lamellipodia (Supplemental Information movie S3). While occasional large prominent Fib organelles were accepted by MC (Fig. 1a; Supplemental Information movie S1), most exchange was of smaller organelles, most readily seen when MC were labeled with DiO (Fig. 2; Supplemental Information movie S2). The precise identity of organelles transferred could not be defined from the images collected. However, the size and shape of the large organelles transferred was most consistent with mitochondria, and we have since verified that mitochondria can be exchanged by CPP in separate preliminary studies. We have no data on the specific identity of the smaller organelles below the size of mitochondria, and this awaits further study.

**FIGURE 1.**
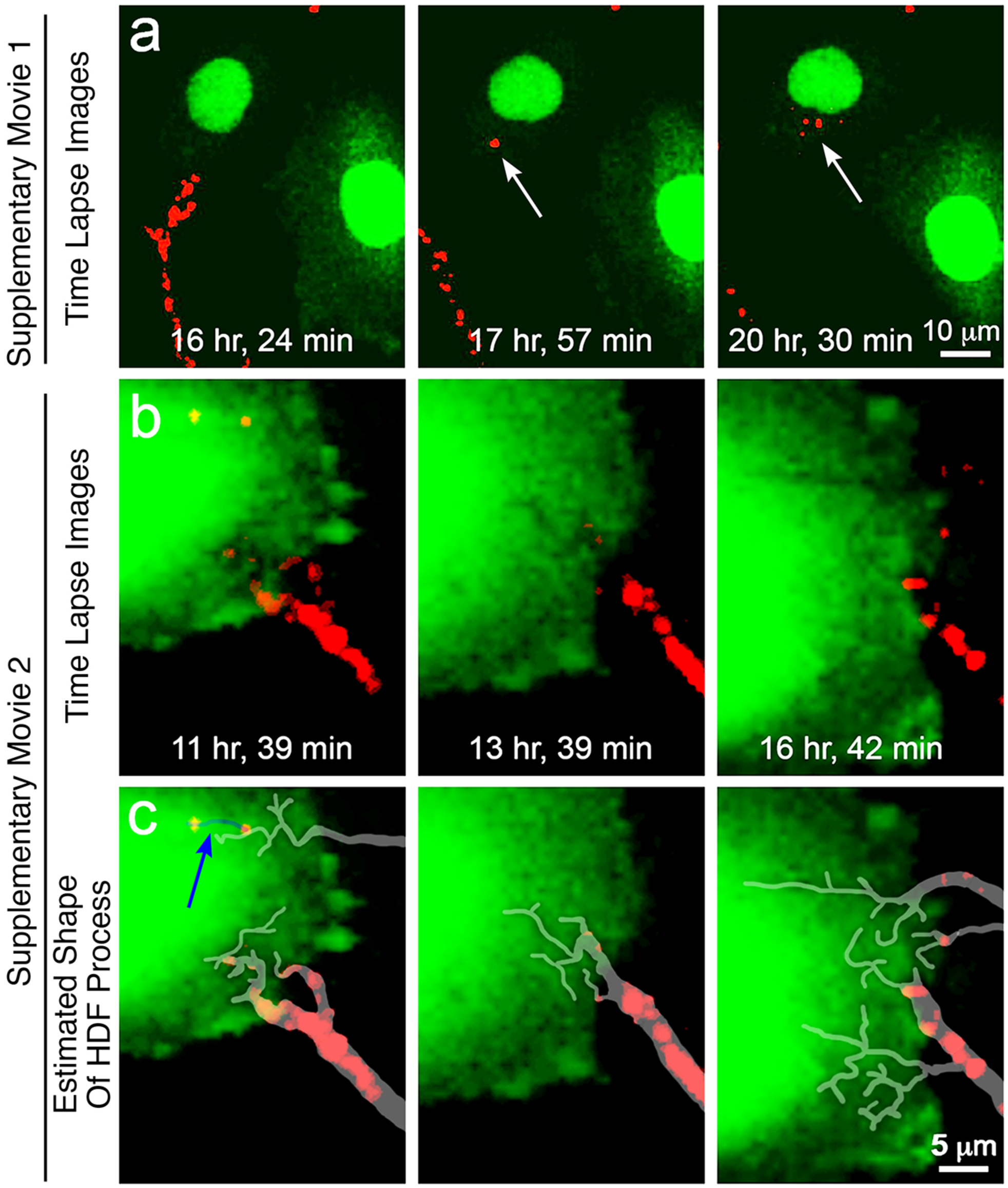
Frames from time-lapse CLSM recordings of Fib (pre-labelled DiD, red) co-cultured with MM200-B12 melanoma cells (GFP label); (Supplemental Information movies S1, S4) **(a)** A single large Fib organelle (red) was deposited into a green-labelled MM200-B12 cell (white arrow). **(b)** The location of Fib cell-projections was revealed by dark grooves made in the less stiff MM200-B12 cell, as well as by red-labelled organelles. **(c)** Inferred locations of cell-projections are marked with white transparency. One cell projection bearing red organelles speared into the MM200-B12 cell (blue transparency and arrow).

**FIGURE 2.**
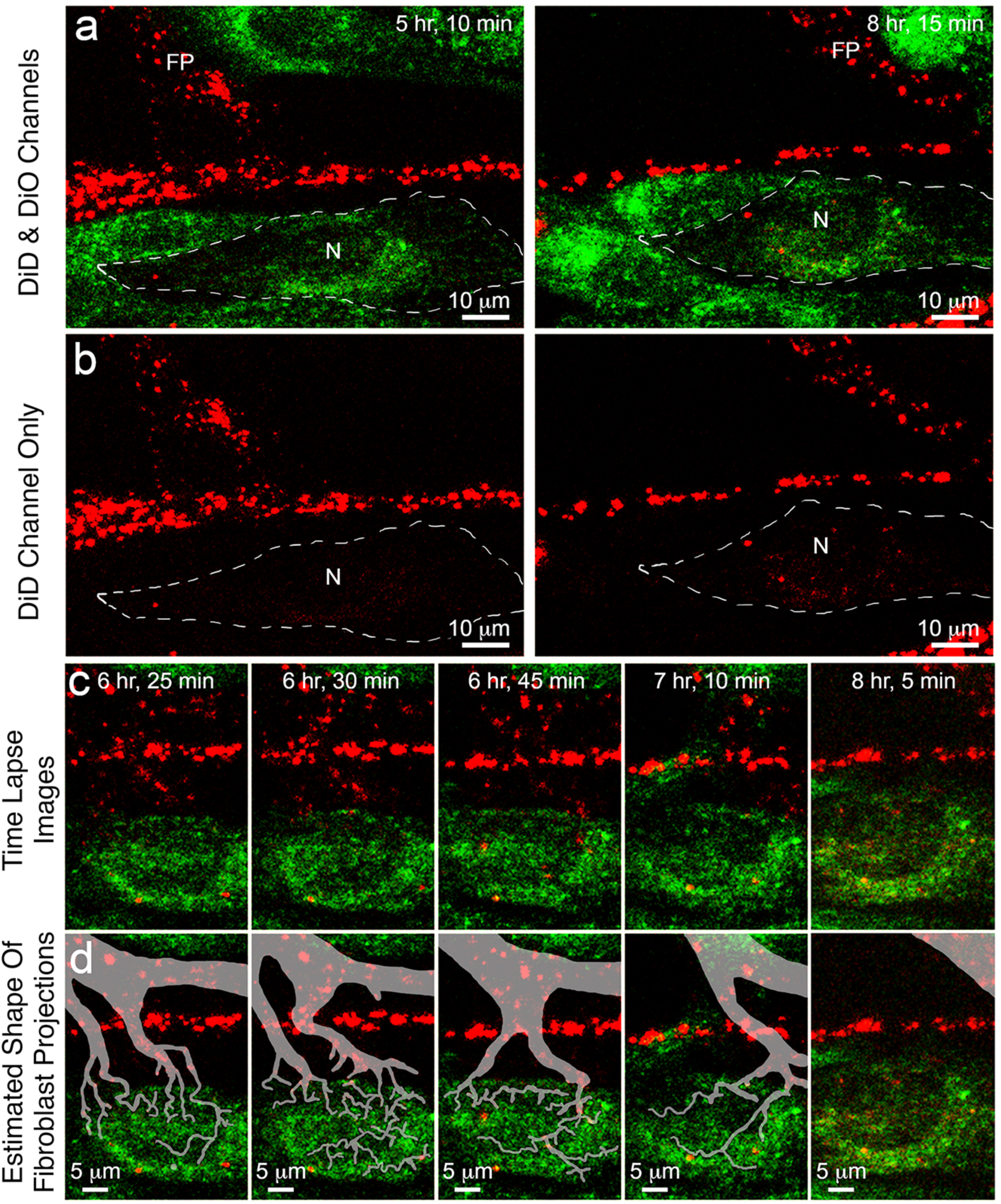
Frames from a time-lapse CLSM recording of Fib (pre-labelled DiD, red) co-cultured with MM200-B12 melanoma cells (pre-labelled DiO, green); (Supplemental Information movie S2). **(a, b)** A MC (white dashed outline) received red Fib organelles, readily seen when excluding the green channel (b). **(c)** This was from a broad Fib cell-projection (FP) that swept past the MC, indenting and grooving the recipient MC with numerous small branching cell-projections, all lost by 8 h 5min. **(d)** White transparency marks inferred locations of Fib cell-projections grooving the MC.

### Fine Fib cell-projections distinct from TNT transferred Fib organelles to MC

Fib cell-projections transferring organelles to MC could not be precisely resolved because they moved between time-lapse frames and were mostly transparent by CLSM, but general form was nonetheless inferred from CLSM z-stack images (Fig. 2c,d; Supplemental Information Fig. S4). Fib cell-projections were clearly more stiff than MC, and formed deep grooves indenting MC surfaces. These often contained DiD labelled cargo (Figs. 1b,c; 2c,d; Supplemental Information Fig. S4, movies S2, S4 and S5). These fine Fib cell-projections appeared as tree-like branching networks terminating in filopodia-like extensions (Figs. 1b,c, 2c,d; Supplemental Information Fig. S4, movies S2, S4 and S5). The presence of branching and rapidly moving filopodial structures that indented adjacent cells was confirmed by examination of time-lapse holotomographic microscopy (Fig 3, Supplemental Information Figs S5, S6). Field of view restrictions limited holotomography recordings to 28 cells, although the bodies of 20 of these cells were mostly visible for at least part of the time recorded. Organellar transfer observed by CLSM coincided with retraction events (Supplemental Information movies S1 to S3), and holotomography appeared consistent with this (Fig. 3, Supplemental Information movie S6). Examination of CLSM optical levels confirmed transferred organelles to be within and not on the surfaces of cells (Supplemental Information Figs S4, S7).

**FIGURE 3.**
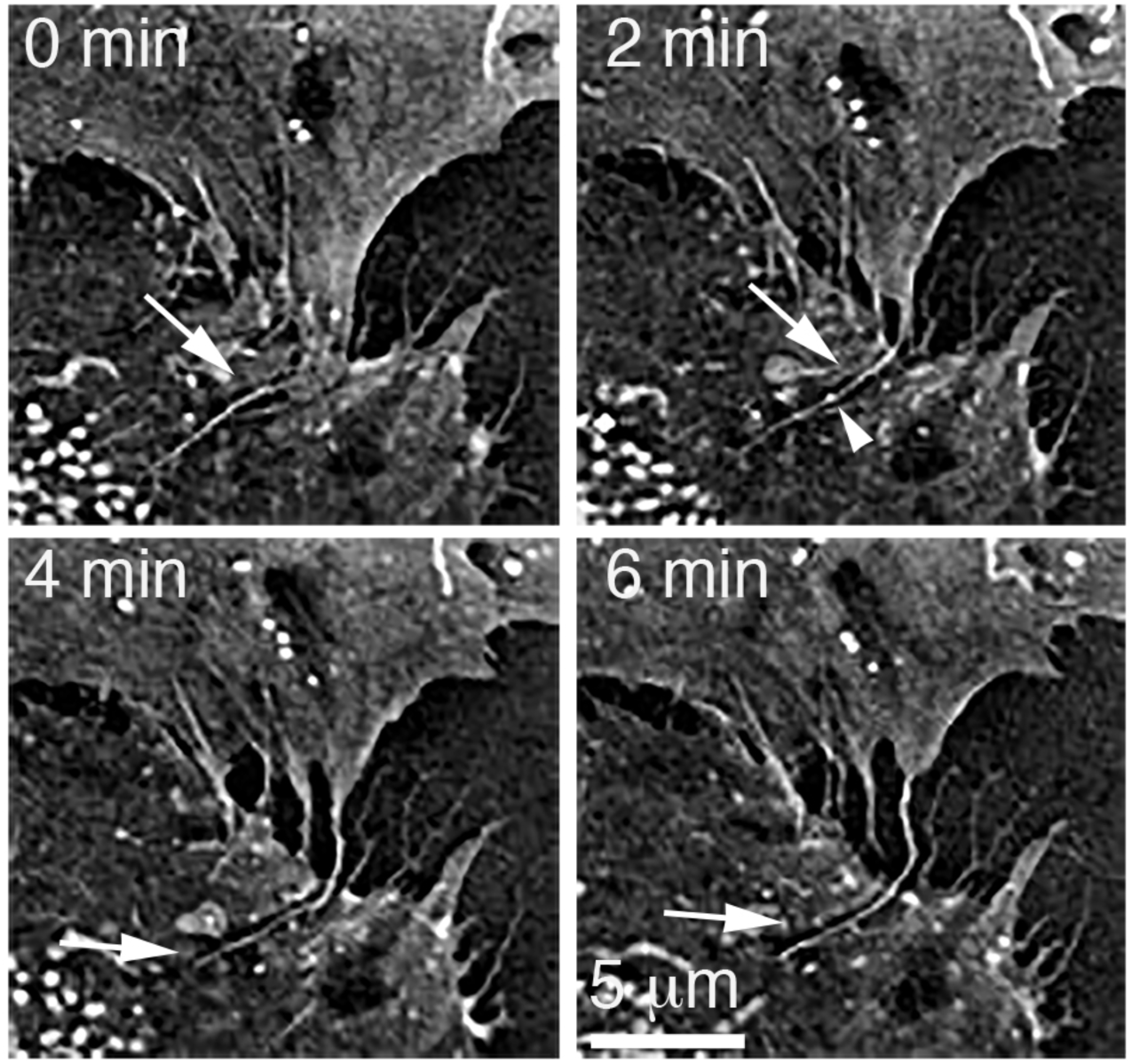
Frames from a time-lapse holotomographic recording showing a rapidly moving cell projection with possible transfer of an organelle in a co-culture of SAOS-2 with Fib; (Supplemental Information movie S6). A single optical level is shown at increasing time points. Numerous and sometimes branching cell-projections were seen, some on the plastic culture surface and others reaching onto the surface of the adjacent cell. Filopodia varied greatly with regard to mobility. One highly mobile cell-projection changed shape in adjacent frames (arrow), and between 2 and 4 min time points appeared to transfer bright organellar cargo (arrow head) to the adjacent cell.

TNT differ markedly from the transferring cell-projections in the current study (Supplemental Information Fig S8, S9). Unlike the transient, fast-moving, short, and branching structures here seen to be mechanically supported by the culture surface or cells; TNT often persist hours, extend long distances, are non-branching, and are suspended free above the culture surface (6, 7, 12, 13, 21-23). Cytoplasmic transfer via cell-projections in the size range of filopodia seems a novel function. Some precedent is, however, established by filopodial transfer of melanosomes, but precise details of melanosome transfer remain uncertain and may be by phagocytosis (29-31).

It was clear organellar transfers must have involved transient cytoplasmic micro-fusions between adjacent cells similar to those involved in TNT formation (6-8, 12, 13, 21-23), but in this instance occurring in cell-projections. The basis for this remains unknown.

### TNT, exosomes, fragmenting budding and non-specific label transfer did not account for observations

Occasional TNT were seen by both CLSM and holotomography, but contributed little to observed transfers (Supplemental Information Fig. S8, S9). If shed membrane vesicles had played a significant role, the ‘snowing’ of vesicles onto cells would have produced slow, diffuse and near uniform uptake of label, independent of cell-projection retraction. Instead, however, we saw: highly localized organellar transfer, with label uptake varying greatly between immediately adjacent cells; brief and rapid bursts of organellar transfer between individual time-lapse frames; and an association between retraction of cell-projections and transfer events. As such, images were inconsistent with either a role for shed membrane vesicles, or phagocytosis of occasional Fib fragments. Admixture of organelles with differing label within cells suggested multiple uptake events, while intimate physical contact between Fib and MC was insufficient for diffusion of DiD into MC (Supplemental Information Figs. S7, S10). We considered if donor cell fragments were ‘torn off’ and phagocytosed during cell-projection retraction. However, neither the budding of cell-projections, or formation of receptor cell surface spikes expected from such a mechanism, were ever seen.

### The CPP hypothesis

Lacking clear resolution of the cell-projections responsible for transfer, we hypothesized CPP as a mechanism to account for our data. Transient increases in hydrodynamic pressure occur during retraction of cell-projections, thus returning cytoplasm to cell bodies. Micro-fusions that establish intercellular cytoplasmic continuities are known to occur early in TNT formation, and should such a micro-fusion exist between a cell-projection and neighboring cell during retraction, raised pressure within the cell-projection partially equilibrates by cytoplasmic flow into the neighbor. Because fluid flows towards least resistance, relative differences in cell stiffness affect the extent of transfer (Fig. 4a).

**FIGURE 4.**
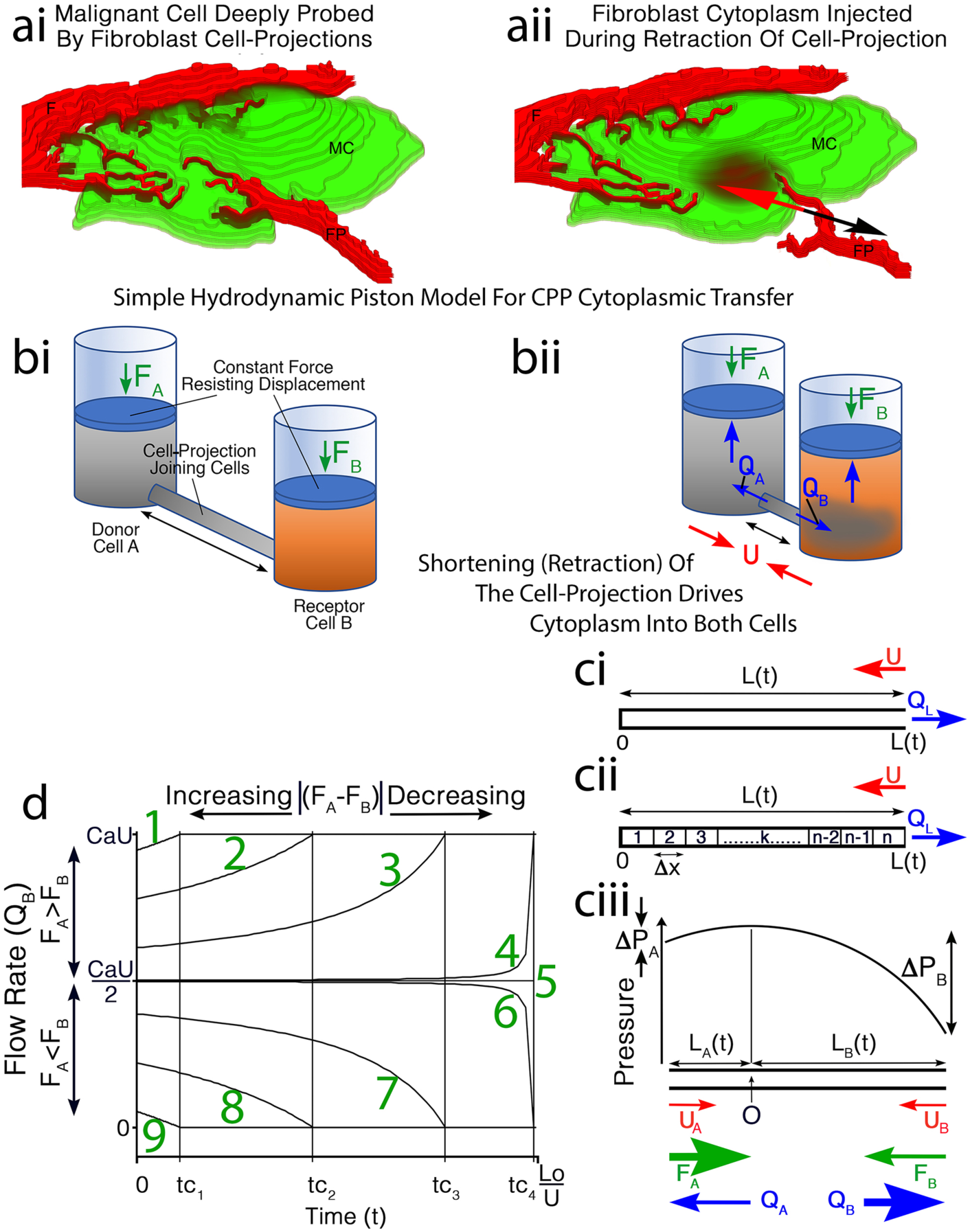
The CPP hypothesis. (**a)** Cartoon from a confocal image containing a MC deeply grooved by Fib (F) cell-projections (FP). **(aii)** If a micro-fusion establishes a transient inter-cellular cytoplasmic continuity, retraction of the cell-projection (black arrow) drives Fib cytoplasm into the MC (red arrow). **(bi)** This is modelled by piston-cylinders connected by a cylindrical tube (black arrow) containing fluid from the ‘Donor Cell’. **(bii)** Shortening of the tube at constant rate U (red arrows) mimics cell-projection retraction, driving contents into both ‘Donor’ and ‘Receptor Cells’ at flow rates Q_A_ and Q_B_ respectively (blue arrows), against constant reactive resistance forces proportionate to cell stiffness (F_A_, F_B_, green arrows). **(ci)** A close-ended tube with length L(t) contracts at constant rate U (red arrow), generating flow of fluid within the tube of Q_L_ (blue arrow). **(cii)** Dividing into n cylinders of Δx length, with Δx approaching zero, relates L(t) to pressure and Q_L_. **(ciii)** Two such tubes of lengths L_A_(t) and L_B_(t) are opened and joined at their origins (O), contracting at U_A_ and U_B_ (U_A_+U_B_=U) (red arrows), with flow (Q_A_ and Q_B_, blue arrows) against constant forces (F_A_ and F_B_, green arrows). Pressure is maximal at O, and equals F_A_ and F_B_ at the open ends, establishing ΔP_A_ and ΔP_B_. Where F_A_>F_B_, L_A_(t)<L_B_(t) and Q_A_<Q_B_; this reverses when F_A_<F_B_; while when F_A_=F_B_, Q_A_=Q_B_. **(d)** Q_B_ is plotted over time for retraction to extinction of a cell-projection in 9 separate circumstances where all variables are identical except for F_A_ and F_B_, to generate 9 separate ‘curves’ as labelled. Notably, when F_A_>F_B_, there is a time (tc) when L_A_=0, Q_A_=0, and remaining flow is Q_B_=CaU, where Ca= tube cross-sectional area. When F_A_<F_B_, L_B_=0 at tc, so that Q_B_=0, and remaining flow is Q_A_. Decreasing F_A_-F_B_ increases tc.

### Mathematical model for CPP

CPP was described mathematically for a simple two chamber hydrodynamic system joined by a cylindrical connector, each chamber representing either Donor Cell A or Receptor Cell B, and the cylindrical connector representing a cell-projection from Cell A. We assume a constant rate of cell-projection retraction modelled by a constant rate of shortening for the cylindrical connector (U), that expels fluid into both Donor Cell A and Receptor Cell B at flow rates *Q*_*A*_ and *Q*_*B*_ respectively (Fig. 4b). Only *Q*_*B*_ was determined, since it is only *Q*_*B*_ which delivers cytoplasm to the opposing cell (Fig. 4b). Stiffness of donor Cell A and Receptor Cell B are considered proportionate to the reactive force of resistance to flow for Cells A and B respectively *(F*_*A*_, *F*_*B*_).

We applied the well described Hagen-Poiseuille relationships, where resistance to flow in a cylindrical tube *(ρ)* is given by Eq. 1, and the flow rate *(Q)* of a Newtonian fluid of viscosity *(η)* through a cylindrical tube of length *(L)* with radius *(r)*, due to a pressure difference *(ΔP)* is as per Eq. 2 (32).

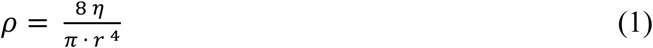

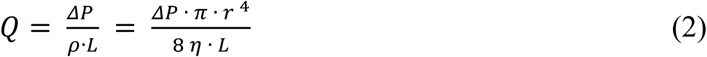

Bulk modulus was assumed to be negligible, as the fluid volumes exchanged are very small and cytoplasm was assumed to behave as a Newtonian fluid. Uptake of cytoplasm by receptor cells via CPP was assumed to have negligible effect on receptor cell cytoplasmic viscosity and cell stiffness. In absence of highly detailed structural data on the cell-projections that mediate CPP, and for purposes of necessary simplification, we have modelled cell projections as simple contracting cylindrical tubes. The hydraulic system is not confined but consists instead of two ‘open’ halves, each of which operates against a separate constant reaction force term in the form of a reaction pressure, *F* (*F*_*A*_ and *F*_*B*_ for Donor and Receptor cells respectively) (Fig. 4). Hence each half has a separate *ΔP* as also demanded by Pascal’s law (33).

A central component of the CPP mechanism is resistance to cytoplasmic flow into each of the cells. We first considered if cells offered Hookean spring-like resistance to flow, but initial investigation demonstrated that this could not explain the experimental data. Instead, we have considered that cytoplasmic resistance behaves similarly to a Bingham plastic, where viscous flow occurs only when external force exceeds a defined yield point. We have assumed that for each cell, this yield point is proportional to the median stiffness of the cell measured by AFM, and we suggest this reflects disruption of bonds between either or both cytosolic or cytoskeletal elements (34). This contrasts with the simple viscous flow we assume for contents of cell-projections, on basis of the greatly simplified internal structure of cell-projections compared with cell bodies.

Consequently the effect of each cell on the flow from the projection can be accounted for by a single force term equivalent to the yield point *(P*^*Y*^), of the cell cytoplasm. These force terms are the reaction pressures *F*_*A*_ and *F*_*B*_. The yield point occurs when the deformation of the material under stress reaches a critical level. Assuming that this critical deformation is approximately the same for both cells, the yield point can be assumed to be proportional to the stiffness of the cell (*S)*, as measured by AFM. If *Z* is the constant of proportionality then *P*_*A*_^*Y*^*=ZS*_*A*_, and *P*_*B*_^*Y*^*=ZS*_*B*_.

It seems reasonable to assume only modest effect of the simplifying assumptions outlined above, in diverging calculated estimations made in the current study from events in-vivo.

The mathematical approach is summarized below and details are in Supplemental Information. A cylindrical tube with cross sectional area *(Ca)* is closed at one end and contains a Newtonian fluid. It undergoes constant contraction at rate *(U)* and has length *L(t)* at time *(t)* as measured from ‘0’ at it’s origin (Fig. 4cii). By dividing the cylinder into n segments of length *(Δx)*, considering the contribution of each segment to the total flow expelled from the contracting tube, and letting *Δx* approach 0, an expression relating flow to the position *(x)* along the tube is derived. Eqs. 1 and 2 are used to develop an expression for pressure drop along the length of the tube as per Eq. 3.

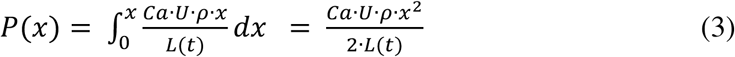

The above outlined system is replicated, and one of these elements is rotated so that the two tubes abut end to end, sharing the same origin. This now represents a cell-projection joining Donor Cell A ‘at left’, with Receptor Cell B ‘at right’. The origin represents a point (Po) where during contraction of the cell-projection, there is maximum pressure *(P*_*max*_) and no flow. The length of tube between the origin and Donor Cell A is *L*_*A*_, and that to Receptor Cell B is *L*_*B*_, with rates of contraction *U*_*A*_ and *U*_*B*_ respectively. The entire system has length *L(t)* and rate of retraction *U = U*_*A*_ *+ U*_*B*_ (Fig. 3ciii). The tube is open to Cells A and B, and flow in both directions is resisted by constant reaction pressures *F*_*A*_ and *F*_*B*_ (Fig. 4ciii). These reaction pressures are equal to the yield points *P*_*A*_^*Y*^ and *P*_*B*_^*Y*^. Where *F*_*A*_*=F*_*B*_, then *L*_*A*_*(t) =L*_*B*_*(t) = L(t)/2*, and both cells receive equal flow at CaU/2. Where *F*_*A*_ ≠ *F*_*B*_ total flow into Receptor Cell B *(Q*_*B*_) is given by Eq. 4 up to time *tc*.

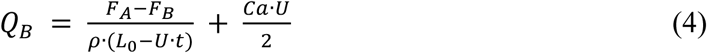

The detailed basis for the impact of differences between *F*_*A*_ and *F*_*B*_ on determining both *tc* and *Q*_*B*_ are described in Supplemental Information. Note that *P*_*max*_*(t)* decreases with time (Fig. 4ciii), so where *F*_*A*_*>F*_*B*_, at time *tc*: *L*_*A*_*(t)* reaches 0; *P*_*max*_*(t)* reaches the yield point of cell A (*P*_*A*_^*Y*^) so that no further flow into Cell A is possible; and all remaining flow is to the right at *Q*_*B*_ = CaU, with the effect that Receptor Cell B receives more flow than Donor Cell A (Fig. 4ciii). The reverse applies when *F*_*A*_*<F*_*B*_, in that *P*_*max*_*(t)* reaches the yield point of Cell B at time *tc*, when *L*_*B*_ is extinguished, and all remaining flow is to Cell A at *Q*_*A*_ = CaU.

The behavior of the system is apparent from Fig. 4d, which shows *Q*_*B*_ as a family of ‘curves’ numbered 1 to 9 for varying values of *F*_*A*_ and *F*_*B*_ from *t* = 0 to *t = L*_*0*_*/U.* Note that *L*_*0*_*/U* defines the maximum time during which flow *Q*_*B*_ is possible, because the cell-projection is extinguished at that time. Similarly, *Q*_*B*_ cannot exceed CaU. For all values of *F*_*A*_ and *F*_*B*_, total summated flow into both Cells A and B *(Q*_*T*_) at any time *(t)* equates to CaU, such that *Q*_*T*_ *= Q*_*A*_ *+ Q*_*B*_ *=* CaU. Where *F*_*A*_ *= F*_*B*_, *Q*_*A*_ *= Q*_*B*_ = CaU/2 at all time points, to give a horizontal line crossing Fig. 4d (line drawn as ‘curve’ 5). ‘Curves’ 1 to 4 drawn above ‘curve’ 5 at CaU/2, show *Q*_*B*_ where *F*_*A*_ *> F*_*B*_, while ‘curves’ 6 to 9 drawn below the horizontal at CaU/2 (5) show *Q*_*B*_ where *F*_*A*_ *< F*_*B*_.

Considering Fig. 4ciii illustrating *ΔP* along the length of a cell-projection where *F*_*A*_ *> F*_*B*_, both *ΔP*_*A*_ and *ΔP*_*B*_ reduce at the same rate while the cylinder shortens, but because *F*_*A*_ *> F*_*B*_, *ΔP*_*A*_ is extinguished before *ΔP*_*B*_, and this is at time tc when: *F*_*A*_ = maximum pressure at ‘point 0’ *L*_*A*_ *= 0*; flow into Cell A ceases; and all remaining flow is into Cell B at *Q*_*B*_ = CaU. In effect, ‘point O’ shifts to the left during contraction of the cell-projection, and time *tc* is the moment when ‘point O’ meets the opening of the cell-projection into Cell A. Please note that Fig. 4ciii illustrates circumstances when *F*_*A*_ *> F*_*B*_, and relates to ‘curves’ above CaU*/2* in Fig. 4d (‘curves’ 1 to 4). Where *F*_*A*_ *< F*_*B*_, *L*_*A*_ is larger than *L*_*B*_, and *L*_*B*_ extinguishes before *L*_*A*_, to give ‘curves’ such as those illustrated in Fig 4d below *Q*_*B*_ = CaU/2 (‘curves’ 6 to 9).

Although Fig. 4d shows ‘curves’ for Q_B_ with differing *F*_*A*_ and *F*_*B*_, because *Q*_*A*_ + *Q*_*B*_ = CaU, the curve for *Q*_*A*_ of the system is always given by symmetrical reflection of the ‘curve’ for *Q*_*B*_ about the horizontal at CaU/2. From this, if conditions are such that ‘curves’ 1, 2, 3 and 4 are for either *Q*_*B*_ or *Q*_*A*_, then *Q*_*A*_ and *Q*_*B*_ are each respectively given by ‘curves’ 9, 8, 7 and 6. This is consistent with the paired symmetry of ‘curves’ 1, 2, 3 and 4 with respectively 9, 8, 7 and 6 expected for *Q*_*B*_, where the absolute values for *F*_*A*_*-F*_*B*_ are the same. Decreasing values for *F*_*A*_*-F*_*B*_ drive the system towards perfect symmetry and constant *Q*_*B*_ at CaU/2. Increasing values for *F*_*A*_*-F*_*B*_ drive the system towards the extremes for *Q*_*B*_ at either CaU or 0, dependent if *F*_*A*_ is larger or smaller than *F*_*B*_ respectively. These extreme values for *Q*_*B*_ are reached at time *tc*, which approaches 0 as *F*_*A*_*-F*_*B*_ increases. Integration of ‘curves’ for *Q*_*B*_, gives total volumes transferred to Cell B throughout retraction of the cell-projection.

### Predictions from the mathematical model

Examination of Eqs. 2 and 4 identifies variables that when raised, predict increased *Q*_*B*_ (*ΔP, F*_*A*_*-F*_*B*_, *Ca, U, r*) and decreased *Q*_*B*_ (*ρ, η, L*_*0*_). Of these, by far the most influential variable is *r*, reflecting the power function in Eq. 2.

Imagine two cell populations D and E, exchanging cytoplasm with each other via CPP. Variability in cell stiffness within each of the cell populations D and E is represented by a range of values for *F*_*D*_ and *F*_*E*_ respectively. Assume that there are an equal number of transfers from Cells D to E, as there are from E to D, and that all other variables for CPP in Eq. 4 are the same for all transfers from Cells D to E, as they are for transfers from Cells E to D.

Considering Fig. 4d, in the particular circumstance where the range and distribution of values for *F*_*D*_ and *F*_*E*_ are identical, if all other relevant variables are also identical, then transfers from Cells D to E are essentially mirrored by transfers from Cells E to D, and ‘curves’ for *Q*_*B*_ into both populations will be as often above the horizontal (‘curves’ 1, 2, 3 and 4) as below (‘curves’ 6, 7, 8 and 9). No preference in the distribution of volume transfers would be seen between the two cell populations.

On the other hand, if Cells D have a distribution of cell stiffness values such that *F*_*D*_ is often greater than *F*_*E*_, then there will be skewing of the frequency with which different ‘curves’ for *Q*_*B*_ arise. For CPP transfers from Cells D to E, amongst the ‘curves’ 1 to 9 drawn in Fig. 4d, the frequency of their occurrence would reduce with increasing ‘curve number’. This would contrast with CPP transfers from Cells E to D, where the frequency of occurrence of ‘curves’ drawn 1 to 9 in Fig. 4d would increase with increasing ‘curve number’. Because volumes transferred reduce with increasing ‘curve number’ as drawn in Fig. 4d, and also because there is skewing of ‘curve number’ downwards for transfers from Cells D to E, as opposed to skewing of ‘curve number’ upwards for transfers from Cells E to D, the model predicts that there is preferential CPP transfer from populations of cells with high cell stiffness, to populations of cells with low cell stiffness.

A further prediction can also be made from Fig. 4d. Imagine two cells, A and B connected as in Fig. 4b, and experiencing CPP till extinction of the cell projection. In the first instance, let *F*_*A*_ be very much greater than *F*_*B*_, so that the ‘curve’ drawn as ‘1’ in Fig. 4d shows *Q*_*B*_ to be at the maximum of CaU for most of the time during which cell-projection retraction occurs. Let all conditions be the same, except that *F*_*B*_ is increased, initially modestly so that ‘curve’ 2 now shows *Q*_*B*_, and then in separate cases where FB is further increased to generate ‘curves’ 3 and 4 for *Q*_*B*_. Once *F*_*B*_ has risen to be equal to *F*_*A*_, ‘curve’ 5 for *Q*_*B*_ appears as a horizontal at CaU/2. Allowing *F*_*B*_ to increase further, generates first ‘curve’ 6, and then ‘curves’ 7 to 9 as *F*_*B*_ rises further still. Integrating curves 1 to 9 shows that the total volume transferred by CPP reduces with increasing *F*_*B*_. From this, an inverse relationship is predicted between the stiffness of individual cells within any given population of receptor cells, and the volume of cytoplasm acquired by CPP from the partner donor cell population.

These predictions are illustrated in computer simulations shown in Fig. S11 of Supplemental Information, showing how cell stiffness relates to volume and fluorescence transfer.

We earlier showed that Fib have higher but overlapping stiffness compared with SAOS-2 (24), so that based on the above, the mathematical model predicts: a) preferential CPP transfer of fluorescent label from Fib to SAOS, compared with transfer from SAOS-2 to Fib; b) an inverse correlation between SAOS-2 stiffness and CPP fluorescence uptake from co-cultured Fib; and c) an inverse correlation between Fib stiffness and CPP fluorescence uptake from co-cultured SAOS-2.

### Experimental measurement of cell stiffness and fluorescence in co-cultured cells satisfied predictions of the mathematical model

We studied SAOS-2, because most of our earlier and subsequent work on phenotypic effects of transfer has been with this cell line (1, 18, 19, 25). Fluorescence microscopy of a co-culture of Fib with SAOS-2, revealed the most evident transfer of fluorescent label was from Fib to SAOS-2, with less obvious fluorescence transfer from SAOS-2 to Fib (Fig. 5a). Although stiffness varied greatly across surfaces of individual cells, stiffness fingerprints confirmed Fib were stiffer and had lower cell height compared with SAOS-2 (Mann Whitney U Test, p < 0.0001) (Fig 5b,c,d). These data thus satisfied the prediction that there would be preferential CPP transfer of fluorescent label from Fib to SAOS, compared with transfer from SAOS-2 to Fib.

**FIGURE 5.**
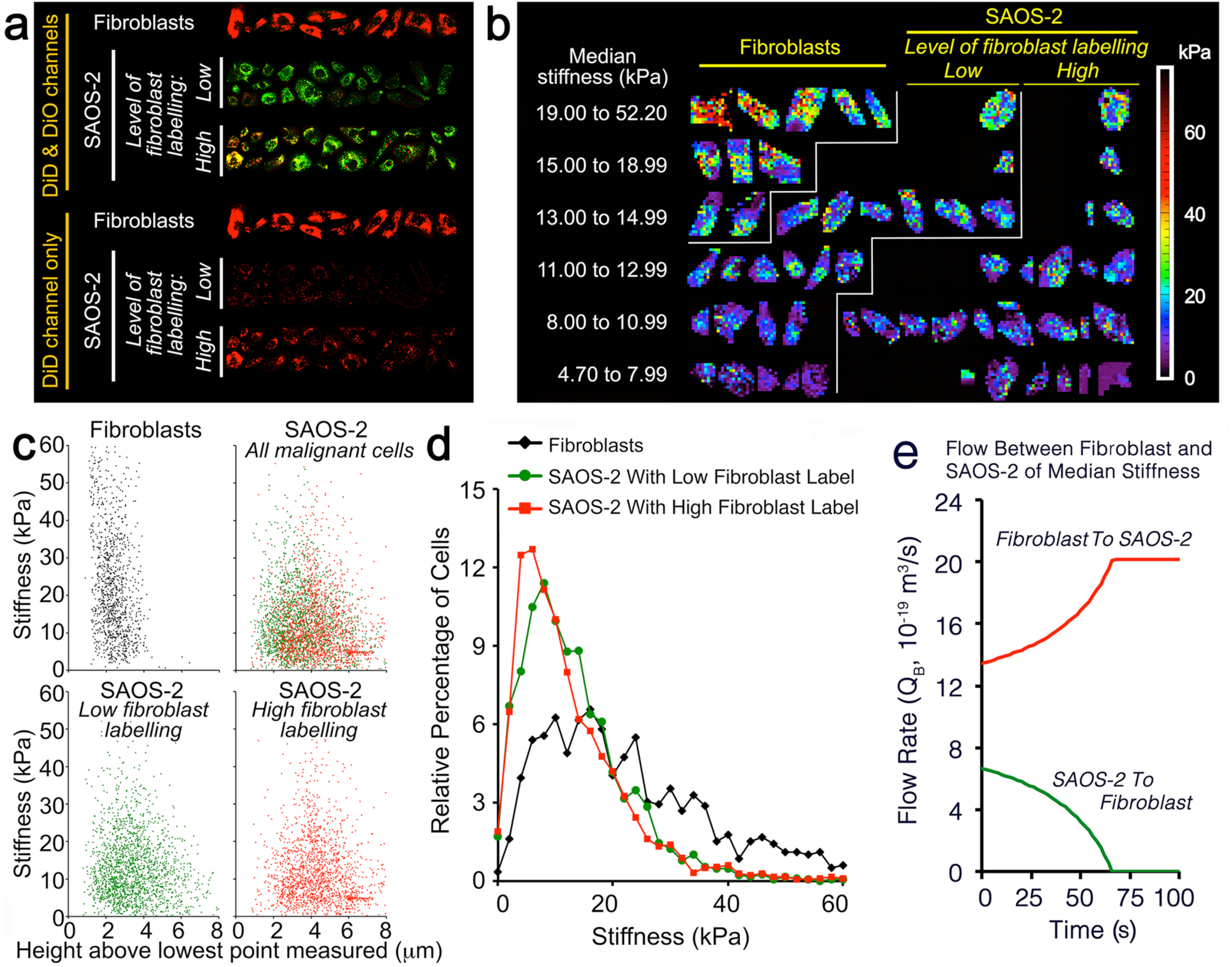
Relationship between cell stiffness and fluorescence in SAOS-2 (pre-labelled DiO, green) co-cultured for 24 h with Fib (pre-labelled DiD, red). (**a**) DiD levels were quantitated to identify Fib, SAOS-2 with high Fib label, and SAOS-2 with low Fib label. (**b**) Cells are grouped and separated by stair-lines according to identity as being either: Fib (top left group); SAOS-2 with low Fib labelling (middle group); or SAOS-2 with high Fib labelling (right hand group). Within each group, cells are arranged in tiers of increasing median AFM stiffness. Median AFM stiffness varied greatly within these groups, although Fib were generally stiffer than SAOS-2, while SAOS-2 with high Fib label were usually less stiff than SAOS-2 with low Fib label. (**c**) Stiffness fingerprints supported this, showing Fib (black dots) had greater stiffness and lower cell height than SAOS-2 (red and green dots) (Mann Whitney U Test, p < 0.0001). Further, stiffness of SAOS-2 with low Fib labeling (green dots) was higher than for SAOS-2 with high Fib labelling (red dots), and the reverse applied for height measures (Mann Whitney U Test, p < 0.0004). (**d**) Relative percentage distribution plots binned at 2 kPa for stiffness. **(e)** CPP was modelled between a Fib and SAOS-2, each of median cell stiffness, examining exchange from Fib to SAOS-2 (red), as well as from SAOS-2 to Fib (green). Results were consistent with the CPP hypothesis.

Label transfer varied amongst SAOS-2, with some SAOS-2 having high and others negligible Fib labeling (Fig. 5a). SAOS-2 with high Fib labeling had lower stiffness compared with SAOS-2 with low Fib labeling (Mann Whitney U Test, p < 0.0004) (Fig. 5b,c,d). In addition, higher uptake of fluorescence correlated with greater cell height (Mann Whitney U Test, p < 0.0004). These data thus satisfied the further prediction of an inverse correlation between SAOS-2 stiffness and CPP fluorescence uptake from co-cultured Fib.

Fig 5e shows CPP flow rates for transfer between a Fib and SAOS-2 cell, each with median stiffness (18,765 Pa 11,181 Pa respectively), considering each cell in turn as donor or receptor, and applying biologically reasonable assumptions for: cell-projection retraction rate (1 × 10^−6^m/s), viscosity (2.5 × 10^−3^ Pa.s), length at time zero (100 μm), and radius (0.8 μm) of the cell-projection. Significant flow was calculated from the mathematical model, and transfer from the Fib exceeded that from the SAOS-2, consistent with predicted preferential CPP transfer from Fib to SAOS-2.

Similar to observations in SAOS-2 (Fig. 5), uptake of fluorescence from SAOS-2 by Fib was negatively correlated with Fib cell stiffness, and there was also a positive correlation with cell height (Mann Whitney U Test, p < 0.0001) (Fig. 6). This was consistent with the prediction of inverse correlation between Fib stiffness and CPP fluorescence uptake from co-cultured SAOS-2.

**FIGURE 6.**
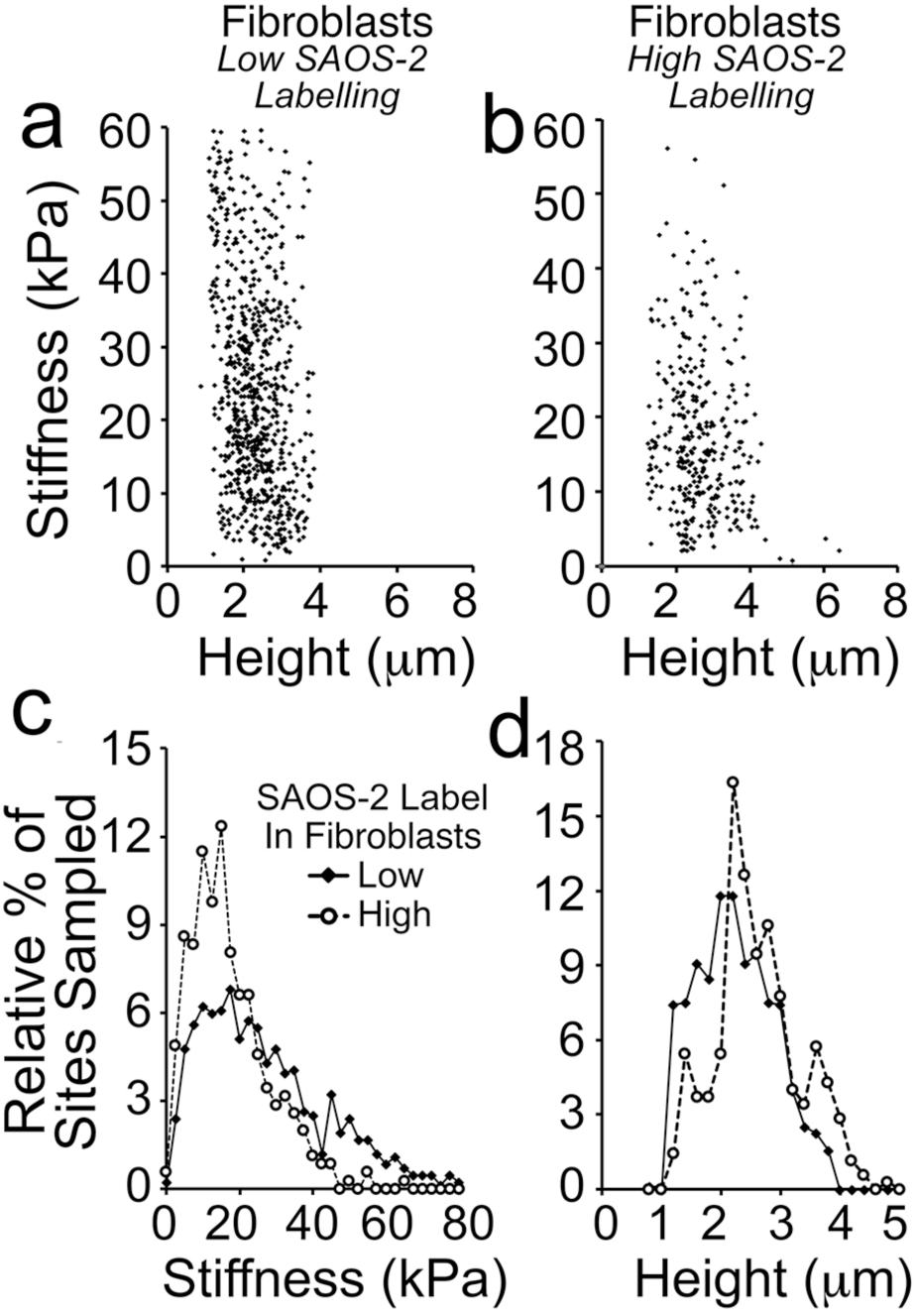
Stiffness fingerprints and proportional distribution plots of AFM stiffness and height records for Fib in co-culture with DiO pre-labelled SAOS-2 according to level of SAOS-2 labelling. **(a**,**b)** Fib with low SAOS-2 labelling had higher stiffness and lower cell height measurements (a), compared with Fib with high SAOS-2 labelling (b) (Mann Whitney U Test, p < 0.0001). **(c**,**d)** Proportional distribution plots binned at 2.5 kPa for stiffness and 0.2 μm for height measurements, confirmed the visual impressions from stiffness fingerprints.

Experimental results thus satisfied all predictions from the mathematical model.

### Results of numerical MATLAB simulations agreed with experimental observations

One objective of computer simulations was to determine if it was possible to explain experimental observations, applying biologically reasonable assumptions to our mathematical model. Simulations concorded well with experimental results, and also satisfied predictions of the model. Applying biologically reasonable assumptions (Supplemental Information, Figs. S2, S3), in simulations gave fluorescence transfers that closely approximated experimental fluorescence transfers (Fig. 7a,b). Median fluorescence levels of donor and receptor cells are shown in Table 1, and demonstrate similarity between experimental and simulated results, as well as proportionately more transfer from Fib to SAOS-2 than in the reverse direction. The inverse relationship between receptor cell stiffness and uptake was seen in simulations for transfer to both Fib and SAOS-2 (Fig. 7c,d).

**Table 1.** Median fluorescence levels from experimental observations and simulations.

|  | Median Fluorescence per Cell<br>(Fluorescence Units) |  |  | Receptor Fluorescence /<br>Donor Fluorescence |  |
| --- | --- | --- | --- | --- | --- |
|  | Donor<br>Cells | Experimental<br>Receptor Cells | Simulated<br>Receptor Cells | Experimental | Simulated |
| Red (DiD)<br>Fluorescence | 16.904 | 0.641 | 0.630 | 0.038 | 0.037 |
| Green (DiO)<br>Fluorescence | 31.849 | 0.504 | 0.414 | 0.016 | 0.013 |

Overall fluorescence levels for simulated receptor cells was comparable to that seen in experimental results. When expressed as ratios relative to donor cell fluorescence, Fib which were the receptors for green DiO fluorescence, had lower uptake relative to red DiD fluorescence by SAOS-2 receptor cells. Proportionate uptake was similar between simulation results and experimental data, with fluorescence uptake of SAOS-2 being 2.40 fold that of Fib (0.038/0.016) by experiment, and 2.81 fold that of Fib by simulation (0.037/0.013).

**FIGURE 7.**
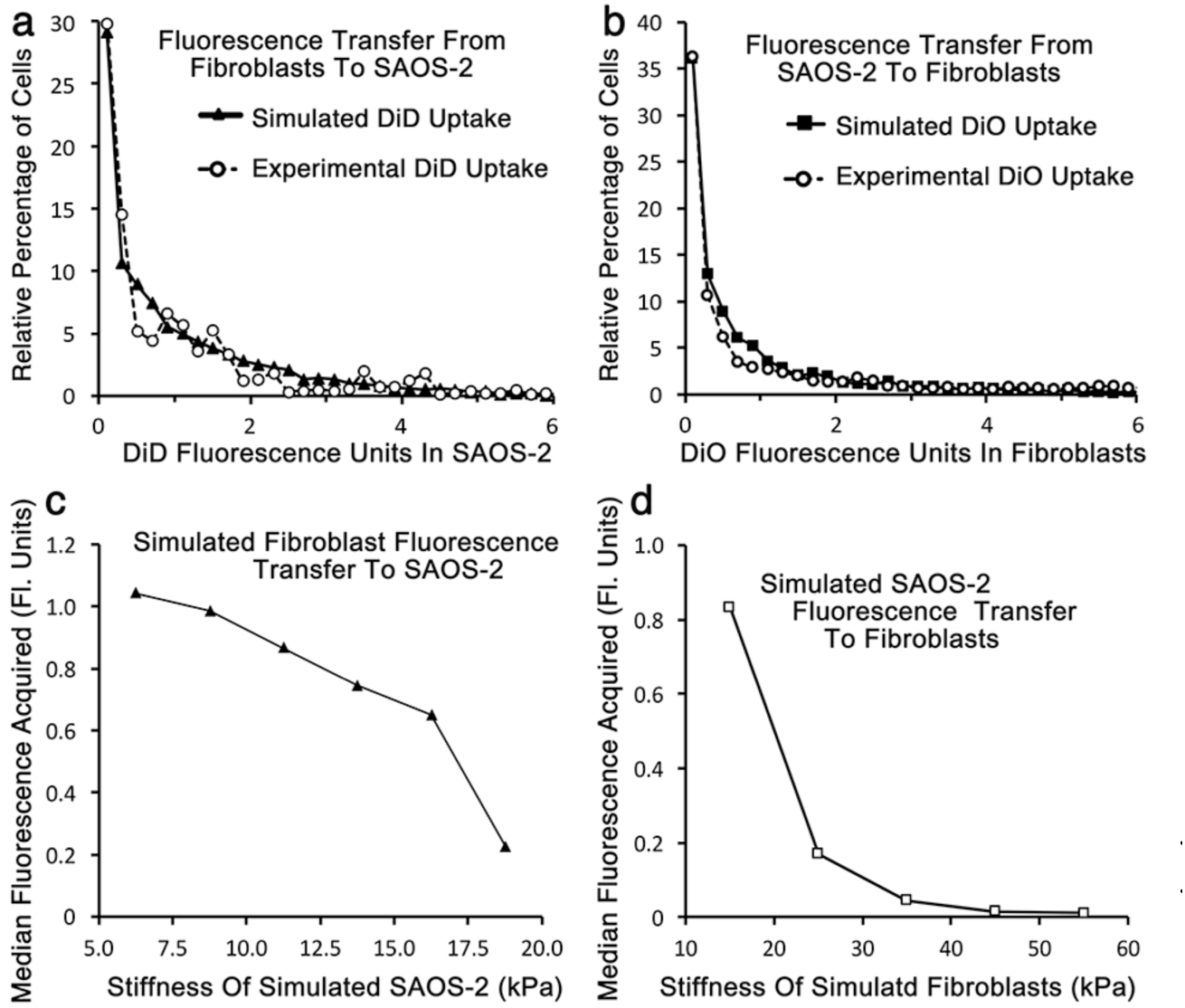
Comparison of simulated with experimentally observed transfer and the relationship between simulated recipient cell stiffness and fluorescence uptake. **(a**,**b)** Experimental and simulated results had good concordance where: each SAOS-2 had from 1 to 3 donor Fib (1 to 3 for Fib recipients); there were from 0 to 2 exchange events from each Fib to each SAOS-2 (3 to 8 for recipient Fib); cell-projection retraction ranged from 0.5 to 1.4 μm/s for donor Fib (1 to 5 μm/s for donor SAOS-2); the proportion of maximal possible time for individual transfer events was from 0 to 0.9 for Fib cell-projection retraction (0.6 to 0.9 for SAOS-2 cell-projection retraction); the length of donor Fib cell-projections was from 5 to 120 μm (40 to 90 μm from donor SAOS-2); the radius of Fib donor cell-projections was from 0.55 to 1.75 μm (0.7 to 2.5 μm for donor SAOS-2); and the viscosity of cytoplasm was from 1.5 to 4.0 mPa.s for both donor cells. **(c**,**d)** An inverse relationship between SAOS-2 stiffness and median fluorescence acquired by CPP was seen (c), with a similar result for Fib receiving SAOS-2 fluorescence (d) (p < 0.0001, Mann Whitney U Test).

Calculated pressures for these simulated transfers were generally modest (for transfer from Fib to SAOS-2: median 0.58 Pa, 7.48 × 10^−5^ Pa to 4.39 Pa; for transfer from SAOS-2 to Fib: median 0.377 Pa, 8.51 × 10^−6^ Pa to 6.35 Pa). Predominantly low pressures required to account for results, support plausibility for CPP.

### Simulations predicted large proportional volume transfers between cells

Donor cell fluorescence is highly variable (Supplemental Information Fig. S1) (1), so that direct estimation of CPP volume transfers required to account for fluorescence levels in recipient cells was not previously possible. Nonetheless, this difficulty was overcome by the current computer simulations, where the effect of variable donor cell fluorescence was included in calculation.

Volume exchange was expressed as percentages relative to the average volume of a single Receptor Cell B, and the distribution of cells according to volumes transferred in the simulations shown in Fig. 7, are illustrated as histograms in Fig. 8. More detail is available in Supplemental Information (Table S1). Most simulated recipient cells accepted appreciable donor cell cytoplasm, consistent with visual impressions (1) (Figs. 2,5). Amongst simulated SAOS-2, 55.5% had over 3% volume acquired from simulated Fib, and 5.2% of simulated SAOS-2 had over 19% volume acquired from simulated Fib. Consistent with occasional experimentally observed SAOS-2 with very high Fib DiD labelling, 10 simulated SAOS-2 cell acquired between 35% and 47% of their volume from simulated Fib. Proportional volume transfers to Fib were generally lower than for SAOS-2 (Fig. 8; Supplemental Information Table S1).

**FIGURE 8.**
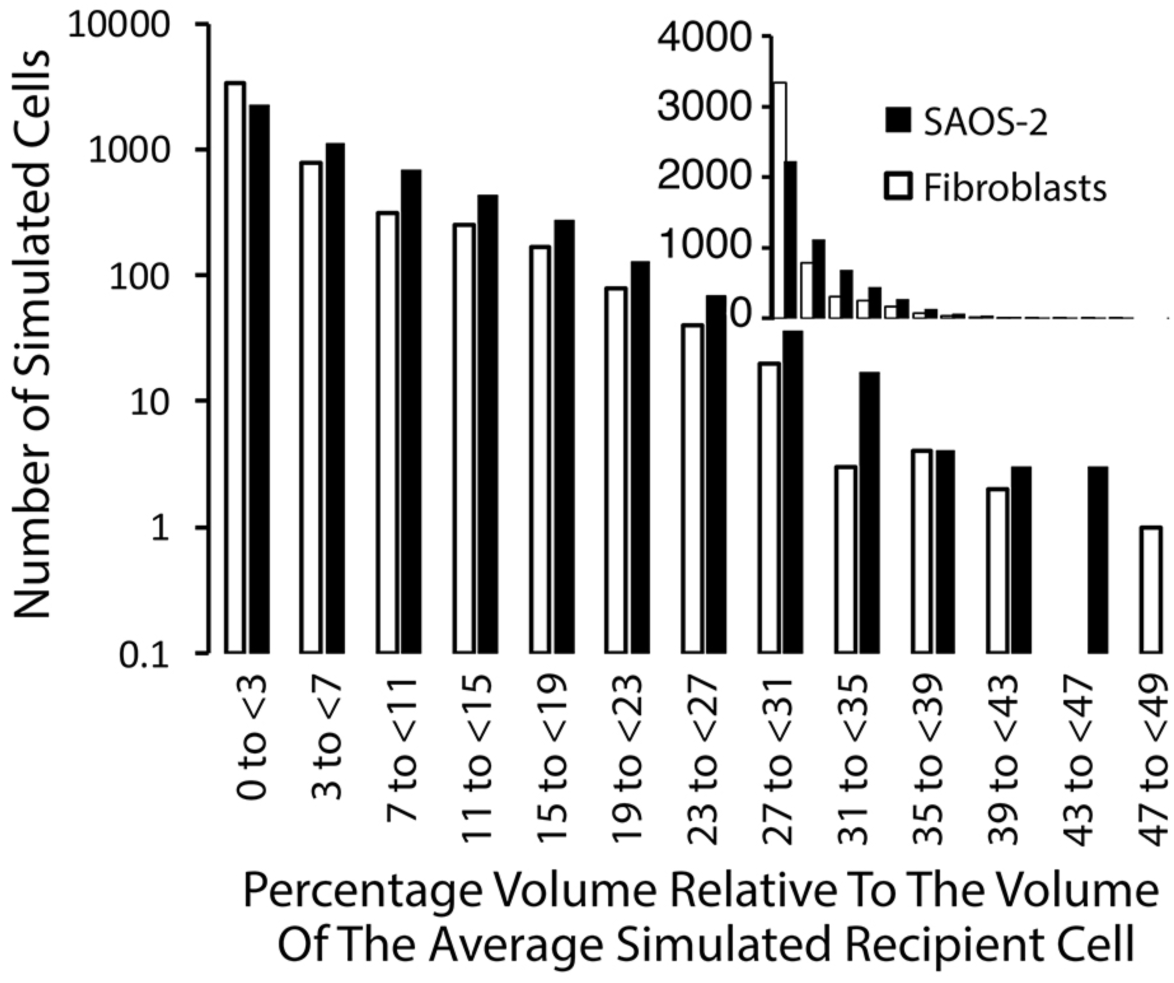
Histogram illustrating the number out of 5000 simulated recipient SAOS-2 or Fib, according to the amount of simulated CPP cytoplasm uptake, expressed as percentage relative to the average recipient cell volume. Simulated SAOS-2 accepted proportionately more of their cellular volume from Fib compared with transfer from SAOS-2 to Fib. Data is provided on both log and linear scales to aid visualization. Further detail is available in Table S1 of Supplemental Information.

In as much as variability in donor cell fluorescence undermines direct estimation of volume transfers by CPP, the same applies for estimation of volume exchanges via shed vesicles and TNT. Nonetheless, it is not plausible that exosomes or other shed vesicles could account for the large volume transfers calculated as due to CPP in the current study. Similarly, while TNT are reported to transfer organelles, time-lapse recordings reveal such transfers to be relatively few by comparison with CPP. This is consistent with the absence of a clear mechanical motive force in TNT, as opposed to the active pumping mechanism of CPP. As such, the current simulations appear to offer the first estimation of cytoplasmic volume transfers between mammalian cells.

### Limitations of microscopy, modelling and simulations

Although CLSM and holotomographic data were consistent with each other and our interpretation of CPP, further more clear visualization of individual CPP events requires additional work beyond the immediate scope of the current paper. Discrete punctate labelling of organelles aided observation of potentially small inter-cellular exchanges, but use of this as a proxy marker for cytoplasmic flow does generate some uncertainty for quantitation. A preference of CPP for the transfer of smaller as opposed to larger organelles seems likely. This appears consistent with experimental observations, and suggests size dependent selection for CPP transfer of organellar sub-populations, and this requires further study.

It is possible that fluorescent lipophilic label in the plasma membranes of donor cells contributed to the fluorescence transfers observed. However, given intense organellar labelling compared with mostly undetectable label in plasma membranes, as well as evident transfer of organelles in CLSM time-lapse recordings, we suggest that fluorescence acquisition by transfer of plasma membranes would be negligible compared with that of organelles acquired in transferred cytoplasm.

Some aspects of our modeling bear further discussion. Turbulent flow is near impossible in radius values used, supporting use of the Hagen-Poiseuille relationships. Cytoplasmic viscosity is non-uniform and dependent on scale. Viscosity is low but varies across micro-volumes of the cell, dependent on contents. When measured at the whole cell level, viscosity is high due to the admixture of organelles and cytoskeletal elements (27, 28, 35). At the scale here modeled, cytoplasmic viscosity ranges upwards from that close to water to 4 mPa.s (27, 28). The effect of organelles suspended in cytoplasm is difficult to anticipate. While cytoplasm itself at the scale studied may have low viscosity in the order of 1.5 mPa.s, we have made reasonable accommodation for the effect of organelles by including higher viscosity values in simulations.

Preliminary simulations applying a normal distribution for variables other than stiffness and fluorescence, generated ‘a central hump’ in fluorescence profiles inconsistent with experimental fluorescence. Using a distribution with a flattened profile achieved simulation outcomes more similar to experimental results, suggesting a uniform distribution for key variables *in-vivo*. Confirmation awaits improved structural and temporal resolution of events in living cells. Modest divergence of simulated from experimental results, likely reflects limitations inherent to the model, including possible skewedness and unknown dependencies between variables.

## CONCLUSIONS

The observed relationships between cell stiffness and fluorescence transfer, would not be expected if TNT or shed vesicles played a significant role, and this further supports our interpretation of CPP from time-lapse observations. Taken together, data support our CPP hypothesis as what seems to be a previously unrecognized mechanism for inter-cellular cytoplasmic exchange, and this report forms a reasonable theoretical framework to further investigate this hypothesis. CPP is in some ways similar to the hydrodynamic mechanisms described for formation of lamellipodia (36), blebbing (37, 38), and the formation of lobopodia (39).

We speculate that CPP contributes additionally to a variety of otherwise described processes, including transfer of melanosomes and mitochondria (2, 4, 5, 20, 29-31), and the development of cancer associated Fib (40).

Our earlier work showed CPP causes significant phenotypic change, including altered morphology and cytokine synthesis of recipient cells (1, 18). The inflammatory cytokine Tumor Necrosis Factor-α increased transfer from fibroblasts to SAOS-2, and this seemed due to increased binding of SAOS-2 via ICAM-1 (1, 25). One limitation of those earlier reports, was that we only studied cell phenotype in co-culture, and not cells separated after co-culture according to the extent of CPP uptake (1, 18). This is addressed in more recent work (19), in which we study MC sub-populations separated by fluorescence activated cell sorting on basis of the level of fibroblast label uptake. We found that acceptance of fibroblast marker increased MC migration and cell size (19). Internal complexity was also increased, as expected from acquisition of additional organelles from fibroblasts (19). Because CPP generates sub-populations of MC with altered morphology, we suggest CPP might contribute to MC morphological diversity *in-vivo*, and hence to histopathological pleomorphism relevant to cancer diagnosis and prognosis (41). Also, since MC migration is important in cancer invasion and metastasis (41), we suggest CPP contributes to these aspects of cancer. MC diversity is central to cancer progression (41), so CPP driven MC diversity may play a role (1, 18, 19).

CPP seems mechanically more akin to intercellular exchange via TNT than via exosomes, because micro-fusions establish physical cytoplasmic continuity of neighboring cells in both TNT and CPP exchange, while exosome transfer does not require cell to cell contact. Despite this, the biological response of MC to CPP transfer of fibroblast cytoplasm in our separate work (1, 18, 19), seems more similar to the published response of cells to uptake of exosomes, than to TNT mediated transfer. For example, exosome uptake alters cell morphology (14, 15, 42), but we find no literature of TNT mediating this effect. Similarly, while exosomes from a variety of sources can increase migration of several cell types (16, 43-46), there is less evidence for a similar effect for cellular contents transferred via TNT (17). Seemingly different effects of cytoplasmic transfer by CPP and TNT, underscore the distinction between the two processes.

We have now reported CPP transfer with various melanoma, ovarian cancer, lung cancer and osteosarcoma cell lines (1, 19), while preliminary experiments suggest this also occurs between other cell types including: endothelium, smooth muscle cells and pulmonary basal cells. Cancer rarely generates new biology, but instead usually perverts established mechanisms. From this, and given altered phenotype following cytoplasmic transfer (1-17, 19, 20), we suggest CPP may contribute to cell differentiation and phenotypic control in other biological settings including: embryogenesis, development, inflammation and wound healing.

## Supporting information

Matlab Files

## AUTHOR CONTRIBUTIONS

HZ conceived and conducted all experiments and modelling and prepared the manuscript, with exception of holotomography conducted by BC. EK assisted with cell culture. NP and KM assisted with AFM, while YR, VB, SF and KM assisted with CLSM and fluorescence quantitation. GR and JC gave key input for mathematical and MATLAB modelling respectively. GWL assisted preparing movies. All authors and MASM contributed to critical analysis and manuscript preparation. The authors declare no conflict of interest.

## ACKNOWLEDGEMENTS

We thank the Memorial Sloan Kettering Cancer Center, including via MSKCC P30 CA008748 Cancer Center Support Grant, as well as the Australian Dental Research Fund for their support of this work. We also thank an anonymous donor for their kind contribution. We also thank Dr R Norden of the Graduate School of Biomedical Engineering, University of NSW for his advice.

