## Supplemental Information Text for "Potential hydrodynamic cytoplasmic transfer between mammalian cells: Cell-projection pumping"

### CONTENTS


#### Contents
***Pages 2 to 3***

#### Detailed materials and methods
***Pages 4 to 7***

#### Detailed derivation of the mathematical model
***Pages 8 to 14***

#### References
***Page 15***

#### Supplemental figures
***Page 16:*** **Fig. S1** Histograms for experimental data for stiffness, DiD fluorescence and DiO fluorescence in co-cultured SAOS-2 and fibroblasts, as well as stair plots for the ECDF of these data with superimposed linear smoothing, and histograms for 5000 simulated cells generated by the ECDFs shown.

***Page 17:*** **Fig. S2** Histogram output of MATLAB script for simulation of CPP transfer from fibroblasts to SAOS-2 comparable to that seen by experiment *(Fig. 7, Table 1 of the main manuscript, Table S1 Supplemental Information).*

***Page 18:*** **Fig. S3** Histogram output of MATLAB script for simulation of CPP transfer from SAOS-2 to fibroblasts comparable to that seen by experiment *(Fig. 7, Table 1 of the main manuscript, Table S1 Supplemental Information).*

***Page 19:*** **Fig. S4** Consecutive confocal optical sections taken from Supplemental Movie S2 at the time points shown in Fig. 2c,d of the main manuscript.

***Page 20:*** Fig. **S5** Holotomographic images at single optical levels of a co-culture of fibroblasts co-cultured with SAOS-2 showing the presence of branching cell-projections consistent with those inferred from CLSM observations.

***Page 21:*** **Fig. S6** Sequential holotomographic time-lapse images at a single optical level of a co-culture of fibroblasts with SAOS-2, showing rapid movement of a cell projections between time-points. Several separate cell projections were evident at the interface between two adjacent cells.

***Page 22:*** **Fig. S7** Flattened optical confocal microscopy projection and two optical levels of fibroblasts co-cultured for 24 h with MM200-B12, showing the presence of organelles derived from different cells on different optical levels.

***Page 23:*** **Fig. S8** Flattened optical confocal microscopy projections of fibroblasts co-cultured for 24 Hr with MM200-B12, showing tunneling nanotubes.

***Page 24:*** **Fig. S9** Holotomography demonstrating tunneling nanotubes in a co-culture of fibroblasts with SAOS-2 at a single time point and four separate optical levels.

***Page 25:*** **Fig. S10** Flattened optical confocal microscopy projection and 6 optical levels of fibroblasts co-cultured with MM200-B12, showing close entwining of fibroblast processes with malignant cells in absence of non-specific transfer of fibroblast label.

***Page 26:*** **Fig. S11** Scattergrams from computer simulations showing the relationship between stiffness of individual receptor cells and fluorescence acquired from donor cells, where stiffness of the two cell populations is varied, as well as the median CPP volume exchange for each simulation.

#### Legends for movies
***Pages 27 to 29:*** Movies S1, S2, S3, S4, S5, S6

### Supplemental Table

***Page 30:*** Table S1

### MATLAB script

***Pages 31 to 33:*** MATLAB script for generating simulated cell values from input data when the lowest value in input data is significantly greater than zero.

***Pages 34 to 37:*** MATLAB script for generating simulated cell values from input data when the lowest value in input data does not approach zero.

***Pages 37 to 47:*** MATLAB script for simulating cytoplasmic exchange between populations of cells via cell-projection pumping.

### List of additional files provided as Supplemental Information

***Movies:***

- **S1:** CPP transfer from a Fib to a MM200 B12 cell
- **S2:** CPP transfer from a Fib to a MM200 B12 cell
- **S3:** CPP transfer from a MM200B12 cell to a fibroblast
- **S4:** Deep probing and indentation of a MM200B12 cell by fibroblast cell projections
- **S5:** Rotating image of fibroblast deeply probing a MM200 B12 cell
- **S6:** Holotomography time-lapse consistent with CPP

***MATLAB scripts:***

- **ScriptECDF.m:** Script for generating simulated values for cells by ECDF
- **ScriptECDFLowValues.m:** Script for generating simulated values for cells by ECDF when some input values are very low
- **ScriptCPP.m:** Script for simulating CPP

***Simulated Fibroblast and SAOS-2 MATLAB input data files***

- **FibGreen500.mat**
- **FibRed5100.mat**
- **FibStiff500.mat**
- **FibStiff5100.mat**
- **SAOSGreen5100.mat**
- **SAOSRed5000.mat**
- **SAOSStiff5000.mat**
- **SAOSStiff5100.mat**

Please note that the CPP script requires that there are always more simulated Donor Cells (Cells A), than there are simulated Receptor Cells (Cells B). For this reason, donor cell input files are labelled xx5100 with 5100 simulated cells, while receptor cell input files are labelled xx5000 with 5000 simulated cells.

***MATLAB figures:***

- **SimFibToSAOS.fig:** Figure generated from simulated CPP transfer from Fib to SAOS-2
- **SimSAOStoFib.fig:** Figure generated from simulated CPP transfer from SAOS-2 to Fib

### DETAILED MATERIALS AND METHODS

#### Materials

For experiments by confocal laser scanning microscopy (CLSM) and atomic force microscopy (AFM), culture media including M199, DMEM-α Trypsin (0.25%)-EDTA (0.02%) and phosphate buffered saline (PBS), as well as Penicillin (10,000 U/ml)-Streptomycin (10,000 μg/ml) concentrate solution were prepared and supplied by the Memorial Sloan-Kettering Cancer Centre Culture Media Core Facility (New York, NY). Amphoteracin B was purchased from Life Technologies (Grand Island, NY). Gelatin was from TJ Baker Inc (Philipsburgh, NJ). Bovine serum albumin was from Gemini Bioproducts (West Sacramento, CA). For holotomography, culture media were from (Sigma-Aldrich, St. Louis, USA), while bovine calf serum (BCS) was from Bovogen (VIC, Australia) and antibiotics were from CSL Biosciences, (VIC, Australia). Falcon tissue culture flasks, Atomic Force Microscopy (AFM) dishes and centrifuge tubes were purchased from BDBiosciences (Two Oak Park, Bedford, MA). Culture well coverslips were from Lab-Tek (Rochester, NY). Glass bottom dishes for holotomography were from Ibidi (Graefeling, Germany). Human dermal fibroblasts (HDF) were from The Coriell Institute (Camden, NJ). SAOS-2 osteosarcoma cells were from the American Type Culture Collection (VA, USA). MM200-B12 melanoma cells from The Millennium Institute (Westmead, NSW, Australia). The fluorescent labels 1,1'-dioctadecyl-3,3,3',3'-tetramethylindodicarbocyanine perchlorate (DiD), 3,3'-dioctadecyloxacarbocyanine perchlorate (DiO), and Bacmam 2.0 Cell lights Nuclear-GFP baculovirus, were purchased from Molecular Probes by Life Technologies (Grand Island, NY) in the form of DiD and DiO Vybrant cell labelling solutions, and BacMam Cell Light transfection reagent. Paraformaldehyde (PFA) solution (32%) was purchased from Electron Microscopy Supplies (Hatfield, PA). A 6.1 μm spherical polystyrene AFM probe was purchased from NanoAndMore (Lady's Island, SC). The anti-fade reagent used was supplied by the Molecular Cytology core facility at Memorial Sloan Kettering Cancer Center.

#### Cell culture and fluorescent labelling

Cell culture was as earlier described (1-4). HDF were cultured on gelatin coated surfaces (0.1% in PBS) in DMEM-α (15% FCS). Malignant cell (MC) lines were: melanoma MM200-B12 cultured in DMEM-α (10% FCS); and osteosarcoma cells SAOS-2 in M199 (10% FCS).

Labelling solutions of DiD for fibroblasts (1mM) and DiO for MC (2mM) were applied to cells for 30 min in the case of DiD, and 1h for DiO. Monolayers were washed prior to overnight culture and further washing before co-culture. In some experiments, MM200-B12 were transfected with green fluorescent protein (GFP) expressing baculovirus.

#### Co-culture conditions

Co-cultures were on gelatin (0.1% in PBS) coated surfaces with Fibroblasts seeded from 1 to 2 x $10^4$ cells per $cm^2$ into either $25cm^2$ AFM culture plates (4), culture well coverslips for CLSM, or 35mm glass bottom dishes for holotomography, and allowed to adhere overnight before labelling. MC were seeded at near confluence in either 25 $cm^2$ flasks or 6 well culture plates prior to labeling. MC were then harvested with trypsin-EDTA and seeded over fibroblasts in DMEM-α with BSA (4%) at 4 x $10^4$ cells per $cm^2$ for up to 24 h co-culture for CLSM and AFM experiments, and up to 55 h for holotomography.

#### Time-lapse CLSM

Eight separate visual fields of fibroblasts co-cultured with GFP labelled MM200-B12 were recorded for 25 h at 3 min intervals, representing 1.13 $mm^2$ culture surface area. Nine further separate visual fields of DiO pre-labelled MM200-B12 were recorded for 8 h 15 min at 5 min intervals and at slightly higher magnification, representing 0.76 $mm^2$ culture surface area. Monolayers were fixed with paraformaldehyde after co-culture. CLSM was by a Zeiss LSM 5Live line-scanning confocal microscope.

#### Holotomography

Holotomography time-lapse images were collected using a Nano Live 3D Cell Explorer (Lausane, Switzerland) under normal incubation conditions (37°C, 5% $CO_2$, 99% humidity). Recordings were of culture surface areas measuring 6.4 x $10^{-3}$ $mm^2$. 96 optical levels 'Z-positions' were obtained every 2 min to generate 4 separate recordings for: 6 h 10 min; 6 h 38 min; 18 h 58 min; and 19 h 24 min. Following time-lapse imaging, raw Stack TIFF files were then exported from software STEVE (V2.1) and ImageJ (V 1.8.0) was used to combine Stack TIFF files to create TIFF movies for subsequent analysis.

#### Combined atomic force and fluorescence microscopy

Paraformaldehyde fixed monolayers were stored in PBS at 4° C for combined fluorescence-AFM recordings of randomly selected cells (4). An Asylum Research MFP-3D-BIO atomic force microscope coupled with a Zeis Axio Observer A1 fluorescence microscope was used. Bright field and fluorescence images for both DiO and DiD channels were recorded prior to AFM scanning. A 1 μm AFM spherical polystyrene probe was used to record 16 x 16 points of force curves over 50 μm x 50 μm areas. Asylum Research, Software Version IX Young's modulus for each point by the Hertz model (4, 5). Height maps, bright field and fluorescence images were compared to localize discrete AFM measurement points to individual cells, and stiffness fingerprints were prepared (4).

### Morphometric analysis and cell stiffness analysis

ImageJ open source software (http://imagej.net/Contributors) was used to segment and analyze fluorescence images of individual SAOS-2 and fibroblasts. Cell surface profile area was determined, while both Red and green fluorescence was summated for each cell. Fluorescence intensity in both fluorescence channels was expressed in 'Fluorescence Units' (summated fluorescence / surface profile area). Both fibroblasts and SAOS-2 were designated as belonging to one of two groups, being 'high' or 'low' labelling from the opposing cell type. Median AFM stiffness was determined for individual cells, while stiffness fingerprints were also made of cells according to group to address sampling limitations as earlier described (4). Prism 6.0e software (GraphPad Software Inc, La Jolla, CA) was used for statistical analysis.

### Computer simulation of cytoplasmic and fluorescence transfer between fibroblast and SAOS-2 populations by CPP

Estimated cumulative distribution functions (ECDF) were developed in MATLAB (MATLAB by MathWorks Inc) from experimental median cell stiffness and fluorescence data. All MATLAB scripts are provided below. ECDFs were then used to generate simulated populations of cells with distributions for stiffness and fluorescence closely approximating those of experimental data. This method was used to generate 5100 donor fibroblasts, 5100 donor SAOS-2, 5000 recipient SAOS-2, and 5000 recipient fibroblasts (Fig. S1).

Co-culture simulations were in MATLAB of random interactions between simulated donor and receptor cells, making random selection of values from lists of variables used to calculate CPP. Values loaded into these lists had distributions bounded by target minimum and maximum values, while target minima and maxima were established at the start of each simulation. Variables modelled in this way were: the number of Donor Cells A each Receptor Cell B could interact with; the number of transfer events each Receptor Cell B could have with each Donor Cell A; the flow rate *(U)* for each transfer event; the length at time 0 $(L_0)$ of each cell-projection; the radius *(r)* of each cell-projection; and the viscosity of cytoplasm $(\eta)$.Values for these parameters, were inferred on basis of CLSM observations, with exception of viscosity, which was taken from the literature (6, 7). The only exception to this was for the time permitted for each transfer event, the maximum of which is defined by $L_0/U$, and random choice of time was made from a pre-determined proportionate range between 0 and $L_0/U$. MATLAB script for simulations is provided below.

Average SAOS-2 and fibroblast cell height was determined from AFM data (3.89 x $10^{-6}$m and 2.36 x $10^{-6}$m respectively), while average SAOS-2 and fibroblast cell surface area (1.53 x 10-

$^{9}m^2$ and 5.34 $x10^{-9}m^2$ respectively) was by image analysis from separate experiments, and these data were used to calculate fluorescence from volume transfers as detailed below.

Volume and fluorescence transfers for each simulated cell pairing were determined, and summated simulation results compared with experimental results. Maximal pressure generated during individual simulated CPP events was also recorded. Distributions of input variables as well as simulation outcomes were plotted in histograms (Figs. S2, S3).  Data were analyzed using PRISM 7 (GraphPad  Software Inc), and Mann Whitney U Tests where appropriate.

### DETAILED DERIVATION OF THE MATHEMATICAL MODEL

#### General features of the model

Fig. 4a of the main manuscript illustrates the CPP hypothesis in a cartoon of cells interacting. Fig. 4b of the main manuscript shows the simple two chamber system used to model CPP mathematically, relating a constant rate of retraction of the cell projection *(U),* to pressure $(\Delta P_A$ and $\Delta P_B)$ for both chambers from fluid escaping the shortening cell-projection.

Only flow into Receptor Cell B $(Q_B)$ was calculated in the current study, since it is only $Q_B$ which will deliver cytoplasm to the opposing cell (Fig. 4b of the main manuscript). Also, since maximum pressure is located within of the retracting cell-projection at position O (described below, Fig. 4ciii of the main manuscript), there is no actual cytoplasmic flow at O which acts as an effective 'syringe plunger' for flow in both cell directions. $\Delta P$ is exhausted upon complete retraction, thus limiting the total possible volume of flow to that of the cell-projection.

#### Modification of the Hagen-Poiseuille Equation accommodating differences in cell stiffness

Mathematical assumptions made are detailed in the main manuscript.

Calculations were based on the well described Hagen-Poiseuille relationships, where resistance to flow per unit length in a cylindrical tube $(\rho)$ is given by Eq. 1, and the flow rate $(Q)$ of a Newtonian fluid of viscosity $(\eta)$ through a cylindrical tube of length $(L)$ with radius $(r),$ due to a pressure difference $(\Delta P)$ is as per Eq. 2 (8).

$$\rho = \frac{8\,\eta}{\pi \cdot r^{\,4}} \tag{1}$$

$$Q = \frac{\Delta P}{\rho \cdot L} = \frac{\Delta P \cdot \pi \cdot r^{\,4}}{8\,\eta \cdot L} \tag{2}$$

Distribution of fluid during retraction of the cell-projection in Fig. 4a,b of the main manuscript is influenced by the relationship between stiffness of Donor Cell A $(S_A)$ and Receptor Cell B $(S_B)$, and this requires modification of the relationships described in Eqs. 1 and 2 for calculation of that portion of total flow distributed to Receptor Cell B $(Q_B)$.

Fig. 4ci of the main manuscript shows a cylindrical tube of known length $L(t)$ at time $(t)$ as measured from '0' at it's origin which is closed and marked to the left, and in which there is contraction at a constant rate $(U),$ as indicated in Eq. 3.

$$U = \frac{dL(t)}{dt} \tag{3}$$

The tube contains a Newtonian fluid, and is divided into n cylinders of equivalent length $(\Delta x)$ as shown in Eq. 4, indexed from $n$=1 to n (Fig. 3cii of the main manuscript).

$$\Delta x = \frac{L(t)}{n} \quad (4)$$

Because the tube undergoes a constant contraction, each cylinder also contracts by *UΔt/n* to displace a volume of fluid *(ΔV)* as in Eq. 5, where *Ca* is the cross-sectional area of the tube.

$$\Delta V = \frac{Ca \cdot U(\Delta t)}{n} \quad (5)$$

In this way, each cylinder donates an equivalent volume *(ΔV)* and flow rate increment *(Δq = ΔV/Δt),* to the total flow rate of the cylinder as in Eq. 6, substituting for *n* from Eq. 4.

$$\Delta q = \frac{Ca \cdot U}{n} = \frac{Ca \cdot U \cdot \Delta x}{L(t)} \quad (6)$$

Rearrangement of Eq. 6 gives Eq. 7.

$$\frac{\Delta q}{\Delta x} = \frac{Ca \cdot U}{L(t)} \quad (7)$$

As $n \to \infty$, $\Delta x \to 0$, so that the flow rate at *x*, *(Q(x)),* is given by Eq. 8.

$$Q(x) = \int_0^x \frac{Ca \cdot U}{L(t)} dx = \frac{Ca \cdot U \cdot x}{L(t)} \quad (8)$$

Note that the flow rate at the end of the tube where *x=L(t)* is *CaU* as expected (Fig. 4cii of the main manuscript).

Let $\rho$ be the resistance per unit length as given in Eq. 1, so that the resistance offered by any given small cylinder comprising the cell-projection *(ΔR)* is given by Eq. 9. The pressure drop across the cylinder *($\Delta P_{\Delta X}$)* is given by Eq. 10 which is obtained by rearranging Eq. 2 and substituting $\Delta P_{\Delta x}$ for *ΔP, Δx* for *L* and *Δq* for *Q*. Then using Eq. 6 for *Δq* results in final expression in Eq. 10.

$$\Delta R = \rho \cdot \Delta x \quad (9)$$

$$\Delta P_{\Delta X} = \Delta q \cdot \Delta R = \frac{Ca \cdot U}{n} \cdot \rho \cdot \Delta x \quad (10)$$

The pressure drop at any given cylinder *k* and *L=x* is given by Eq. 11, which can be rearranged to Eq. 12.

$$\Delta P_X = \frac{k \cdot Ca \cdot U}{n} \cdot \rho \cdot \Delta x \quad (11)$$

$$\frac{\Delta P_X}{\Delta x} = \frac{k \cdot Ca \cdot U \cdot \rho}{n} \quad (12)$$

Substituting for *1/n* from Eq. 4, and recognizing that *x= kΔx* gives Eq. 13.

$$\frac{\Delta P}{\Delta x} = \frac{Ca \cdot U \cdot \rho \cdot k \cdot \Delta x}{L(t)} = \frac{Ca \cdot U \cdot \rho \cdot x}{L(t)} \quad (13)$$

Allowing $\Delta x \to 0$, gives the expressions in Eq. 14.

$$\frac{\Delta P}{\Delta x} \to \frac{dP}{dx} = \frac{Ca \cdot U \cdot \rho \cdot x}{L(t)} \quad (14)$$

From this, the pressure drop *P(x)* at *x*, is given as expressions in Eq. 15.

$$P(x) = \int_0^x \frac{Ca \cdot U \cdot \rho \cdot x}{L(t)} dx = \frac{Ca \cdot U \cdot \rho \cdot x^2}{2 \cdot L(t)} \quad (15)$$

Consider the above outlined system now replicated, and one of these elements to be rotated so that the two tubes now abut end to end, with the origin $x$=0 being identical for both. This now represents a cell-projection joining Donor Cell A 'to the left', with Receptor Cell B positioned 'to the right'. The origin represents a point in the cell-projection (O) where during contraction of the cell-projection, there is maximum pressure and no flow, the origin functioning as an effective 'syringe stop' for flow in both directions. The length of tube between the origin and Donor Cell A is $L_A$, and that to Receptor Cell B is $L_B$ (Fig. 4ciii of the main manuscript), giving Eq. 16 for length at time $t$.

$$L(t) = L_A(t) + L_B(t) \tag{16}$$

The tube is open to Cells A and B, and flow out of the tube in both directions is resisted by constant reaction pressures $F_A$ and $F_B$ in Cells A and B respectively (Fig. 4ciii of the main manuscript). These reaction pressures are equal to the yield points $P_A{}^Y$ and $P_B{}^Y$ which are proportional but not identical to the measured median cell stiffness of the Donor and Receptor cells ($S_A$ and $S_B$), so that it may be helpful to read '*S*' for '*F*' when making reference to Fig. 4b,c of the main manuscript.

Noting that $U$ is constant, Eq. 16 gives Eq. 17 following simplification, where: $U = L/t$, $U_A=L_A/t$, and $U_B=L_B/t$.

$$U = U_A + U_B \tag{17}$$

The relationships in Eq. 18 follow from the above.

$$\frac{U_A}{U} = \frac{L_A(t)}{L(t)} \qquad \frac{U_B}{U} = \frac{L_B(t)}{L(t)} \tag{18}$$

If $P_{max}(t)$ is the pressure at the origin (O), then from Eq. 15, the pressure at $x_B$ going to the right *($P_R$)* is given by Eq. 19, ultimately reaching and being balanced by the hydrodynamic force resisting flow by Cell B *($F_B$)* to the right, with pressure at $x_A$ going left *($P_L$)* reaching the hydrodynamic force resisting flow by Cell A *($F_A$)* to the left (Eq. 19), as illustrated in Fig. 4ciii of the main manuscript.

$$P_R(x) = P_{max}(t) - \frac{Ca \cdot U_B \cdot \rho \cdot x_B^2}{2 \cdot L_B(t)} \qquad P_L(x) = P_{max}(t) - \frac{Ca \cdot U_A \cdot \rho \cdot x_A^2}{2 \cdot L_A(t)} \tag{19}$$

Since $L_A = x_A$, and $L_B = x_B$, Eq. 19 simplify to Eq. 20.

$$P_R(x) = P_{max}(t) - \frac{Ca \cdot U_B \cdot \rho \cdot L_B(t)}{2} = F_B \qquad P_L(x) = P_{max}(t) - \frac{Ca \cdot U_A \cdot \rho \cdot L_A(t)}{2} = F_A \tag{20}$$

Rearrangement of Eq. 20 gives Eq. 21.

$$F_A + \frac{Ca \cdot U_A \cdot \rho \cdot L_A(t)}{2} = F_B + \frac{Ca \cdot U_B \cdot \rho \cdot L_B(t)}{2} \tag{21}$$

Substituting for $L_A$ from Eq. 16 gives Eq. 22.

$$F_A + \frac{Ca \cdot U_A \cdot \rho \cdot (L(t) - L_B(t))}{2} = F_B + \frac{Ca \cdot U_B \cdot \rho \cdot L_B(t)}{2} \tag{22}$$

Substitution for: $U_B$ from Eq. 17; $U_A/U$ from Eq. 18; and $L_A$ from Eq. 16, gives Eq. 23 for $L_B$.

$$L_B(t) = \frac{F_A - F_B}{Ca \cdot U \cdot \rho} + \frac{L(t)}{2} \quad (23)$$

The algebraic relationships leading to Eq. 23 apply equally to generate Eq. 24.

$$L_A(t) = \frac{F_B - F_A}{Ca \cdot U \cdot \rho} + \frac{L(t)}{2} \quad (24)$$

From Eqs. 23 and 24, when $F_A = F_B$, then $L_A(t) = L_B(t) = L(t)/2$, and both cells receive equivalent flow as expected from symmetry of the system. When $F_A > F_B$, a time is reached when $L_A(t)$ reaches 0 and all remaining flow is to the right, and Receptor Cell B receives more flow than Cell Donor Cell A (Fig. 4ciii of the main manuscript), while the reverse applies when $F_A < F_B$.

Note that in Eq. 20 $P_{max}(t)$ decreases with time so where $F_A > F_B$, $P_{max}(t)$ reaches the yield point of cell A ($P_A^Y$) when $L_A = 0$, and no further flow into Cell A occurs; while where $F_A < F_B$, $P_{max}(t)$ reaches the yield point of cell B ($P_B^Y$) when $L_B = 0$, and no further flow into Cell B occurs.

Equation 25 follows from Eq. 8.

$$Q_B = \frac{Ca \cdot U \cdot L_B(t)}{L(t)} \quad (25)$$

To establish the total flow transferred to Receptor Cell B *($Q_B(t)$)*, $L_B(t)$ from Eq. 23 is substituted into Eq. 25, which with simplification gives Eq. 26.

$$Q_B = \frac{F_A - F_B}{\rho \cdot L(t)} + \frac{Ca \cdot U}{2} \quad (26)$$

Because $U$ is constant, $L(t)$ is given by Eq. 27, and substitution for $L(t)$ in Eq. 26 gives Eq. (28).

$$L(t) = L_0 - U \cdot t \quad (27)$$

$$Q_B = \frac{F_A - F_B}{\rho \cdot (L_0 - U \cdot t)} + \frac{Ca \cdot U}{2} \quad (28)$$

Note that Eq. 28 can only apply while $(F_A - F_B)/\rho(L_0 - Ut) \leq CaU/2$, because once $(F_A - F_B)/\rho(L_0 - Ut) = CaU/2$, $L_A = 0$ and $L_B = L$ where $S_A > S_B$, with all remaining flow being to the right into Receptor Cell B at a rate of $Q_B = CaU$. We define the time at which this occurs as time $tc$, which is given by Eq. 29. Please note that where $S_A < S_B$, $L_B = 0$ and $L_A = L$ and all remaining flow is to the left into Donor Cell A after time $tc$.

$$\frac{F_A - F_B}{\rho \cdot (L_0 - U \cdot tc)} = \frac{Ca \cdot U}{2} \quad (29)$$

Expansion, rearrangement and simplification of Eq. 29, gives Eq. 30 for $tc$, where the absolute value of second term on the right is required to accommodate occasions when $F_A < F_B$.

$$tc = \frac{L_0}{U} - \left| \frac{2(F_A - F_B)}{Ca.\rho \cdot U^2} \right| \quad (30)$$

Fig. 4d of the main manuscript is a graphical representation of $Q_B$ from $t = 0$ to $t = L_0/U$ at which time $L_B = 0$ and no further flow $Q_B$ is possible. When $F_A > F_B$, Eq. 28 for $Q_B$ applies for $t \leq tc$, and $Q_B = CaU$ for $tc < t < L_0/U$. Increasing values of $(F_A - F_B)$ reduce $tc$, while as $(F_A - F_B)$ approaches 0, $tc$ approaches $L_o/U$ and $Q_B$ approximates $CaU/2$ from above for increasing time when

$F_A > F_B$, and from below when $F_A < F_B$. Also, when $F_A < F_B$, Eq. 28 predicts that $Q_B$ approaches 0 as $t$ approaches $tc$, after which $Q_B$ remains 0 due to exhaustion of $L_B$ to 0 at $tc$.

#### *Calculation of volumes transferred*

Volumes transferred can be calculated by integration of curves for $Q_B$ such as show in Fig. 4d of the main manuscript. Let the total volume transferred to Receptor Cell B be $V_B$.

Where $F_A > F_B$, the relationships outlined above determine $V_B(t)$ as per Eq. 31, where the first term relates to those parts of the curves in Fig. 4d of main manuscript where $Q_B$ is rising, and the second term relates to the following horizontal parts of curves once $Q_B$ reaches CaU.

$$V_B(t) \;=\; \int_0^{tc} \left(\frac{Ca \cdot U}{2} + \frac{F_A - F_B}{\rho \cdot (L_0 - U \cdot t)}\right) dt \;+\; \int_{tc}^{L_0/U} (Ca \cdot U)\; dt \tag{31}$$

Similarly, where $F_A < F_B$, $V_B(t)$ is given by Eq. 32, where all curves illustrated in Fig. 4d of the main manuscript reduce towards $Q_B = 0$, and only one term is required for integration.

$$V_B(t) \;=\; \int_0^{tc} \left(\frac{Ca \cdot U}{2} + \frac{F_A - F_B}{\rho \cdot (L_0 - U \cdot t)}\right)\; dt \tag{32}$$

Also, where $F_A = F_B$, $V_B(t)$ is given by Eq. 33.

$$V_B(t) \;=\; \int_0^{L_0/U} \left(\frac{Ca \cdot U}{2}\right)\; dt \tag{33}$$

To aid integration in Eqs. 31 and 32, define $w(t)$ as in Eq. 34, such that when $t = 0$, $w = L_0$; and when $t = tc$, $w = L_0\text{-}Utc$.

$$w(t) \;=\; L_0 - U \cdot t \tag{34}$$

Differentiating Eq. 34 gives Eq. 35.

$$dw \;=\; -U \cdot dt \tag{35}$$

From this, the second term in Eqs. 31 and 32 can be expressed and integrated with regard to $w$ as in Eq. 36, where ln$(w)$ is the natural logarithm of $w$. Substitution of Eq. 34 into Eq. 36 gives the expression in Eq. 37.

$$-\frac{1}{U} \int_{L_0}^{(L_0 - U \cdot tc)} \left(\frac{F_A - F_B}{\rho \cdot w}\right) dw \;=\; -\frac{F_A - F_B}{\rho \cdot U} \cdot \ln(w)\, \Big|_{L_0}^{L_0 - tc \cdot U} \tag{36}$$

$$= -\frac{F_A - F_B}{\rho \cdot U} \cdot \begin{cases} \ln\left(\frac{L_0 - U \cdot t}{L_0}\right), & if\;\; t \leq tc \\ \ln\left(\frac{L_0 - U \cdot tc}{L_0}\right), & if\;\; t > tc \end{cases} \tag{37}$$

From Eq. 37, the integral of $V_B(t)$ for $0 < t < tc$ in Eqs. 31 and 32 is as in Eq. 38.

$$\frac{Ca \cdot U \cdot t}{2} \;-\; \frac{F_A - F_B}{\rho \cdot U} \cdot \ln\left(\frac{L_0 - U \cdot t}{L_0}\right) \tag{38}$$

Combining the result of Eq. 38 with Eqs. 31, 32, and 33 provides Eqs. 39 to 42 used to determine $V_B$ in computer simulations for all conditions tested in the current study, where: $tm$ is the time at which any given contraction event ceases; and $Z$ is a constant correcting for the assumed linear relationship between median cell stiffness and both $F_A$ and $F_B$, such that $ZS_A=F_A$, and $ZS_B=F_B$.

Where $S_A>S_B$, and $tm$ is $\leq tc$, $V_B$ is calculated by Eq. 39.

$$V_B = \frac{Ca \cdot U \cdot tm}{2} - \frac{Z \cdot (S_A - S_B)}{\rho \cdot U} \cdot \ln(\frac{L_0 - U \cdot tm}{L_0}) \tag{39}$$

Where $S_A>S_B$, and $tm$ is $> tc$, $V_B$ is calculated by Eq. 40, noting that $tm$ cannot exceed $L_0/U$.

$$V_B = \frac{Ca \cdot U \cdot tc}{2} - \frac{Z \cdot (S_A - S_B)}{\rho \cdot U} \cdot \ln(\frac{L_0 - U \cdot tc}{L_0}) + Ca \cdot U \cdot (tm - tc) \tag{40}$$

Where $S_A<S_B$, and $tm$ is $\leq tc$, $V_B$ is calculated by Eq. 39.

Where $S_A<S_B$, and $tm$ is $> tc$, $V_B$ is calculated by Eq. 41.

$$V_B = \frac{Ca \cdot U \cdot tc}{2} - \frac{Z \cdot (S_A - S_B)}{\rho \cdot U} \cdot \ln(\frac{L_0 - U \cdot tc}{L_0}) \tag{41}$$

Where $S_A=S_B$, $V_B$ is calculated by Eq. 42.

$$V_B = \frac{Ca \cdot U \cdot tm}{2} \tag{42}$$

*Calculation of fluorescence acquired by simulated receptor cells*

As outlined above, cell fluorescence was expressed in 'Fluorescence Units' determined by summated fluorescence / cell surface profile area. There was wide variability in experimental fluorescence intensity of donor cells, so that fluorescence of individual simulated donor cells was taken into account when calculating fluorescence transfer to individual simulated receptor cells.

Imagine donor cell A with: cell surface profile area $Ar_A$; average cell height $Ht_A$; and fluorescence intensity $Fl_A$. Similarly imagine receptor cell B with: cell surface profile area $Ar_B$; average cell height $Ht_B$; and fluorescence intensity $Fl_B$ after acquisition of a volume $(V_B)$ of cytoplasm from donor cell A. The volume of donor cell A $(Vol_A)$ is given by $Ar_AHt_A$. Fluorescence in donor cell A is carried by a discrete number of fluorescent elements $(Fln)$, assumed to have uniform fluorescence and distribution throughout cell cytoplasm, such that the concentration of fluorescent elements per unit volume for donor cell A $(FlCon_A)$ is given by Eq 43. Note that $Fl_A = Fln/Ar_A$ which can be rearranged to $Fln = Fl_AAr_A$.

$$FlCon_A = \frac{Fln}{Vol_A} = \frac{Fln}{Ar_A \cdot Ht_A} \tag{43}$$

The number of fluorescent elements transferred from donor cell A to receptor cell B in $V_B$ $(Flnt)$ is given by multiplication of Eq. 43 by $V_B$ as in Eq. 44.

$$Flnt = \frac{V_B \cdot Fln}{Ar_A \cdot Ht_A} \tag{44}$$

Since $Fln = Fl_A Ar_A$, substitution into Eq. 44 give Eq. 45. Because $Fl_B = Flnt / Ar_B$, $Fl_B$ is given by multiplication and simplification of Eq. 45 as shown in Eq. 46, and this was used to calculate fluorescence acquired from individual simulated CPP events in computer simulations.

$$Flnt = \frac{V_B \cdot Fl_A}{Ht_A} \tag{45}$$

$$Fl_B = \frac{V_B \cdot Fl_A}{Ar_B \cdot Ht_A} \tag{46}$$

Total fluorescence acquired by individual simulated receptor cells from multiple separate simulated exchange events, was determined by summation of all values of $Fl_B$ for the individual simulated receptor cell.

#### *Calculation of pressures at time zero*

Maximum pressure drop at time zero for Cell A and Cell B, was calculated from Eq. 15, substituting $L_A$ and $L_B$ for $L(t)$ *and* $x$, to give $\Delta P_A$ and $\Delta P_B$ respectively.

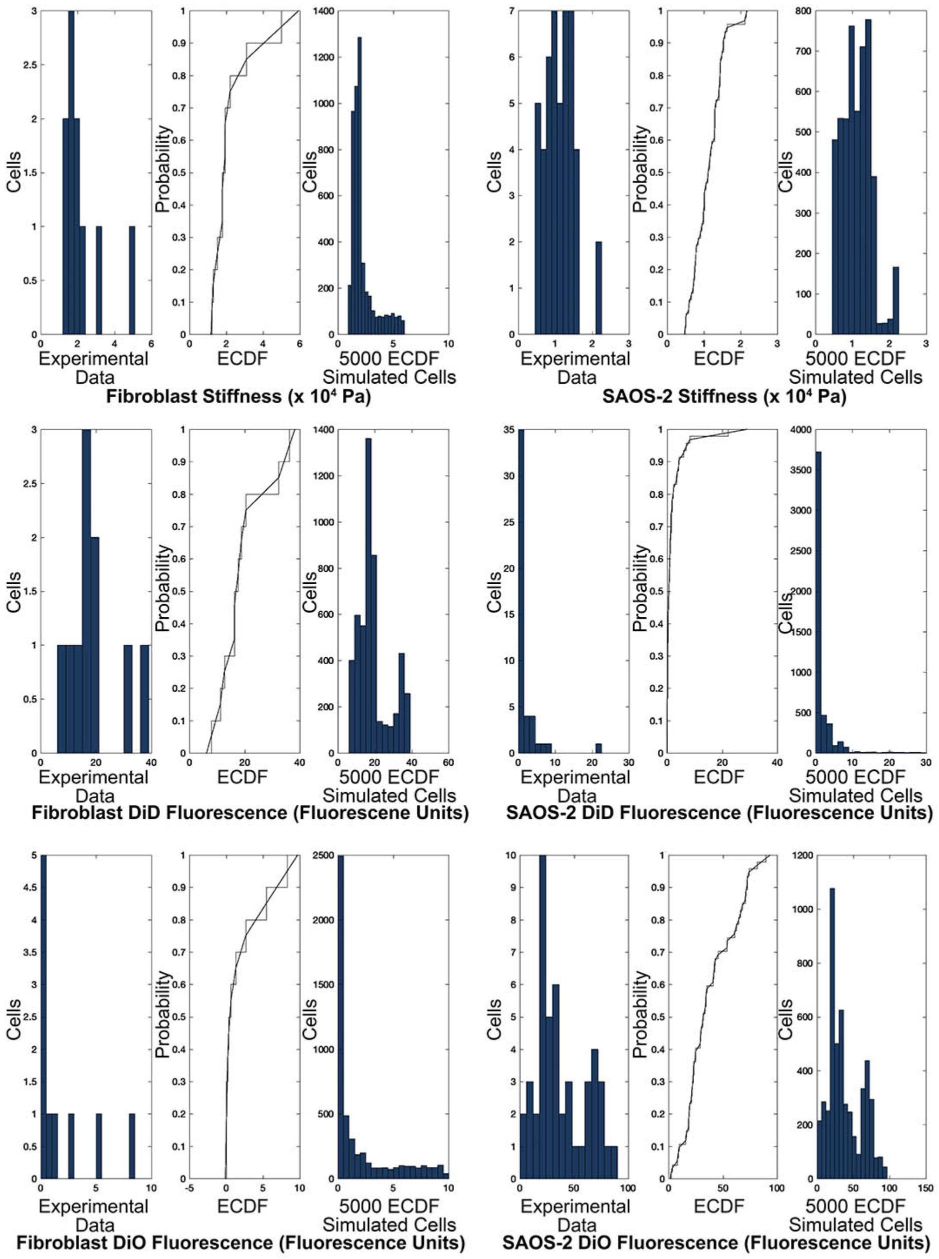


FIGURE S1 Histograms for experimental data for stiffness, DiD fluorescence and DiO fluorescence in co-cultured SAOS-2 and fibroblasts, as well as stair plots for the ECDF of these data with superimposed linear smoothing, and histograms for 5000 simulated cells generated by the ECDFs shown. Histograms of simulated cells had distribution profiles very similar to that of experimental data, despite the limited sampling available. Binning for histograms was: 3000 kPa for fibroblast stiffness; 1500 kPa for SAOS-2 stiffness; 3 DiD fluorescence units for fibroblast DiD; 1.5 DiD fluorescence units for SAOS-2 DiD; 0.5 DiO fluorescence units for fibroblast DiO; and 6 DiO fluorescence units for SAOS-2 DiO.

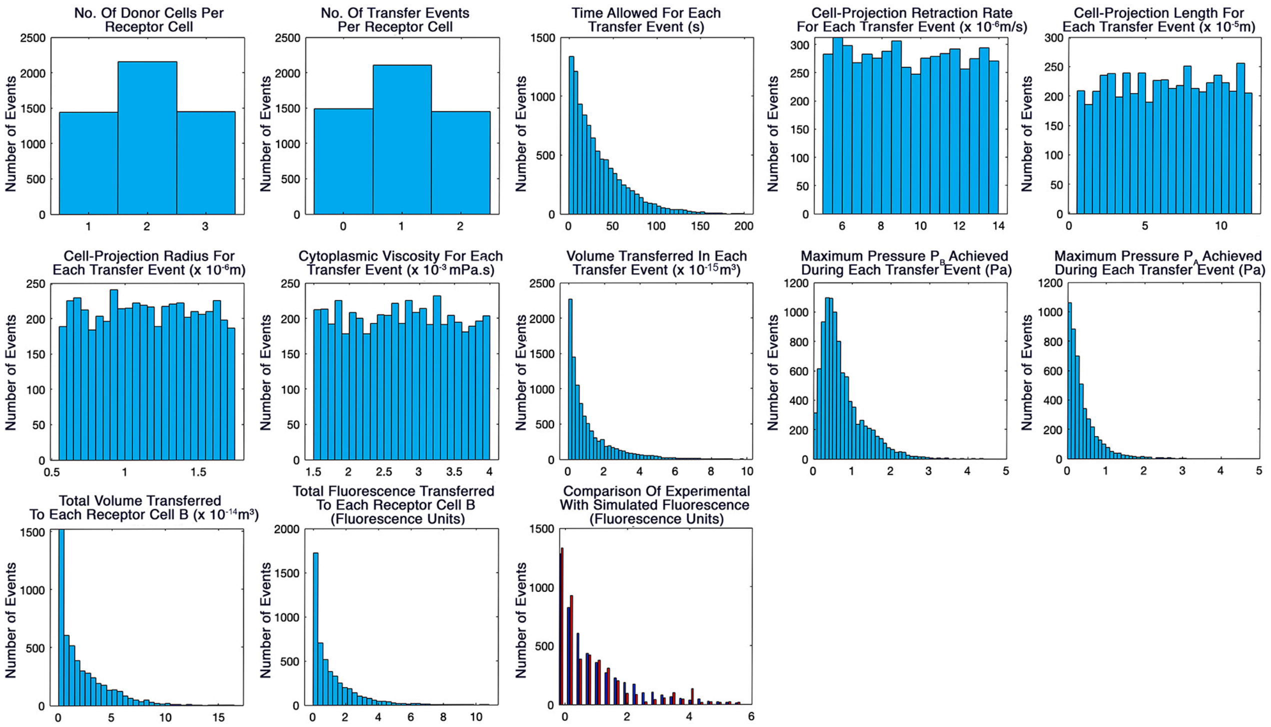


FIGURE S2 Histogram output of MATLAB script for simulation of CPP transfer from fibroblasts to SAOS-2 comparable to that seen by experiment (Fig. 7, Table 1 of the main manuscript; Supplemental Information Table S1). Modest central tendency was seen for the number of donor cells receptor cells interacted with, as well as for the number of transfer events per receptor cell. Input variables for cell-projection retraction rate, length, radius and viscosity had essentially uniform distributions, while time permitted for transfer events was within range of time-lapse observations. Pressures generated during transfer were modest, while volume and fluorescence transfer was appreciable, closely approximating experimental fluorescence data.

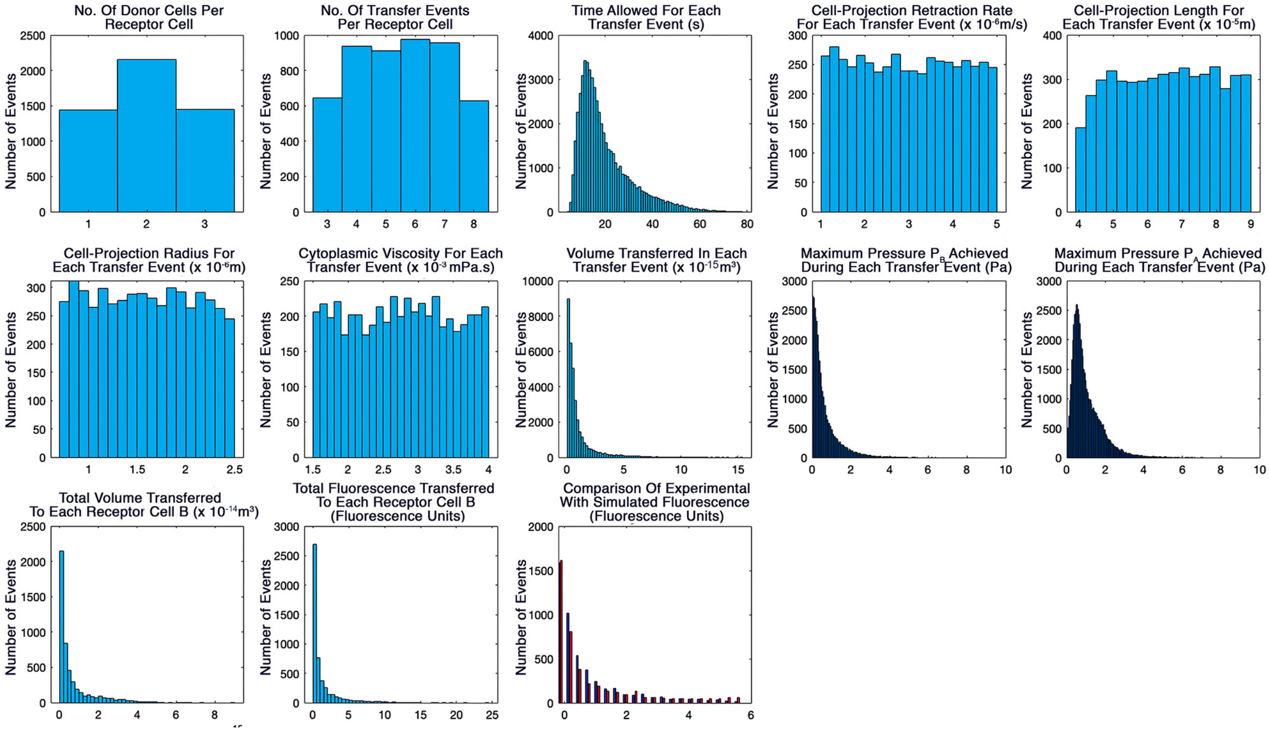


FIGURE S3 Histogram output of MATLAB script for simulation of CPP transfer from SAOS-2 to fibroblasts comparable to that seen by experiment (Fig. 7, Table 1 of the main manuscript; Supplemental Information Table S1). Modest central tendency was seen for the number of donor cells receptor cells interacted with, as well as for the number of transfer events per receptor cell. Input variables for cell-projection retraction rate, length, radius and viscosity had essentially uniform distributions, while time permitted for transfer events was within range of time-lapse observations. Pressures generated during transfer were modest while volume and fluorescence transfer was appreciable, closely approximating experimental fluorescence data.

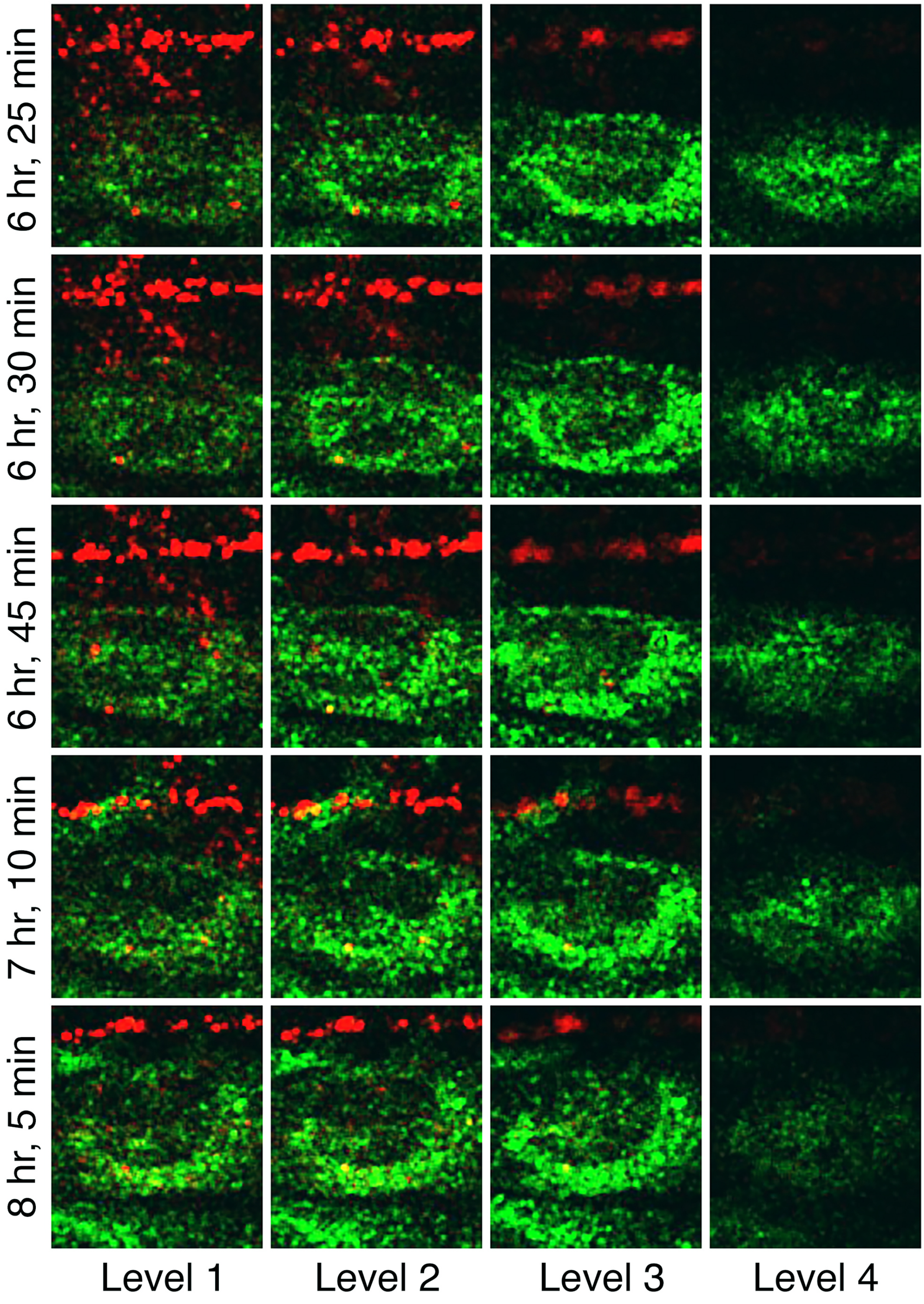


FIGURE S4 Consecutive confocal optical sections taken from Supplemental Movie S2 of a co-culture of MM200-B12 with fibroblasts at the time points shown in Fig. 2c,d of the main manuscript. MM200-B12 organelles were marked green with DiO. Fibroblast organelles marked red with DiD and transferred to MM200-B12, were clearly within the body of the malignant cell. The inferred paths taken by the fibroblast cell projections marked in Fig. 2d, were evident from examination of grooves in the malignant cell passing through and across the optical sections shown, and these sometimes contained fibroblast organellar cargo.

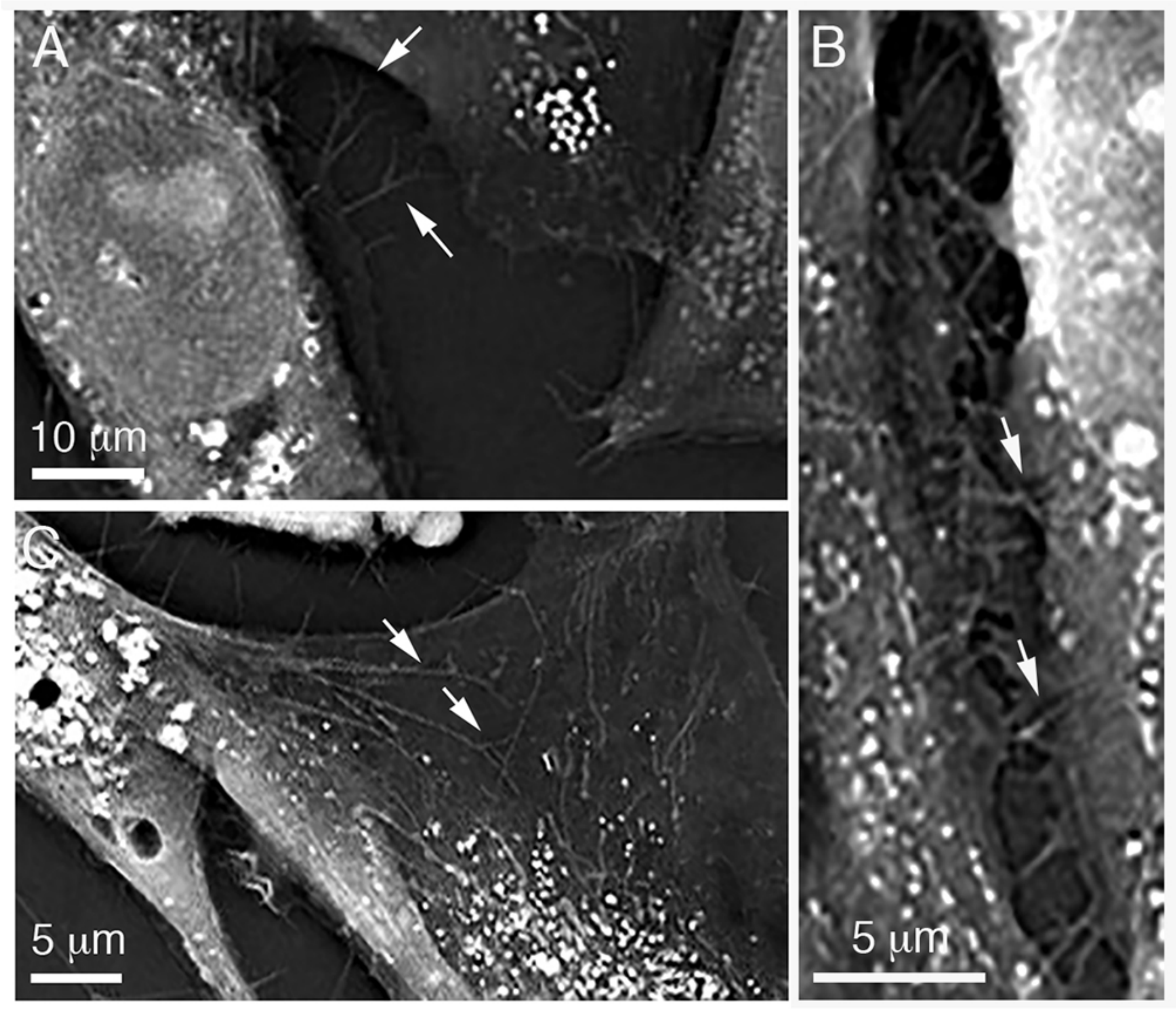


FIGURE S5 Holotomographic images at single optical levels of a co-culture of fibroblasts with SAOS-2 showing the presence of branching cell-projections consistent with those inferred from CLSM observations. A) Both branching (arrows) and non-branching cell-projections spread across the culture surfaces from cells. Cell-projections were often densely packed, while some grooved adjacent cells (B, arrows), and others spread across the surfaces of cells (C, arrows).

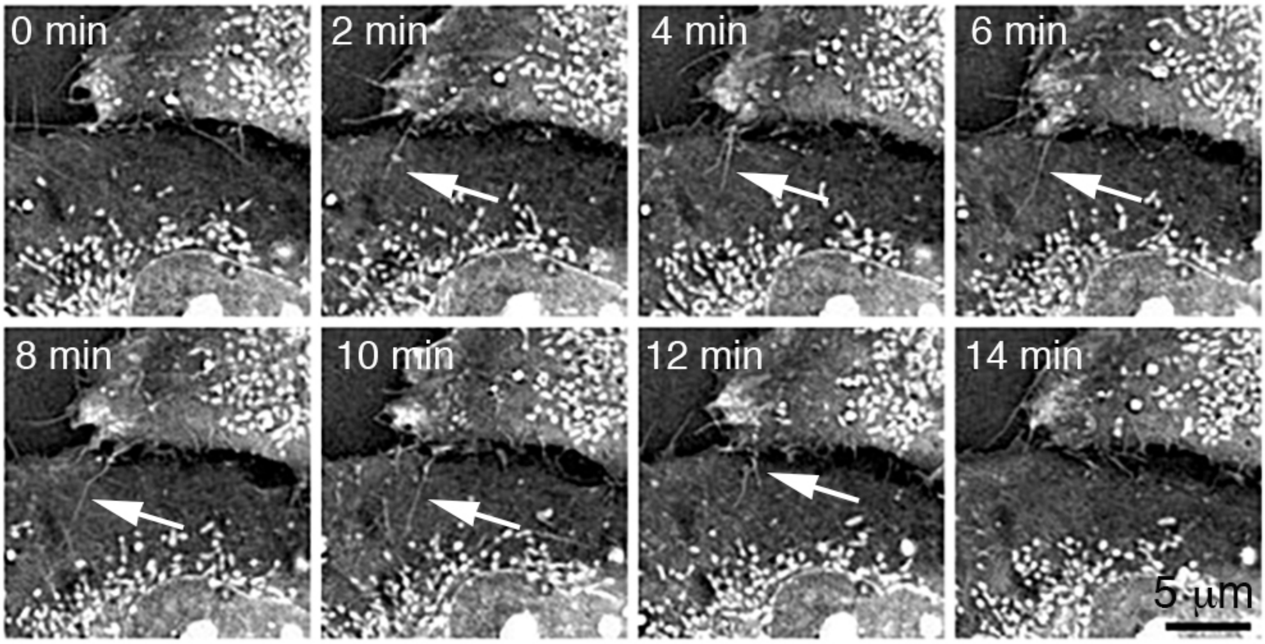


FIGURE S6 Sequential holotomographic time-lapse images at a single optical level of a co-culture of fibroblasts with SAOS-2, showing rapid movement of cell projections between time-points. Several separate cell projections were evident at the interface between two adjacent cells. These varied with regard to movement. One particularly active cell-projection (arrows) appeared at 2 min, and changed shape including formation of a branch at 12 min, before being extinguished by 14 min.

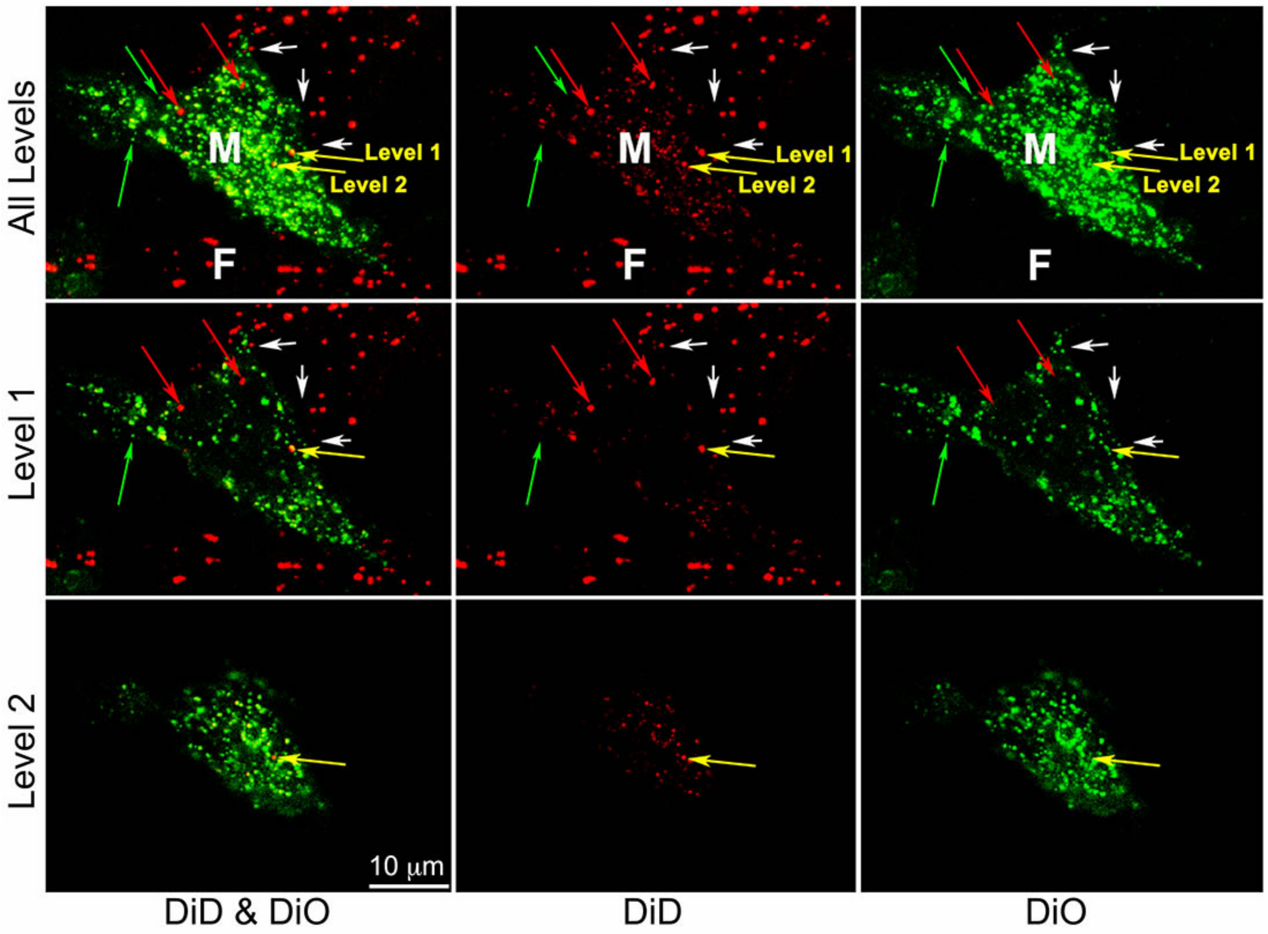


FIGURE S7 Flattened optical confocal microscopy projection and two optical levels of fibroblasts (pre-labelled with DiD-red) co-cultured for 24 Hrs with MM200-B12 (pre-labelled with DiO-green), showing channels for DiD and DiO alone or together. The malignant cell shown (M) had many organelles bearing both DiO and DiD. Two fibroblasts (F) were in close association with this cell, and one of these had broad cell-projections approaching the malignant cell (white arrows). Although many malignant cell organelles had both DiO and DiD labelling, some organelles appeared to have only DiO (green arrow), and there were occasional organelles where only DiD labelling was seen (red arrows), both suggestive of recent acquisition of organelles from neighboring cells. In addition, organelles were noted that were primarily marked with DiD, but which also had some DiO marker (yellow arrows) indicative of label mixing following organellar membrane recycling. Examination of separate optical levels (Levels 1 and 2) confirmed that organelles with mixed DiD and DiO labelling marked with yellow arrows, had actual dual labelling and were not an artifact of two coincidentally overlapping and oppositely labelled organelles. Observations are consistent with frequent transfer of organelles from adjacent fibroblasts and MM200-B12.

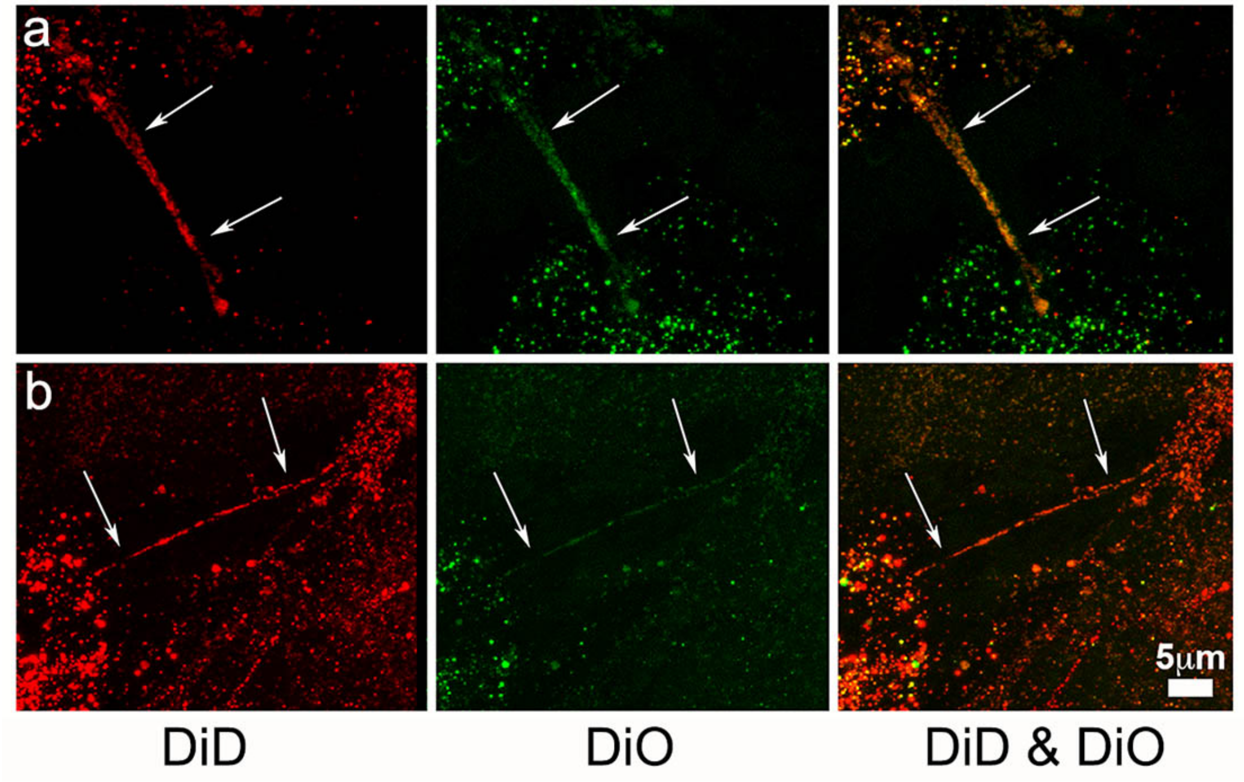


FIGURE S8 Flattened optical confocal microscopy projections of fibroblasts (pre-labelled with DiD-red) co-cultured for 24 Hr with MM200-B12 (pre-labelled with DiO-green), showing channels for DiD, DiO and both channels combined. Tunneling nanotubes (white arrows) were occasionally seen, often connecting malignant cells and bearing both DiD and DiO fluorescent markers. These did not, however, appear responsible for transfer of DiD labelled organelles from fibroblasts to malignant cells. Similar to data shown above in Fig. S7, there was significant diversity in the extent of DiD labelling amongst organelles within individual cells, suggestive of frequent uptake of fibroblast DiD. Occasional DiO organelles with little or no DiD in malignant cells otherwise heavily labelled with DiD, suggested recent uptake of DiO organelles from other malignant cells, consistent with our earlier report (1).

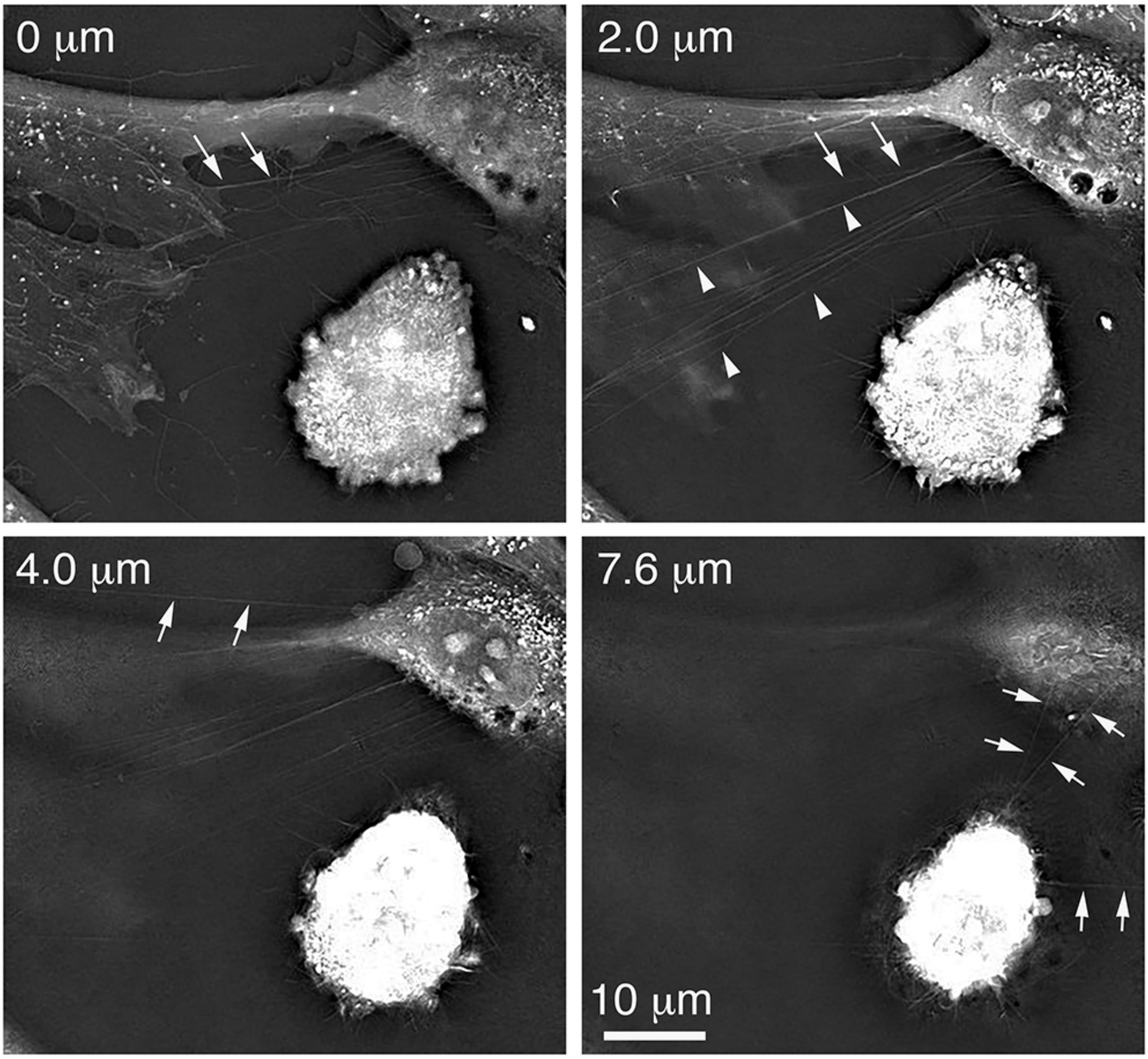


FIGURE S9 Holotomography demonstrating tunneling nanotubes in a co-culture of fibroblasts with SAOS-2 at a single time point and at four separate optical levels (0 µm, 2 µm, 4 µm and 7.6 µm). Tunneling nanotubes were recognized as straight linear structures suspended above the culture surface between cells (arrows and arrow heads). These sometimes spanned multiple optical levels (white arrows at optical levels 0 µm and 2 µm), while on other occasions they were more clearly apparent within a single optical level (arrow heads at 2 µm, arrows at 4 µm and 7.6 µm). Tunneling nanotubes were thus clearly different to the rapidly moving and occasionally branching cell-projections, that snaked across culture and cell surfaces and appeared responsible for the transfer events described in this report.

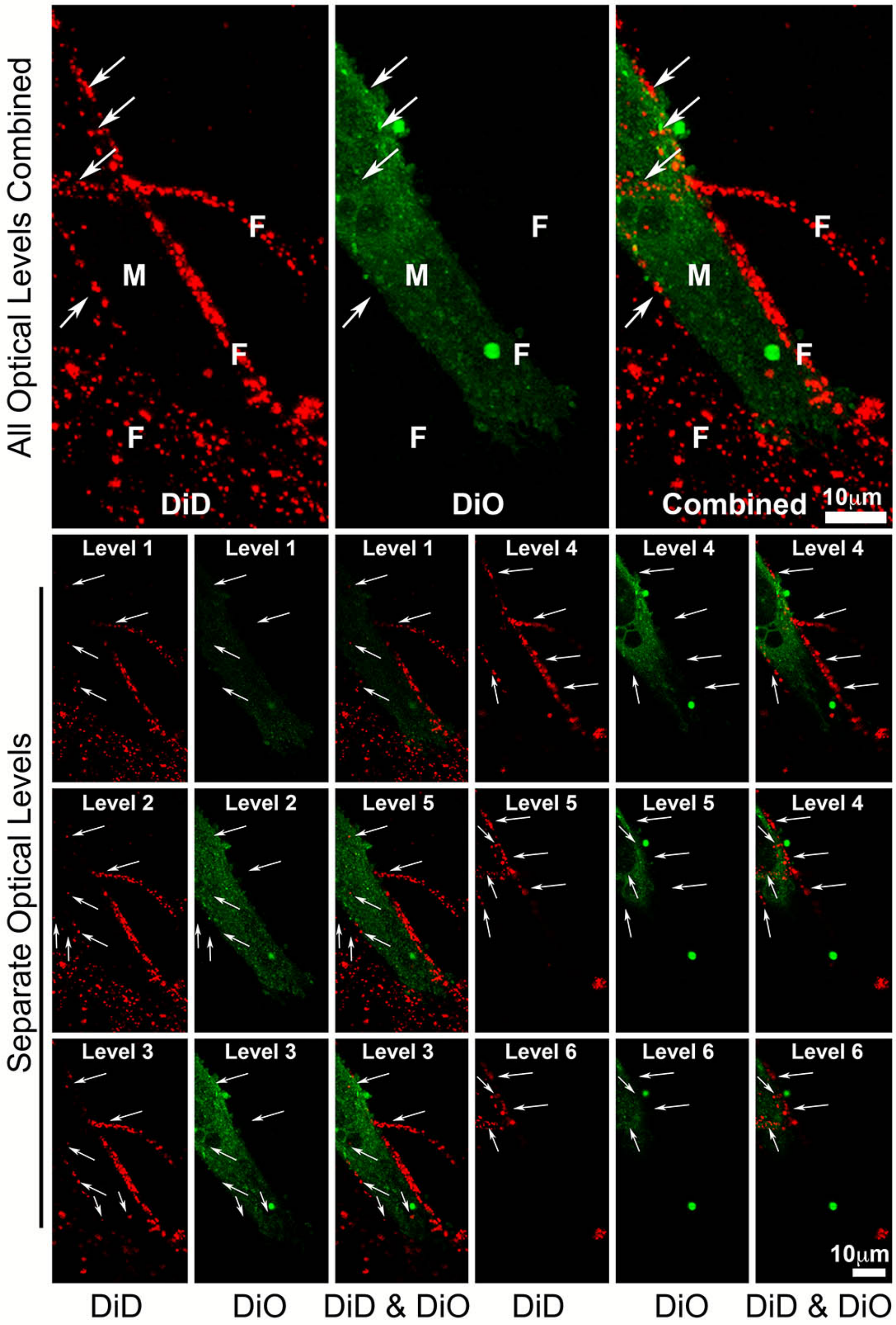


FIGURE S10 Flattened optical confocal microscopy projection and 6 optical levels of fibroblasts (pre-labelled with DiD-red) co-cultured with MM200-B12 (pre-labelled with DiO-green), showing channels for DiD and DiO alone and together. The malignant cell (M) shown had no clear DiD (red) labelling, despite complex entwining of the cell by fibroblast (F) processes indenting and grooving the malignant cell (white arrows), confirming that close physical association between malignant cells and fibroblasts was insufficient for transfer of fibroblast DiD label into malignant cells. This together with highly diverse fibroblast labelling of malignant cells across co-cultures, was inconsistent with either non-specific label exchange or an appreciable role for exosomes or other shed membrane vesicles in the label transfer studied.

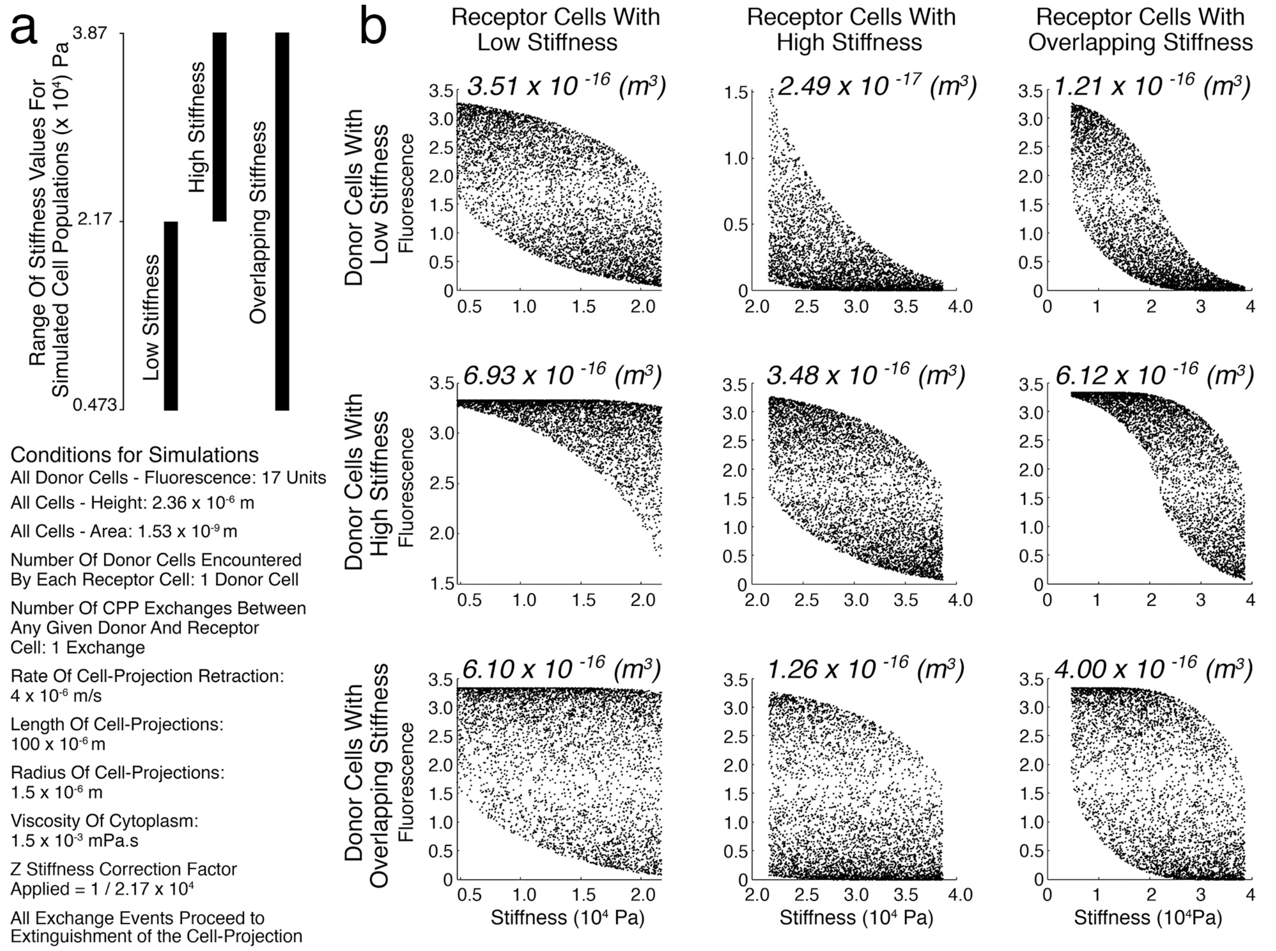


FIGURE S11 Scattergrams from computer simulations showing the relationship between stiffness of individual receptor cells and fluorescence acquired from donor cells, where stiffness of the two cell populations is varied, as well as the median CPP volume exchange for each simulation expressed in units of $m^3$. **(a)** Three cell populations were modelled: ‘Low Stiffness’ where stiffness had the same range as determined by experiment for SAOS-2; ‘High Stiffness’ with an equivalent range of stiffness, but where the lowest stiffness value was the same as the highest for the ‘Low Stiffness’ cell population; and ‘Overlapping Stiffness’ with stiffness covering the full range of both ‘High’ and ‘Low’ stiffness cell populations. 5100 donor and 5000 receptor cells were modelled in all simulations, with each cell assigned a unique stiffness value in equidistant steps from lowest to highest stiffness. The order of cells was randomized prior to simulations. For helpful simplicity, a single CPP event was modelled for each cell, and all variables were identical for all CPP exchanges as indicated. **(b)** As predicted from the mathematical model, considering pairings between each of the three cell populations modelled, there was preferential CPP transfer from more to less stiff cell populations, as reflected by higher median CPP volume exchanges. Also, although the shape of data clouds varied between donor-receptor pairings and identity of receptor cells, negative correlation between receptor cell stiffness and CPP fluorescence uptake was a consistent feature, as expected from the mathematical model. While more complex distributions for input variables will generate more complex data clouds, these simulations support generality for these two principal predictions from the mathematical model.

### LEGENDS FOR MOVIES

MOVIE S1 Confocal microscopy time-lapse movie showing the flattened optical projection of a field recorded for 3 h 36 min during a 25 h time-lapse experiment co-culturing MM200-B12 (pre-labelled with nuclear and cytoplasmic GFP, Green) with fibroblasts (pre-labelled with DiD, Red). Please note that Fig. 1a of the main manuscript, provides still images from this movie. A fibroblast cell-projection evident by multiple DiD (red) labeled organelles approached the lightly GFP labelled malignant cell towards the top left of the visual field. A single large organelle was deposited in the malignant cell during a retractive movement of the fibroblast cell-projection. Towards the end of the movie, labeling from the transferred organelle was distributed amongst other organelles in the recipient cell, interpreted as evidence for organellar turnover and membrane recycling.

MOVIE S2 Confocal microscopy time-lapse movie showing the flattened optical projection of a field recorded during an 8 h 15 min time-lapse experiment co-culturing MM200-B12 (pre-labelled with DiO, Green) with fibroblast (pre-labelled with DiD, Red), with channels for both DiD and DiO shown in the left image and for DiD only in the image on the right. Please note that Fig. 2 in the main manuscript, provides still images from this movie. Several malignant cells were seen, but attention is drawn to the MM200-B12 cell towards the screen outlined with a dashed line in Fig 2 of the main manuscript. Two partial images of fibroblasts were in the visual field, with one fibroblast cell-projection approaching the outlined malignant cell from the top left of the visual field, and sweeping towards the right, and another traversing the visual field in the approximate mid height of the images shown. Little DiD (red) labeling was seen in the malignant cell marked of interest in Fig. 2 of the main manuscript at the beginning of the time-lapse shown. However, numerous small and at least one large organelle were transferred into the malignant cell from the fibroblast cell-projection sweeping across the visual field by the end of the experiment, and these were seen as a cloud of small red particles surrounding the malignant cell nucleus, especially apparent in the image of the DiD channel only. Organelles appeared to be transferred across multiple branching filopodia-like fibroblast cell-projections, that were mostly transparent with exception of the grooves they formed in the surfaces of malignant cell and occasional DiD positive organelles (highlighted in Fig. 2 of the main manuscript). Towards the end of the movie, the donating fibroblast cell-projection withdrew from the malignant cell, and also drew with it the coincidentally horizontal and separate fibroblast cell-projection, across which it had passed. The withdrawal appeared associated with a particularly strong deposition event, while the energy inherent in retraction of the fibroblast cell-projection seemed underscored by the physical displacement of the geometrically near normal non-exchanging fibroblast process.

MOVIE S3  Confocal microscopy time-lapse movie showing the flattened optical projections of a field recorded for 3 h 35 min of MM200-B12 (pre-labelled with DiO) co-cultured with HDF (pre-labelled with DiD). A malignant cell entering the visual field from the top right displayed greater motility compared with most other malignant cells observed, and appeared to actively probe HDF with which it came in contact. Adherent lamelipodia from the malignant cell transferred DiO organelles to a HDF moving towards the left of the visual field.  Transfer of DiO organelles to the HDF was related to retraction of the malignant cell from the recipient HDF. This was the only occasion during which clear organellar transfer was observed by time-lapse from a malignant cell to a fibroblast.

MOVIE S4  Confocal microscopy time-lapse movie showing the flattened optical projection of a field recorded for 6 h during a 25 h time-lapse experiment co-culturing MM200-B12 (pre-labelled with nuclear and cytoplasmic GFP, Green) with fibroblast (pre-labelled with DiD, Red). Please note that Figs 1b&c of the main manuscript, provide still images from this movie. Two fibroblasts (red) interacted with a strongly labeled malignant cell (green), such that one fibroblast approached the cell from the top left of the visual field, and moved out of frame to the right, while another fibroblast extended a cell-projection to the malignant cell from the bottom of the visual field.  The fibroblast in the upper visual field interacted with the malignant cell at both its leading and trailing ends, while the fibroblast cell-projection approaching from the lower visual field repeatedly probed the malignant cell with branching cell-projections.  Both fibroblasts deeply indented the malignant cell with their cell-projections, providing a clear visual impression that the fibroblasts were much more stiff and mobile compared with malignant cell. Notably, the deep indentations demonstrated that the otherwise transparent fibroblast cell-projections comprised branching structures that terminated in long filopodia-like extensions that varied in precise shape from frame to frame, but nonetheless retained broadly similar form across several adjacent time points. While the movie shown makes the shape of fibroblast cell-projections particularly clear, exchange events for this cell were not as clear as that shown in ESM-2 and 3.

MOVIE S5 Movie of a rotating confocal microscopy image from a fixed monolayer of MM200-B12 (pre-labelled with DiO, Green) co-cultured with HDF (pre-labelled with DiD, red). The malignant cell was approached by a large fibroblast cell-process which deeply grooved the malignant cell, passing from below to above the malignant cell to create the appearance of the malignant cell having been 'speared' by the fibroblast. Grooving spaces in the malignant cell were apparent in the rotating image, that were inferred as having been formed by fibroblast cell-projections (separate grooves marked dark blue, light blue and white). A fibroblast cell-projection containing fibroblast organelles is marked with a red arrow.

MOVIE S6 Movie of holotomography at a single optical level of a co-culture of SAOS-2 with HDF illustrated in Fig. 3 of the main manuscript. Numerous cell-projections were both on the culture surface and spread across neighbouring cells. These varied greatly with regard to movement, such that some were comparatively static and others highly mobile. Two of the cell-projections extending from the cell in the upper field of view were particularly more mobile than others (white arrow, orange arrow), and both of these deeply indented and grooved the surface of the cell in the lower field of view. The cell projection marked with the arrow in Fig. 3 and with the orange arrow in this movie, appeared to transfer bright organellar cargo to the neighbouring cell.

### SUPPLEMENTAL TABLE

**Table S1.** The proportionate percentage distribution of simulated recipient SAOS-2 and Fib, according to the volume acquired from the opposing cell type in simulated co-culture generating fluorescence profiles approximating those seen by experiment, expressed as percentages relative to the average volume of the respective recipient cell type (Fig. 8 of the main manuscript).

| % Volume Relative to Average Recipient Cell | Simulated SAOS-2 Number | % Relative to All Simulated SAOS-2 | % of Simulated SAOS-2 with Transfer over Threshold | Simulated Fibroblast Number | % Relative to All Simulated Fibroblasts | % of Simulated Fibroblasts with Transfer over Threshold |
|---|---|---|---|---|---|---|
| **0 to < 1** | 1301 | 26.02 | 100 | 2062 | 41.26 | 100 |
| **1 to < 3** | 925 | 18.5 | 73.98 | 1274 | 25.46 | 58.74 |
| **3 to < 5** | 661 | 13.22 | 55.48 | 523 | 10.46 | 33.28 |
| **5 to < 7** | 455 | 9.1 | 42.26 | 263 | 5.26 | 22.82 |
| **7 to < 9** | 376 | 7.52 | 33.16 | 176 | 3.54 | 17.56 |
| **9 to < 11** | 312 | 6.24 | 25.64 | 135 | 2.58 | 14.02 |
| **11 to < 13** | 232 | 4.64 | 19.4 | 141 | 2.82 | 11.44 |
| **13 to < 15** | 205 | 4.1 | 14.76 | 110 | 2.26 | 8.62 |
| **15 to < 17** | 164 | 3.28 | 10.66 | 88 | 1.76 | 6.36 |
| **17 to < 19** | 110 | 2.2 | 7.38 | 80 | 1.6 | 4.6 |
| **19 to < 21** | 64 | 1.28 | 5.18 | 52 | 1.08 | 3 |
| **21 to < 23** | 64 | 1.28 | 3.9 | 26 | 0.52 | 1.92 |
| **23 to < 25** | 45 | 0.9 | 2.62 | 21 | 0.42 | 1.4 |
| **25 to < 27** | 23 | 0.46 | 1.72 | 19 | 0.38 | 0.98 |
| **27 to < 29** | 21 | 0.42 | 1.26 | 16 | 0.32 | 0.6 |
| **29 to < 31** | 15 | 0.3 | 0.84 | 4 | 0.08 | 0.28 |
| **31 to < 33** | 6 | 0.12 | 0.54 | 0 | 0 | 0.2 |
| **33 to < 35** | 11 | 0.22 | 0.42 | 3 | 0.06 | 0.2 |
| **35 to <37** | 3 | 0.06 | 0.2 | 4 | 0.08 | 0.14 |
| **37 to <39** | 1 | 0.02 | 0.14 | 0 | 0 | 0.06 |
| **39 to <41** | 3 | 0.06 | 0.12 | 2 | 0.04 | 0.06 |
| **41 to <43** | 0 | 0 | 0.06 | 0 | 0 | 0.02 |
| **43 to <45** | 2 | 0.04 | 0.06 | 0 | 0 | 0.02 |
| **45 to <47** | 1 | 0.02 | 0.02 | 0 | 0 | 0.02 |
| **47 to <49** | 0 | 0 | 0 | 1 | 0.02 | 0.02 |
| **Total** | 5000 | 100 | | 5000 | 100 | |

Simulations predicting fluorescence profiles for recipient cells similar to experimental results (Fig. 7 of the Main Manuscript), also predicted transfer of appreciable volumes of cytoplasm to recipient cells. 55.5% of simulated SAOS-2 had over 3%, and 5.2% of SAOS-2 had over 19% volume acquired from simulated Fib. Consistent with occasional experimentally observed SAOS-2 with very high Fib DiD labelling, 10 simulated SAOS-2 cell acquired between 35% and 47% of their volume from simulated Fib. Simulated Fib also acquired appreciable cytoplasm from simulated SAOS-2, but this was generally less compared with exchange in the reverse direction.

### MATLAB SCRIPT

#### MATLAB script for generating simulated cell values from input data when the lowest value in input data is significantly greater than zero.

```
% ECDF SCRIPT FOR DATA NOT APPROACHING ZERO
%THIS IS A SCRIPT TO: Generate a determined number of values
%of a variable extrapolated from raw input data first establishing
%an Estimated Cumulative Distribution Function (ECDF), and then using
%the inverse of this to generate a set of random values with
%the same distribution as that of the raw data
%........................................................

%VARIABLES THAT MUST BE CHANGED WITH EACH NEW INPUT DATA
Input=RawDataList;

%Target value number can be changed as required.
Target=5000;

%The bin size of the input data for display in a histogram = bs as desired.
bs=1500;


%--------------------------------------------------------------

%DISPLAYING INPUT DATA
%Determinint the maximum value in Histdat
mx=max(Input);
%Determining the bounds of the binned range, upper =top
divis=(mx/bs);
divnum=round(divis);
top=(divnum*bs)+bs;
%Plotting a histogram
edges = [0 bs:bs:top-bs top];
subplot(1,3,1);
h = histogram(Input,edges);
%Clearing values no longer needed
clear divis h mx top

%--------------------------------------------------------------


%GENERATING AN EMPIRICAL CUMULATIVE DISTRIBUTION FUNCTION

% Where F = the Probability
[F,xValues]=ecdf(Input);

%Smoothing steps via a piecewise linear estimate
xSmooth=xValues(2:end);
FSmooth=(F(1:end-1)+F(2:end))/2;

%--------------------------------------------------------------------


%CALCULATING THE CONSTANTS FOR LINEAR EQUATIONS FOR THE SMOOTHING LINES
%OF THE ECDF

%Creating and Loading a matrix (ConstTab) to receive results where:
%Column 1= Minimum variable in range from xSmooth - (x1 Value1);
%Column 2= Maximum variable in range from xSmooth - (x2 Value2);
%Column 3= Minimum variable in range from FSmooth - (y1 Prob1);
%Column 4= Maximum variable in range from FSmooth - (y2 Prob2);
%Column 5=y2-y1 (Prob2 - Prob1); Column 6=x2-x1 (Value2 - Value 1)
%Column 7 = Gradient (Column 5/Column 6);
%Column 8 = y intercept

%Creating a matrix and loading values into columns 1 to 4
```

```
l=length(FSmooth);
ConstTab=zeros(l+1,8);
%Loading minimum and maximum values for x and y ranges
for a=1:l
    ConstTab(a+1,1)=xSmooth(a,1);
    ConstTab(a,2)=xSmooth(a,1);
    ConstTab(a+1,3)=FSmooth(a,1);
    ConstTab(a,4)=FSmooth(a,1);
end

%Calculating y2-y1 and loading into Column 5
for a=1:l
    ConstTab(a+1,5)=...
        ConstTab(a+1,4)-ConstTab(a+1,3);
end

%Calculating x2-x1 and loading into Column 6
for a=1:l
    ConstTab(a+1,6)=...
        ConstTab(a+1,2)-ConstTab(a+1,1);
end

%Calculating gradient by (Column 5/Column 6)
for a=1:l
    ConstTab(a+1,7)=...
        ConstTab(a+1,5)/ConstTab(a+1,6);
end

%Calculating Y Intercept by b=y-mx
for a=1:l
    ConstTab(a+1,8)=...
        ConstTab(a+1,3)-(ConstTab(a+1,7).*ConstTab(a+1,1));
end

% Transferring results from Row 1 to Row 2, and Row l-1 to Row l
ConstTab(1,:) = ConstTab(2,:);
ConstTab(l+1,:) = ConstTab(l,:);

%Extending y values to 0 and 1 respectively
ConstTab(1,3)=0;
ConstTab(l+1,4)=1;

%Creating a SmoothGraph matrix from ConstTab that includes probability
%values of 0 and 1, together with relevant gradient and Y intercept values
%to allow plotting and overlay of the smoothed function from which
%values are ultimately calculated, over an ECDF stairs plot
%Column 2 =ConstTab Column 3 (Probability);
%Column 3= ConstTab Column 7 (Gradient);
%Column 4= ConsTab Column 8 (Y Intercept);
SmoothGraph=zeros(l+1,4);
SmoothGraph(1:l+1,2)=ConstTab(:,3);
SmoothGraph(1:l+1,3)=ConstTab(:,7);
SmoothGraph(1:l+1,4)=ConstTab(:,8);

%Loading 0 and 1 probability values into SmoothGraph
SmoothGraph(1:2)=0;
SmoothGraph(l+1,2)=1;

%Calculating Values for SmoothGraph from the formula y=ax+b
%=rearranged to x = (y-b)/a
%= (Column 2 - Column 4)/Column 3
for i=1:l+1
    SmoothGraph(i,1)=(SmoothGraph(i,2)-SmoothGraph(i,4))/SmoothGraph(i,3);
end

%Displaying stepwise ECDF with overlayed smoothed linear estimate curve
subplot(1,3,2)
stairs (xValues,F)
hold on
plot (SmoothGraph(:,1),SmoothGraph(:,2))
```

```
hold off
title ('ECDF Stairs and Smoothed')
xlabel ('Values')
ylabel ('Probability')

 %-----------------------------------------------------------------


%GENERATING A TARGET NUMBER OF RANDOMLY SELECTED VALUES FROM THE
%ECDF CURVE DEFINED BY THE LINER EQUATIONS SPECIFIED FROM ConstTab

%Creating a matrix ProbList for calculation of the
%Target number of values (where: y=ax+b)
%Column 1= Variable (=x); Column 2= Probability Randomly Selected(=y)
%Column 3 = Gradient (=a From ConstTab Column 7);
%Column 4 = Y Intercept (=b From ConstTab Column 8)
ProbList=zeros(Target,4);

%Creating a list of random numbers and loading into ProbList
Rand=rand(Target,1);
ProbList(:,2)=Rand(:,1);

%Selecting gradient and Y Intercept values for loading into
%ProbList
i=[1:Target];
j=[1:l+1];
for i=1:Target
    for j=1:l+1
        if ProbList(i,2) >= ConstTab(j,3) &&...
                ProbList(i,2) < ConstTab(j,4)
            ProbList(i,3) = ConstTab(j,7);
            ProbList(i,4) = ConstTab(j,8);
        end
    end
end

%Calculating random values from the formula y=ax+b
%=rearranged to x = (y-b)/a
%= (Column 2 - Column 4)/Column 3
for i=1:Target
    ProbList(i,1)=(ProbList(i,2)-ProbList(i,4))/ProbList(i,3);
end

ResultList=ProbList(:,1);

%-----------------------------------------------------------------


%DISPLAYING OUTPUT DATA = ResultList

%Determining the maximum value in ResultList
mx=max(ResultList);
%Determining the bounds of the binned range, upper =top
divis=(mx/bs);
divnum=round(divis);
top=(divnum*bs)+bs;
%Plotting the histogram
edges = [0 bs:bs:top-bs top];
subplot(1,3,3);
h = histogram(ResultList,edges);

%Clearing unwanted values from the workspace
clear bs divis h mx top a ConstTab divnum edges F FSmooth i Input j l
clear ProbList Rand SmoothGraph Target xSmooth xValues

 %END SCRIPT
```

**MATLAB script for generating simulated cell values from input data when the lowest value in input data does not approach zero.**

```
%ECDF SCRIPT FOR DATA APPROACHING ZERO
% THIS IS A SCRIPT TO: Generate a determined number of values
%of a variable extrapolated from raw input data first establishing
%an Estimated Cumulative Distribution Function (ECDF), and then using
%the inverse of this to generate a set of random values with
%the same distribution as that of the raw data
%NOTE: This Script replaces negative values, and values < lowest
%value in input data, with ?0?. If negative values are expected
%in the output, relevant part of script must be removed by ?%?.


%---------------------------------------------------------------

%VARIABLES THAT MUST BE CHANGED WITH EACH NEW INPUT DATA

Input=RawDataList;

%Target value number can be changed as required.
.Target=5000;

%The bin size of the input data for display in histogram = bs as desired.
bs=0.5;


%---------------------------------------------------------------

%DISPLAYING INPUT DATA

%Determining the maximum value in Histdat
mx=max(Input);
%Determining the bounds of the binned range, upper =top
divis=(mx/bs);
divnum=round(divis);
top=(divnum*bs)+bs;
%Plotting a histogram
edges = [0 bs:bs:top-bs top];
subplot(1,3,1);
h = histogram(Input,edges);
%Clearing unwanted values from the workspace
%clear bs
clear divis h mx top


%---------------------------------------------------------------

%GENERATING AN EMPIRICAL CUMULATIVE DISTRIBUTION FUNCTION

% Where F = the Probability
[F,xValues]=ecdf(Input);

%Smoothing steps via a piecewise linear estimate
xSmooth=xValues(2:end);
FSmooth=(F(1:end-1)+F(2:end))/2;

%-----------------------------------------------------------------

%CALCULATING CONSTANTS FOR LINEAR EQUATIONS FOR THE SMOOTHING LINES OF THE ECDF

%Creating and loading a matrix (ConstTab) to receive results where:
%Column 1= Minimum variable in range from xSmooth - (x1 Value1);
%Column 2= Maximum variable in range from xSmooth - (x2 Value2);
%Column 3= Minimum variable in range from FSmooth - (y1 Prob1);
```

```
%Column 4= Maximum variable in range from FSmooth - (y2 Prob2);
%Column 5=y2-y1 (Prob2 - Prob1); Column 6=x2-x1 (Value2 - Value 1)
%Column 7 = Gradient (Column 5/Column 6);
%Column 8 = y intercept

%Creating the matrix and loading values into Columns 1 to 4
l=length(FSmooth);
ConstTab=zeros(l+1,8);
%Loading minimum and maximum values for x and y ranges
for a=1:l
    ConstTab(a+1,1)=xSmooth(a,1);
    ConstTab(a,2)=xSmooth(a,1);
    ConstTab(a+1,3)=FSmooth(a,1);
    ConstTab(a,4)=FSmooth(a,1);
end

%Calculating y2-y1 and loading into Column 5
for a=1:l
    ConstTab(a+1,5)=...
        ConstTab(a+1,4)-ConstTab(a+1,3);
end

%Calculating x2-x1 and loading into Column 6
for a=1:l
    ConstTab(a+1,6)=...
        ConstTab(a+1,2)-ConstTab(a+1,1);
end

%Calculating Gradient by (Column 5/Column 6)
for a=1:l
    ConstTab(a+1,7)=...
        ConstTab(a+1,5)/ConstTab(a+1,6);
end

%Calculating Y Intercept by b=y-mx
for a=1:l
    ConstTab(a+1,8)=...
        ConstTab(a+1,3)-(ConstTab(a+1,7).*ConstTab(a+1,1));
end

% Transferring results from Row 1 to Row 2, and Row l-1 to Row l
ConstTab(1,:) = ConstTab(2,:);
ConstTab(l+1,:) = ConstTab(l,:);

%Extending y values to 0 and 1 respectively
ConstTab(1,3)=0;
ConstTab(l+1,4)=1;

%Creating a SmoothGraph matrix from ConstTab that includes probability
%values of 0 and 1, together with relevant gradient and Y intercept values
%to allow plotting and overlay of the smoothed function from which
%values are ultimately calculated, over an ECDF stairs plot
%Column 2 =ConstTab Column 3 (Probability);
%Column 3= ConstTab Column 7 (Gradient);
%Column 4= ConsTab Column 8 (Y Intercept);
SmoothGraph=zeros(l+1,4);
SmoothGraph(1:l+1,2)=ConstTab(:,3);
SmoothGraph(1:l+1,3)=ConstTab(:,7);
SmoothGraph(1:l+1,4)=ConstTab(:,8);

%Loading 0 and 1 probability values into SmoothGraph
SmoothGraph(1:2)=0;
SmoothGraph(l+1,2)=1;

%Calculating values for SmoothGraph from the formula y=ax+b
%=rearranged to x = (y-b)/a
%= (Column 2 - Column 4)/Column 3
for i=1:l+1
    SmoothGraph(i,1)=(SmoothGraph(i,2)-SmoothGraph(i,4))/SmoothGraph(i,3);
end
```

```
%Displaying stepwise ECDF with overlaid smoothed linear estimate curve
subplot(1,3,2)
stairs (xValues,F)
hold on
plot (SmoothGraph(:,1),SmoothGraph(:,2))
hold off
title ('ECDF Stairs and Smoothed')
xlabel ('Values')
ylabel ('Probability')

%------------------------------------------------------------------

%GENERATING A TARGET NUMBER OF RANDOMLY SELECTED VALUES FROM THE
%ECDF CURVE DEFINED BY THE LINER EQUATIONS SPECIFIED FROM ConstTab

%Creating a matrix ProbList for calculation of the
%Target number of values (where: y=ax+b)
%Column 1= Variable (=x); Column 2= Probability Randomly Selected(=y)
%Column 3 = Gradient (=a From ConstTab Column 7);
%Column 4 = Y Intercept (=b From ConstTab Column 8)
ProbList=zeros(Target,4);

%Creating a list of random numbers and loading into ProbList
Rand=rand(Target,1);
ProbList(:,2)=Rand(:,1);

%Selecting Gradient and Y Intercept Values for Loading into
%ProbList
i=[1:Target];
j=[1:l+1];
for i=1:Target
    for j=1:l+1
        if ProbList(i,2) >= ConstTab(j,3) &&...
                ProbList(i,2) < ConstTab(j,4)
            ProbList(i,3) = ConstTab(j,7);
            ProbList(i,4) = ConstTab(j,8);
        end
    end
end

%Calculating Random Values from the formula y=ax+b
%=rearranged to x = (y-b)/a
%= (Column 2 - Column 4)/Column 3
for i=1:Target
    ProbList(i,1)=(ProbList(i,2)-ProbList(i,4))/ProbList(i,3);
end

ResultList=ProbList(:,1);

%------------------------------------------------------------------

%REMOVING NEGATIVE VALUES AND VALUES LESS THAN THE LOWEST POSITIVE
%VALUE OF INPUT

MinIn=min(Input);
for i=1:Target
    if ResultList(i,1) < MinIn;
        ResultList(i,1) = 0;
    end
end

%------------------------------------------------------------------

%DISPLAYING OUTPUT DATA = ResultList

%Determine the maximum value in ResultList
mx=max(ResultList);
%Determining the bounds of the binned range, upper =top
divis=(mx/bs);
```

```
divnum=round(divis);
top=(divnum*bs)+bs;
%Plotting a histogram
edges = [0 bs:bs:top-bs top];
subplot(1,3,3);
h = histogram(ResultList,edges);

%Clearing unwanted values from the workspace
clear bs divis h mx top a ConstTab divnum edges F FSmooth i input
clear j l MinIn ProbList Rand SmoothGraph Target xSmooth xValues Input

%END SCRIPT
```

### MATLAB script for simulating cytoplasmic exchange between populations of cells via cell-projection pumping.

```
%CPP SIMULATION SCRIPT
%-------------------------------------------------------------------
%-------------------------------------------------------------------
%THIS IS A SCRIPT FOR:
%Modeling transfer of cytoplasm and fluorescent label via Cell-Projection
%Pumping (CPP)between a determined number of cells A to Cells B.

%The cells modeled are SAOS-2 osteosarcoma cells, and HDF fibroblasts,
%while HDF and SAOS-2 can both be assigned to be cell A.

%Nonetheless, this script can be readily modified to model exchange between
%any population of cells, given relevant distributions for†
%stiffness and fluorescence, as well as average values for cell height
%and cell surface profile area.

%Cells are connected via cell-projections modelled as cylindrical†
%structures with variable radius 'r' and length 'l'.

%Pressure arises by shortening of cell-projections, and drives flow
%of cytoplasm outwards from cell-projections towards Cells A and B.

%The maximum possible flow out of any single cell-projection is the volume
%of the cell-projection.†

%It is assumed that the contents of cell-projections are always from
%Cell A.

%Modeling is for a defined number of Cells B, receiving
%cytoplasm from a defined number of Cells A.

%Stiffness values for both cells A and B are drawn at
%random from stiffness value lists determined separately via
%Estimated Cumulative Distribution Functions (ECDF), which have
%in turn be determined from experimental data.

%Each Cell A also has a single defined fluorescence level, according to
%the observed fluorescence labelling in co-cultures of SAOS-2 with HDF,
%and also established by ECDF.

%Experimental data is also used to determine by ECDF the fluoresence
%acquired by Cells B from Cells A, for comparison with simulation†
%results.

%The cumulative simulated volume of cytoplasm and fluorescent label uptake
%by Cells B is determined, dependent on the variables shown below,
%all determined from† distributions between defined target minima and†
%maxima. Maximum pressure for each exchange event is also calculated.

%Key input variables are:
%Rate of cell-projection retraction (Rate) (Units of m/s)
%Number of Cells A with which a given Cell B interacts (ANum)
%Number of Exchanges in any given cell pairing (ExNum)
%Length of cell-projections (Lgh) (Units of m)
%Radius of cell-projections (Radi) (Units of m)
%Viscosity (Visc) (Units of mPa.s)
```

```
%Time for each Exchange to occur (Tim) (Units of s), is selected
%at random between defined minimum and maximum proportionate values
%calculated from the maximum possible time of retraction for any
%given cell-projection

%----------------------------------------------------------------------
%----------------------------------------------------------------------
%VARIABLES TO INPUT FOR EACH SIMULATION

%NOTE: The number of input values for AStiff and AFluor must
%be greater than for BStiff, by at least as much as the
%maximum number of Cells A any given single cell B can interact with.
%It has been convenient for the number of Cells A to exceed that of†
%Cells B by 100.†

%The terms 'FibStiff', 'SAOSStiff', 'FibRed', 'SAOSRed', FibGreen,
%'SAOSGreen', are shorthand for data files used†in this study, but
%are not required names.

%The '%' symbol is used to activate and inactivte the below lines
%as needed.

%Stiffness units are in Pa.

AStiff=FibStiff;
% AStiff=SAOSStiff;

BStiff=SAOSStiff;
% BStiff=FibStiff;

AFluor=FibRed;
% AFluor=SAOSGreen;

ExpFluor=SAOSRed;
% ExpFluor=FibGreen;

...............................
%HEIGHT AND AREA OF CELLS A AND B
%Use '%' to activate or inactivate measures below, dependent on which
%cell type is Cell A and B respectivelly. These values must be changed†
%if other cell types are being measured. Units are in m, and values†
%used in this simulation are from atomic force microscopy measures

%If Cells A are HDF:
AHght=2.36*10^-6;
BAr=1.53*10^-9;

%If Cells A are SAOS-2:
% AHght=3.89*10^-6;
% BAr=5.34*10^-9;

%---------------------------------------------------------------------
%---------------------------------------------------------------------
%DEFINING STIFFNESS CORRECTION CONSTANT Z
%The experimental method for determining stiffness used in the current†
%script is atomic force microsocpy (AFM) of fixed cells. This necessitates
%introduction of a Stiffness Correction Constant (Z).
%Letting Z=1/(absolute value of the median stiffness of all cells in the
%system), generates values for tc (defined below) that permit fair
%testing of the effect of difference in cell stiffness on fluorescence
%transfer between cells by CPP.

MedS=median([AStiff;BStiff]);
Z=(1/abs(MedS));

%---------------------------------------------------------------------
%---------------------------------------------------------------------
%DEFINING VARIABLES BELOW FOR EACH SIMULATION
%Number of Cells B
BNo=5000;

%.............................
```

```
%DISTRIBUTIONS FOR INPUTS
%A 'Normal'distribution (SD) is given by the formula:†
%SD=((Max-Min/2)*0.3).
%For purposes of trialing differing distributions for variables
%it is helpful to distort the distribution away from 'Normal'
%by changing the SD value. Increasing SD above 0.3 flattens and†
%spreads the distribution, while reducing SD to be less than 0.3
%narrows and makes more acute the distribution. A 'wide-flat'
%distribution can thus be achieved with a high 'SD' value, while†
%a 'narrow-sharp' distribution is created by a low 'SD value.†

%Defining the extent to which SD varies from 'Normal' (SD = 0.3)
SD=2;

%Number of Cells A with which any given single Cell B may exchange
MinANum= 1;
MaxANum= 3;
MidANum=MinANum+((MaxANum-MinANum)/2);
SDANum=((MaxANum-MinANum)/2)*SD;

%Number of Exchanges between any single Cell A and Cell B
MinExNum= 0;
MaxExNum= 2;
MidExNum=MinExNum+((MaxExNum-MinExNum)/2);
SDExNum=((MaxExNum-MinExNum)/2)*SD;

%CPP Retraction Rate for any given cell-projection retraction event
MinRate= 0.5*10^-6;
MaxRate= 1.4*10^-6;
MidRate=MinRate+((MaxRate-MinRate)/2);
SDRate=((MaxRate-MinRate)/2)*SD;

%Length of cell-projection in any given exchange between Cell A and Cell B
MinLgh= 5*10^-6;
MaxLgh= 120*10^-6;
MidLgh=MinLgh+((MaxLgh-MinLgh)/2);
SDLgh=((MaxLgh-MinLgh)/2)*SD;

%Radius of cell-projection in any given exchange between Cell A and Cell B
MinRadi= 0.55*10^-6;
MaxRadi= 1.75*10^-6;
MidRadi=MinRadi+((MaxRadi-MinRadi)/2);
SDRadi=((MaxRadi-MinRadi)/2)*SD;

%Viscosity of Cytoplasm exchanged from Cell A to Cell B
MinVisc= 1.5*10^-3;
MaxVisc= 4*10^-3;
MidVisc=MinVisc+((MaxVisc-MinVisc)/2);
SDVisc=((MaxVisc-MinVisc)/2)*SD;


%Time for any given exchange between Cell A and Cell B
%An upper bound for the maximum possible time (PosTim) for CPP is
%established by l/U. MinTim = PropMin*PosTim; MaxTim=PropMax*PosTim

PropMin= 0;
PropMax= 0.9;

%-------------------------------------------------------------------
%-------------------------------------------------------------------
%CREATING MATRIXES WITH VARIABLES HAVING the PROBABILITY DISTRIBUTIONS
%DEFINED ABOVE

% ANum = Number of Cells A with which any given Cell B may exchange
% ExNum = Number of Exchanges any given Cell B may have with a given Cell A
% Lgh = Length of the cell-projection in any given exchange event
% Rate = Rate for any given exchange event
% Radi = Radius of cell-projection in any given exchange event
% Visc = Viscosity in any given exchange event

%USE '%' TO SELECT OR REMOVE 'TWISTER'
%Use of 'Twister' to make random values reproducible -
rng(0,'twister');
```

```
%For ANum = Number of Cells A with which any given Cell B may exchange
ANum=SDANum.*randn(BNo+50,1)+MidANum;
%Rounding values
ANum=round(ANum);
%To remove values beyond the minimum and maximum,
%values are tested for excursion beyond extremes and replaced†
%with random values between minima and maxima.†
%Values are rounded for this variable
a=(1:BNo+50);
for a=(1:BNo+50)
    if ANum(a,1)<MinANum
        b=MinANum+rand*(MaxANum-MinANum);
        ANum(a,1)=round(b);
    end
    if ANum(a,1)>MaxANum
        b=MinANum+rand*(MaxANum-MinANum);
        ANum(a,1)=round(b);
    end
end
%Drawing a histogram
subplot(3,5,1);
histogram(ANum);
title ('Cells A / Cell B');

%For ExNum = Number of Exchanges within any given cell pairing
ExNum=SDExNum.*randn(BNo+50,1)+MidExNum;
%Rounding values
ExNum=round(ExNum);
%To remove values beyond the minimum and maximum,
%values are tested for excursion beyond extremes and replaced†
%with random values between minima and maxima.
%Values are rounded for this variable
a=(1:BNo+50);
for a=(1:BNo+50)
    if ExNum(a,1)<MinExNum
        b=MinExNum+rand*(MaxExNum-MinExNum);
        ExNum(a,1)=round(b);
    end
    if ExNum(a,1)>MaxExNum
        b=MinExNum+rand*(MaxExNum-MinExNum);
        ExNum(a,1)=round(b);
    end
end
%Drawing a histogram
subplot(3,5,2);
histogram(ExNum);
title ('Exchange Number');


%For Rate = Rate of cell-projection retraction for any given exchange event
Rate=SDRate.*randn(BNo+50,1)+MidRate;
%To remove values beyond the minimum and maximum,
%values are tested for excursion beyond extremes and replaced†
%with random values between minima and maxima.
a=(1:BNo+50);
for a=(1:BNo+50)
    if Rate(a,1)<MinRate
        Rate(a,1)=MinRate+rand*(MaxRate-MinRate);
    end
    if Rate(a,1)>MaxRate
        Rate(a,1)=MinRate+rand*(MaxRate-MinRate);
    end
end
%Drawing a histogram
subplot(3,5,4);
histogram(Rate);
title ('Rate');

%For Lgh = Length of cell-projection in any given exchange event
Lgh=SDLgh.*randn(BNo+50,1)+MidLgh;
%To remove values beyond the minimum and maximum,
%values are tested for excursion beyond extremes and replaced†
```

```
%with random values between minima and maxima.†
a=(1:BNo+50);
for a=(1:BNo+50)
    if Lgh(a,1)<MinLgh
        Lgh(a,1)=MinLgh+rand*(MaxLgh-MinLgh);
    end
    if Lgh(a,1)>MaxLgh
        Lgh(a,1)=MinLgh+rand*(MaxLgh-MinLgh);
    end
end
%Drawing a histogram
subplot(3,5,5);
histogram(Lgh);
title ('CellP Length');

%For Radi = Radius of cell-projection in any given exchange event
Radi=SDRadi.*randn(BNo+50,1)+MidRadi;
%To remove values beyond the minimum and maximum,
%values are tested for excursion beyond extremes and replaced†
%with random values between minima and maxima.
a=(1:BNo+50);
for a=(1:BNo+50)
    if Radi(a,1)<MinRadi
        Radi(a,1)=MinRadi+rand*(MaxRadi-MinRadi);
    end
    if Radi(a,1)>MaxRadi
        Radi(a,1)=MinRadi+rand*(MaxRadi-MinRadi);
    end
end
%Drawing a histogram
subplot(3,5,6);
histogram(Radi);
title ('CellP Radius');

%For Visc = Viscosity in any given exchange event
Visc=SDVisc.*randn(BNo+50,1)+MidVisc;
%To remove values beyond the minimum and maximum,
%values are tested for excursion beyond extremes and replaced†
%with random values between minima and maxima.
a=(1:BNo+50);
for a=(1:BNo+50)
    if Visc(a,1)<MinVisc
        Visc(a,1)=MinVisc+rand*(MaxVisc-MinVisc);
    end
    if Visc(a,1)>MaxVisc
        Visc(a,1)=MinVisc+rand*(MaxVisc-MinVisc);
    end
end
%Drawing a histogram
subplot(3,5,7);
histogram(Visc);
title ('Viscosity');

%Clearing values no longer needed
clear MaxANum MinANum MidANum MaxExNum MinExNum MidExNum
clear MaxRate MinRate MidRate MaxLgh MinLgh MidLgh
clear MaxRadi MinRadi MidRadi MaxVisc MinVisc MidVisc
clear a b SD SDANum SDExNum SDRate SDLgh SDRadi SDVisc


%-------------------------------------------------------------------
%-------------------------------------------------------------------
%CREATING MATRICES TO RECEIVE RESULTS

%Interim results of calculations are stored in IntRes where:
%Rows = BNo; Columns = 2*(maxExNum*maxANum)
%Each row contains results for one Cell B
%Odd numbered columns contain Flow Volumes of exchange
%Even numbered columns contain Fuorescene Values of exchange

maxANum = max(ANum);
maxExNum = max(ExNum);
ColumnNum = 2 * maxANum * maxExNum;
```

```
IntRes=zeros(BNo,ColumnNum);

%Results for individual event: Times Maximum Pressure (MxPresRec),
%flow volumes (VolRec) are in matrices
Times=zeros(BNo,ColumnNum/2);
MxPresBRec=zeros(BNo,ColumnNum/2);
MxPresARec=zeros(BNo,ColumnNum/2);
VolRec=zeros(BNo,ColumnNum/2);

%..............................
%CALCULATING FLOW AND FLUORESCENCE TRANSFER BETWEEN CELLS A AND B
%Where:
% Fluorescence and stiffness values of Cell A remains the same
%for all exchange events within the pairing with Cell B.
% Cell B can interact with more than one cell A.
% When Cell B starts to interact with a new Cell A, it shifts one step
%down the column of Cells A for stiffness, fluorescence and area values.

% A = Median stiffness of Cell A
% B = Median stiffness of Cell B
% BAr = Area of Cell B
% c = Number of Cells A that Cell B interacts with
% Ca = Cross sectional area of tube = pi*r^2
% F = Fluorescence value of Cell A
% i = Row number for each Cell B
% l = Length of cell-projection at time 0
% la = Length from origin to open end = l-lb
% lb = Length from origin to open end =((Z*(A-B))/(Ca*U*R))+(l/2)
% PAMax = Delta-Pressure (origin to open end A), time 0 when A<B
% PBMax = Delta-Pressure (origin to open end B), time 0
%† † Note: PAMax=(Ca*U*R*la^2)/(2*la);PBMax=(Ca*U*R*lb^2)/(2*lb)
% r = Radius of cell-projection
% R = Resistance per unit length by R=(8*v)/(pi*r^4)
% tc = Time at which first part of integration ceases
% tmax = Maximum time of retraction of cell-projection = U/l
% U = Constant rate of cell-projection retraction
% v = Viscosity of cytoplasm
% VB = Volume transferred to cell B
% x = Number of exchange events Cell A has with Cell B
% Z = Stiffness Correction Factor

for i=1:BNo
    c=datasample(ANum,1);
    for e=1:c
        x=datasample(ExNum,1);
        for n=1:x
            F=AFluor(i+e-1,1);
            A=AStiff(i+e-1,1);
            B=BStiff(i,1);
            r=datasample(Radi,1);
            l=datasample(Lgh,1);
            v=datasample(Visc,1);
            U=datasample(Rate,1);

            %Determining Ca by pi*r^2, and tmax by l/U
            Ca=pi*r^2;
            tmax=l/U;

            %Time for the transfer event between Cell A and Cell B
%An upper bound is tmax. Minimum possible time MinTim = tmax*PropMin;
%MaxTim=PropMax*tmax
a=rand;
t=(tmax*PropMin)+(a*((tmax*PropMax)-(tmax*PropMin)));


        %Determining R PBMax and VB if both cells have the same stiffness
        if A==B
            R=(8*v)/(pi*r^4);
            lb=((Z*(A-B))/(Ca*U*R))+(l/2);
            PBMax=(Ca*U*R*lb^2)/(2*lb);
            la=l-lb;
            PAMax=(Ca*U*R*la^2)/(2*la);
            VB=Ca*U*t*0.5;
```

```
        end

        %Determining R and tc as required if cells are of unequal stiffness
        if A~=B
            R=(8*v)/(pi*r^4);
            a=abs((2*Z*(A-B))/(Ca*R*U^2));
            tc=tmax-a;
            if tc<0;
                tc=0;
            end
        end

        %Determining PAMax PBMax and VB if Cell A has higher stiffness
        %than Cell B
        if A>B
            lb=((Z*(A-B))/(Ca*U*R))+(l/2);
            PBMax=(Ca*U*R*lb^2)/(2*lb);
            la=l-lb;
            PAMax=(Ca*U*R*la^2)/(2*la);
            if t<=tc
                VB=((Ca*U*t)/2)-((Z*(A-B))/(R*U))*(log((l-(U*t))/l));
            end
            if t>tc
      VB=((Ca*U*tc)/2)-((Z*(A-B))/(R*U))*(log((l-(U*tc))/l))+(Ca*U*(t-tc));
            end
        end

        %Determining PAMax, PBMax & VB if Cell A has lower stiffness
        %than Cell B
        if A<B
            lb=((Z*(A-B))/(Ca*U*R))+(l/2);
            PBMax=(Ca*U*R*lb^2)/(2*lb);
            la=l-lb;
            PAMax=(Ca*U*R*la^2)/(2*la);
            if t<=tc
                VB=((Ca*U*t)/2)-((Z*(A-B))/(R*U))*(log((l-(U*t))/l));
            end
            if t>tc
                VB=((Ca*U*tc)/2)-((Z*(A-B))/(R*U))*(log((l-(U*tc))/l));
            end
            end
            Fl=(VB*F)/(BAr*AHght);
            IntRes(i,((2*maxExNum)*(e-1))+(2*n)-1)=VB;
            IntRes(i,((2*maxExNum)*(e-1))+(2*n))=Fl;
            Times(i,((maxExNum)*(e-1))+(n))=t;
            MxPresBRec(i,((maxExNum)*(e-1))+(n))=PBMax;
            MxPresARec(i,((maxExNum)*(e-1))+(n))=PAMax;
            VolRec(i,((maxExNum)*(e-1))+(n))=VB;
        end
    end
end

%Clearing values no longer needed
clear a A B Ca l le r t tc tmax U VB1a VB1b VB2a VB2b
clear la lb Fl VB PAMax PBMax AHght BAr n F Z

%.............................†
%MAKING CONVENIENT SINGLE COLUMN LISTS FOR INDIVIDUAL Times VB, PBMax,
%PAMax
TimList=Times(:);
VolList=VolRec(:);
PBMaxList=MxPresBRec(:);
PAMaxList=MxPresARec(:);

%Removing non-zero values from TimList VolList, PBMaxList and PAMaxList by:
%Calculating number of elements in source matrices; creating matrices
%of the correct size to receive non-zero results (Suffix-NZ); loading†
%non-zero results
n=ColumnNum*BNo/2;

%Creating TimNZ
z=0;
```

```
for a=(1:n)
    if TimList(a,1)>0
        z=z+1;
    end
end

TimNZ=zeros(z,1);

z=0;
for a=(1:n)
    if TimList(a,1)>0
        z=z+1;
        TimNZ(z,1)=TimList(a,1);
    end
end

%Creating VolListNZ
z=0;
for a=(1:n)
    if VolList(a,1)>0
        z=z+1;
    end
end

VolListNZ=zeros(z,1);

z=0;
for a=(1:n)
    if VolList(a,1)>0
        z=z+1;
        VolListNZ(z,1)=VolList(a,1);
    end
end

%Creating PAMaxListNZ
z=0;
for a=(1:n)
    if PAMaxList(a,1)>0
        z=z+1;
    end
end

PAMaxListNZ=zeros(z,1);

z=0;
for a=(1:n)
    if PAMaxList(a,1)>0
        z=z+1;
        PAMaxListNZ(z,1)=PAMaxList(a,1);
    end
end

%Creating PBMaxListNZ
z=0;
for a=(1:n)
    if PBMaxList(a,1)>0
        z=z+1;
    end
end

PBMaxListNZ=zeros(z,1);

z=0;
for a=(1:n)
    if PBMaxList(a,1)>0
        z=z+1;
        PBMaxListNZ(z,1)=PBMaxList(a,1);
    end
end

%Clearing values no longer needed
clear ANum AStiff c ColumnNum e exNum Fl i Rate MedS Times TimList
clear InPres l Lgh n Radi t Tim v x a z PropMax PropMin
```

```
clear ExNum VolRec MxPresBRec MxPresARec Visc R

%Plotting Graphs of TimeNZ, VolListNZ, PBMaxListNZ and PAMaxListNZ
% NOTE THAT: It is convenient to set the maximum xlim for PAMax and
%PBMax at 10.  This is possible in the current simulations because
%the maximum values of P reached are below 10Pa. In some simulations,
%however, xlim may need to be increased to properly graph these
%results
subplot(3,5,3);
histogram(TimNZ);
title ('Times');

subplot(3,5,8);
histogram(VolListNZ);
title ('Ind Vol');

subplot(3,5,9);
histogram(PBMaxListNZ);
%Note: the range for xlim can be increased by changing the second term
xlim([0 10]);
title ('PB Max');

subplot(3,5,10);
histogram(PAMaxListNZ);
%Note: the range for xlim can be increased by changing the second term
xlim([0 10]);
title ('PA Max');


%-------------------------------------------------------------------
%-------------------------------------------------------------------
%SUMMATING RESULTS

%Creating matrices to receive total Flow Volume and Fluorecence Values
TotalFlow=zeros(BNo,1);
TotalFluor=zeros(BNo,1);

%Summating Total Flow Volume and Total Fluorescence Exchange per Cell B
ColNum=maxExNum*maxANum;
Temp=zeros(1,ColNum);
for i=1:BNo
    for c=1:ColNum
        Temp(1,c)=IntRes(i,(2*c)-1);
        TotalFlow(i,1)=sum(Temp);
    end
end

%.............................†
%GRAPHING TOTAL FLOW RESULT
subplot(3,5,11);
histogram(TotalFlow);
title ('TotalVolume');

for i=1:BNo
    for c=1:ColNum
        Temp(1,c)=IntRes(i,(2*c));
        TotalFluor(i,1)=sum(Temp);
    end
end

%.............................†
%GRAPHING TOTAL FLUORESCENCE RESULT
subplot(3,5,12);
histogram(TotalFluor);
title ('Fluorescence');

%Clearing values no longer needed
clear AFluor BStiff maxANum Temp c ColNum d i ExNum
clear maxExNum VolListNZ VolList PBMaxList PAMaxList


%.............................
%CREATING TotFlowListNZ AND TotFluorListNZ FOR LATTER USE IN RESULTS TABLE
```

```
%Creating TotFlowListNZ
z=0;
for a=(1:BNo)
    if TotalFlow(a,1)>0
        z=z+1;
    end
end

TotFlowListNZ=zeros(z,1);

z=0;
for a=(1:BNo)
    if TotalFlow(a,1)>0
        z=z+1;
        TotFlowListNZ(z,1)=TotalFlow(a,1);
    end
end


%Creating TotFluorListNZ
z=0;
for a=(1:BNo)
    if TotalFluor(a,1)>0
        z=z+1;
    end
end

TotFluorListNZ=zeros(z,1);

z=0;
for a=(1:BNo)
    if TotalFluor(a,1)>0
        z=z+1;
        TotFluorListNZ(z,1)=TotalFluor(a,1);
    end
end



%---------------------------------------------------------------------
%---------------------------------------------------------------------
%GRAPH COMPARING TOTAL FLUORESCENCE RESULT WITH EXPERIMENTAL DATA

%This permits rapid visual comparison of the results of an experimental
%data set with that of the calculated fluorescence result.

%Establishing the number of values less than 6 for TotalFluor.
%Creating a matrix WeedFluor to accept these, and transferring
%Values from TotalFluor into WeedFluor accordingly.

c=0;
for i=1:BNo;
    if TotalFluor(i,1)<=6;
        c=c+1;
    end
end

d=0;
WeedFluor=zeros(c,1);
for i=1:BNo;
    if TotalFluor(i,1)<=6;
        d=d+1;
        WeedFluor(d,1)=TotalFluor(i,1);
    end
end

%Establishing the number of values less than 6 for ExpFluor.
%Creating a matrix WeedExp to accept these, and transferring
%Values from ExpFluor into WeedExp accordingly.
%NOTE: Changing the value '6' below permits the range to be changed.†
%'6' appears in the current script because it was convenient in the†
%particular simulations performed.†
```

```
c=0;
for i=1:BNo;
    if ExpFluor(i,1)<=6;
        c=c+1;
    end
end

d=0;
WeedExp=zeros(c,1);
for i=1:BNo;
    if ExpFluor(i,1)<=6;
        d=d+1;
        WeedExp(d,1)=ExpFluor(i,1);
    end
end

%Plotting results on the same histogram
subplot (3,5,13);
edges =-0.15:0.3:6;
hFluor=histcounts(WeedFluor,edges);
exFluor=histcounts(WeedExp,edges);
bar(edges(1:end-1),[hFluor;exFluor]');
xlim([-0.2 6]);
title ('Exper Vs Sim');

%Clearing values no longer needed
clear BNo c d edges exFluor hFluor i WeedExp  WeedFluor IntRes


%----------------------------------------------------------------------
%----------------------------------------------------------------------
%PREPARATION OF A TALBE SUMMARISING KEY RESULTS

TempRes=zeros(3,5);

%Loading elements into TempRes
TempRes(1,1) = median(PBMaxListNZ);
TempRes(2,1) = min(PBMaxListNZ);
TempRes(3,1) = max(PBMaxListNZ);

TempRes(1,2) = median(PAMaxListNZ);
TempRes(2,2) = min(PAMaxListNZ);
TempRes(3,2) = max(PAMaxListNZ);

TempRes(1,3)= median(TimNZ);
TempRes(2,3) = min(TimNZ);
TempRes(3,3) = max(TimNZ);

TempRes(1,4) = median(TotFlowListNZ);
TempRes(2,4) = min(TotFlowListNZ);
TempRes(3,4) = max(TotFlowListNZ);

TempRes(1,5) = median(TotFluorListNZ);
TempRes(2,5) = min(TotFluorListNZ);
TempRes(3,5) = max(TotFluorListNZ);

% Creating a table ResultsTable
varNames = {'PBMax','PAMax','Time','Flow','Fluor'};
rowNames = {'Median', 'Minimum', 'Maximum'};
ResultsTable = array2table(TempRes,'VariableNames',varNames,...
    'RowNames',rowNames);

%Clearing values no longer needed
clear rowNames TempRes varNames z a

%END SCRIPT
```
